# Major QTLs for seedling traits in barley using a DArT-based linkage map

**DOI:** 10.1101/2019.12.28.889899

**Authors:** Ludovic Capo-chich, Sharla Eldridge, Ammar Elakhdar, Toshihiro Kumamaru, Anthony O. Anyia

## Abstract

Seed vigor is considered as the most critical stage for barley production, and cultivar with high early seedling vigour (ESV) allow plants to form a canopy more quickly. In this study, the QTLs of seedling vigour related-traits were investigated using 185 RILs derived from Xena and H94061120 using DArT approach. In total, 46 significant QTLs for ESV related-traits were detected. The total map length was 1075.1 cM with an average adjacent-marker distance of 3.28 cM. Fourteen QTLs for BY were found on all chromosomes, two of them co-located with QTLs on 1H for GY. The related-traits; LL1, LL2, LA1 and LDW1 had high heritability (>60%). Meanwhile, a significant correlation was observed between GY and BY, which provide the clear image of these traits in the selection process. Our results demonstrate that a pleiotropic QTL related to SLA2, BY, and GY was linked to to the DArT markers bPb-9280 and bPb-9108 on 1H, which could be used to significantly improve the seed vigor by marker-assisted selection and possible future map-based cloning of the gene of intrest.

## 1. Introduction

Barley (*Hordeum vulgare L*.) is the fourth most important cereal crop in the world and one of the earliest domesticated crops used for the human food and animal feed, and therefore a great economic importance. Indeed, it has become an important ideal model for molecular genetics and functional studies particularly for monocot crops because it is early maturing, diploid, self-fertilizing and has a short growth period, owing high-quality genome sequences, and rich in germplasm resources (Heneen, 2011; Qin et al., 2015).

However, the seedling stage is considered as the most critical stage for barley production, and contribute to barley growth and development, including water and nutrient uptake, biotic and abiotic stresse resistance, and can influence biomass, grain yield, and grain quality (Ullmannová et al., 2013; Wang et al., 2017). Moreover, rapid early development of leaf area and above-ground biomass, referred to as early seedling vigour (ESV), is recognized as desirable to improve yield under water-limited environments (Ullmannová et al., 2013; Singh et al., 2017; Wen et al., 2018). ESV determines the potential for rapid and uniform emergence of plants under a wide range of field conditions (Rajjou et al., 2012). It is mainly expressed as increased seedling weight or height, which usually neglects germination speed (Lu et al., 2007). Differences in ESV among cereals have been associated with variation in a number of related traits such as specific leaf area (SLA), leaf area, specific leaf weight (SLW), and the rate of seedling emergence (Heneen, 2011; Li et al., 2014; Mahender et al., 2015). However, the impact of these parameters on plant performance depends on environmental conditions.

A number of studies e.g. (López-Castañeda et al., 1996; Richards et al., 2002; Sundgren et al., 2018) have reported genetic increases in early vigour to be associated with greater biomass and grain yield for crops grown in Mediterranean environments. Phenotypic differences in early vigour among temperate and tropical cereals have been associated with variation in rate of seedling emergence, embryo size and leaf area-to-leaf weight ratio, or specific leaf area (SLA) of seedling leaves (Richards & Lukacs, 2002). As reported by (Richards, 2000), the leaf area can increase without incurring additional costs by decreasing the amount of photosynthetic machinery per unit leaf area. This in turn increases the leaf area per unit weight, measured as the specific leaf area (SLA, leaf area per unit dry mass). SLA was suggested to be suitable for selecting plants with superior early vigour. A high SLA allows the plant to close the leaf canopy at minimal carbon expenses. Thus, when rapid canopy closure improves the water use efficiency, as suggested for small-grain cereals (Richards et al., 2002), maximization of leaf area by increasing the SLA may be important in reducing evaporation from the soil surface (Richards et al., 2002). Rapid leaf area development early in the season has potential to increase water use efficiency and grain yield of winter cereals (Rebetzke et al., 2004). The early vigor is an important trait for waterlogging tolerance. Rapid seedling establishment and a narrow root stele promotes waterlogging tolerance in spring wheat (Sundgren et al., 2018).

Early seed vigor is a complex trait which is regulated by multi-genes affecting seedling leaf; width, length, area, weight and specific leaf area. These characteristics have been widely reported to provide improved productivity of cereal crops in a wide range environments (Botwright et al., 2002).

Advances in biotechnology and in the measurement of physiological traits enable plant breeders to utilize the technologies in their breeding programs. For instance, the development of molecular techniques such as the DArT technology has enabled quantitative trait loci (QTL) for morphological and physiological traits to be identified in plant species (Argyris et al., 2005). QTL analysis has become a powerful tool to dissect complex traits and identify chromosomal regions harboring genes that control these quantitative traits (Tyrka et al., 2011), and has been used extensively to improve the important traits in barley (Liu et al., 2015; Wang et al., 2016). A large number of QTLs associated with seedling seed vigor were reported using biparental segregating populations in many crops (Rajjou et al., 2012; Mansour et al., 2014). The QTLs for yield (Mikołajczak et al., 2016) traits at later growth stages have been characterized (Zhou et al., 2016), and the QTLs for seedling characteristics described were primarily recognized under salt tolerance (Sbei et al., 2014) (Elakhdar et al., 2016), water logging (Broughton et al., 2015), drought tolerance (Chen et al., 2010) and nitrogen stress tolerance (Hoffmann et al., 2012) while, seedling characteristics related to grain yield and biomass at early seed developmental stages were not well investigated.

The goals of the present study are; to determine the extent of genetic variation in early seedling vigour and their effects on biomass and grain yield under the rainfed condition, to assess QTLs of seedling vigour related-traits, and to identify molecular markers linked to seedling vigour related-traits using a recombinant inbreed line (RIL) population.

## 2. Materials and Methods

### 2.1. Plant material

A mapping population of 185 F_5_ recombinant inbred lines (RILs), developed by single-seed descent (SSD) to generation F_2_, was used to construct DArT-based linkage map. The RIL population was generated from cross of ‘Xena’ and ‘H94061120’ barley varieties. The chosen varieties, obtained from the Field Crop Development Centre (FCDC), Lacombe, Alberta, Canada), express a wide range of variation on agronomic traits.

### 2.2. Field experiment conditions

Phenotype evaluation of the 185 RILs was conducted at Vegreville, Alberta, Canada (53°34’N, 113°31’W, 639.3 m above sea level altitude) during two years. The soil texture was Malmo series of an Eluviated Black Chernozemic, under rainfed conditions. According to AgroClimatic Information Service (ACIS) (http://agriculture.alberta.ca/acis/), the average annual precipitation and within-season rainfall (June–August; 2009) was 382 ± 62 and 193 ± 52 mm. A validation experiment was arranged in RCBD with three replications.

### 2.3. Evaluation of phenotypic triats

The expermintal material was evaluated in CRD design with three replications. Twelev quantitative triats including; early seedling, Leaf length (LL, cm) width (LW, cm), Leaf dry weight (LDW, mg), Leaf area (LA, cm^2^) and Specific leaf area (SLA) were messured. Early seedling vigour was assessed at the third leaf stage when the coleoptile tiller was absent. Leaf length width were measured on the first and second leaves. Leaf dry weight was determined after oven-drying at 70°C for 48 hours. Leaf area was measured according to the method descriped (Rebetzke & Richards, 1999). Specific leaf area (SLA) was determined as the ratio of leaf area to dry weight of the first two leaves. The mean squares due to treatment for the studied quantitative ctraits are compared in the RILs along with the two parental lines using analysis of variance.

The following parameters were compared in the RILs along with the two parental lines using analysis of variance: leaf width of the first leaf (LW1), leaf width of the second leaf (LW2), leaf length of the first leaf (LL1), leaf length of the second leaf (LL2), leaf area of the first leaf (LA1), leaf area of the second leaf (LA2), leaf dry weight of the first leaf (LDW1), leaf dry weight of the second leaf (LDW2), specific leaf area of the first leaf (SLA1), specific leaf area of the second leaf (SLA2), seed yield (Syield), and total dry biomass (Bio).

### 2.4. Estimation of genotypic parameters

The recorded data were then used for statistical analysis. Mean values were used for the analysis. Different genetic parameters like phenotypic variance and genotypic variance were computed based on the method suggested by (Burton & Devane, 1953). The phenotypic and genotypic coefficient o f variation (PCV and GCV) were computed with the formula suggested by (Burton, 1952). The Haritability in the broad sense and genetic advance was estimated by adopting the method suggested by (Johnson et al., 1955).

#### Genotypic and phenotypic of variance

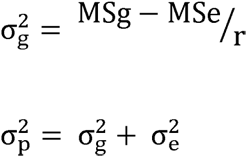

where, MSg, MSp, Mse; mean square due to genotypes, phenotypes and error, respectively. r; number of replication. 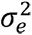; environmental variance

#### Genotypic coefficient and phenotypic Coefficient of variance

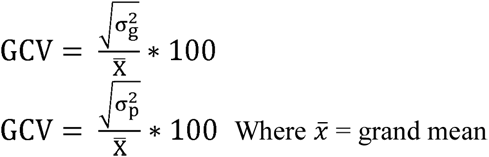

#### Heritability in broad sense

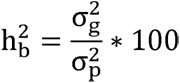

#### Genetic advance

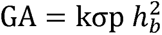

Where k is the selection differential at 5% selection intensity, the value of which is 2.06 *σ_b_* is the phenotypic standard deviation = 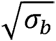

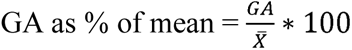

#### Genotyping and Construction of genetic linkage map

Total genomic DNA was extracted from 5 week-old leaves of the 185 RILs along with the parental lines using a protocol developed by Diversity Array Technology (http://www.diversityarrays.com/sites/default/files/resources/DArT_DNA_isolation.pdf.). The extracted DNA was sent to DArT Pty Ltd for genotyping according to the standard barley DArT® array by Triticarte Pty Ltd., Canberra (http://www.triticarte.com.au/). A quality parameter Q, which is the variance of the hybridization intensity between allelic states as a percentage of the total variance, was calculated for each marker. Only markers with a Q and call rate greater than 80% were selected for linkage analysis. Polymorphic loci were selected after discarding those with a minor allele frequency of 0.5, a missing value of more than 20 %, or a common position. The linkage analysis was conducted using JoinMap 4.0 with a recombination frequency of 0.25, and all markers were grouped among the seven chromosomes. Haldane’s map function was used to calculate the recombination rate of the genetic distance in CentiMorgan (cM).

### 2.5. QTL analysis

For QTL mapping, the linkage map constructed with marker data from 185 F_5_ RILs derived cross (Xena × H94061120) was used. To estimate the association of each marker to a trait, composite interval mapping (CIM) was performed using WinQTL Cartographer v2.5 (Wang et al., 2011). Logarithm of odds (LOD) threshold score for QTL (*P* = 0.05) was determined using a 1000-permutation test by shuffling the phenotypes means with the genotypes. A LOD score of 2 indicates that the model containing the estimated QTL effect is 100 times more likely than is the model with no QTL effect. A LOD threshold score of >2.5 at 1000 permutations was considered significant to identify and map the QTLs in the barley population. The 95% confidence intervals of the QTL locations were determined by one-LOD intervals surrounding the QTL peak (Mangin et al., 1994; Rajjou et al., 2008). Composite interval mapping is based on the idea that the residual error term in a QTL analysis is the within genotypic class variance. This residual variance is partly due to experimental error but may also be due to variation caused by segregation of other QTLs outside of the region being tested. To reduce the background genetic segregation variance when conducting interval mapping, CIM first uses regression analysis to choose a subset of markers that have the biggest effects. These are used as “cofactors” in a subsequent interval mapping. When testing positions near a cofactor, that particular cofactor is dropped from the model, so that the QTL effects in that region can be more precisely identified.

## 3. Results

### 3.1. Phenotypic evaluation and correlation of seedling vigour-related traits

The parental lines Xena and H94061120 showed a highly significant differences in the leaf length of the first leaf stage (LL1) (P < 0.05) in each of the two years. Meanwhile, a higher leaf area of the first leaf LA1; 9.31 cm^2^ and the second leaf LA2; 1098 cm^2^, leaf dry weight of the first leaf (mg) LDW1; 0.126 mg, and leaf dry weight of the second leaf (mg) LDW2; 0.15 mg values for Xena than the ones for H94061120 were obtained, although not significantly different (Table 1). The trait phenotypic variance among RILs along with the parental lines showed highly significant differences in seedling vigour-related traits. SLA, LA, LW, LL of the first and second leaves displayed continuous frequency distribution (Figure 1). Nine of the 12 traits considered (Table 1), showed symmetrical distributions, two moderately were skewed distribution and one displayed binomial distribution. Grain yield and biomass were slightly skewed to higher values (Figure 1). There were a significant differences among the RILs for all the traits and means were assumed to be normally distributed. Considerable transgressive segregation was evident for all early vigour traits in this population.

**Figure 1.**
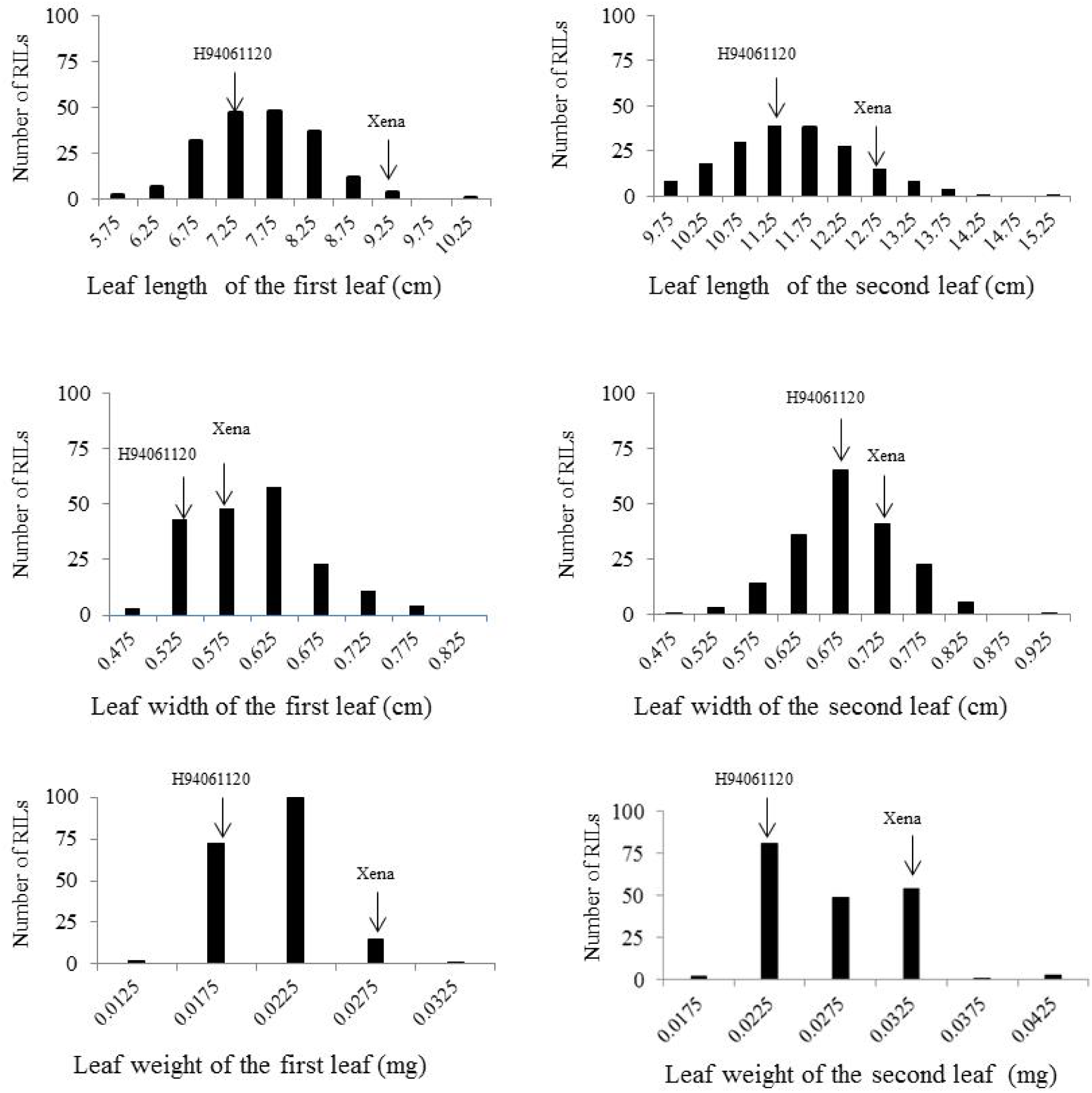
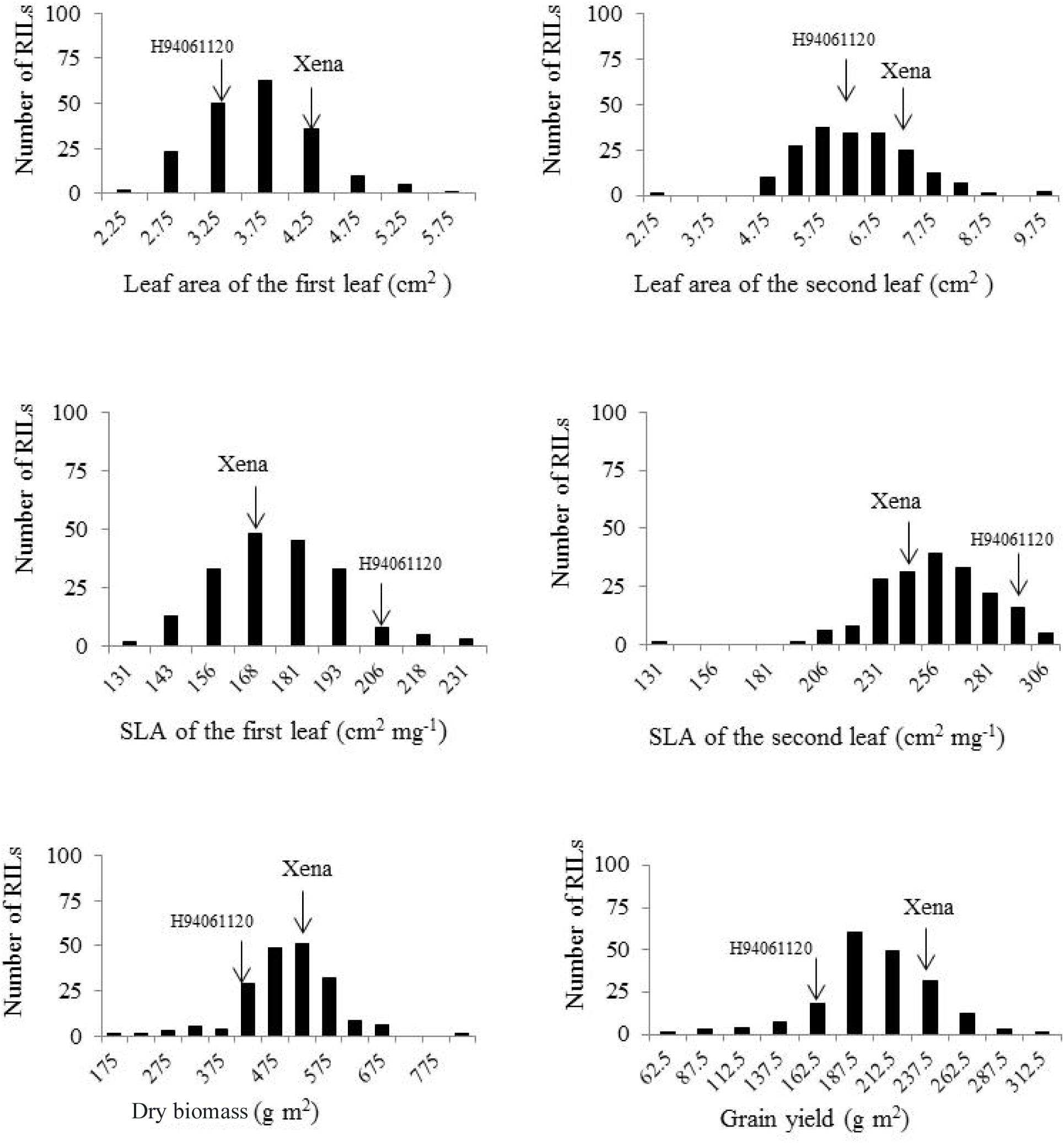
Frequency distribution of the first and second leaf length, leaf width, leaf area, specific leaf area, leaf weight and biomass and grain yield measured on the recombinant inbred lines (RILs) derived from a cross between two-row barley H94061120 (high NSC) × Xena (feed). Parental means are indicated for each trait.

**Table 1.** Means for the parental lines in the parental comparison study for the following traits.

| Traits | Xena | H94061120 | LSD (0.05) |
| --- | --- | --- | --- |
| LL1 | 9.31a† | 6.68b | 1.69 |
| LL2 | 12.88a | 10.43a | 3.35 |
| LW1 | 0.59a | 0.52a | 0.2 |
| LW2 | 0.75a | 0.59a | 0.81 |
| LA1 | 4.61a | 2.83b | 1.52 |
| LA2 | 1098a | 7.68b | 9.32 |
| LDW1 | 0.126a | 0.073b | 0.01 |
| LDW2 | 0.150a | 0.093b | 0.06 |
| SLA1 | 176.17a | 190.99a | 86.96 |
| SLA2 | 263.25a | 376.94a | 283.61 |
| BY | 561.33a | 523.23a | 275.91 |
| GW | 234.23a | 212.33a | 182.78 |

LL1; leaf length of the first leaf (cm), LL2; leaf length of the second leaf(cm), LW1; leaf width of the first leaf (cm), LW2; leaf width of the second leaf (cm), LA1; leaf area of the first leaf (cm<sup>2</sup>), LA2; leaf area of the second leaf (cm<sup>2</sup>), LDW1; leaf dry weight of the first leaf (mg), LDW2; leaf dry weight of the second leaf (mg), SLA1; specific leaf area of the first leaf (cm<sup>2</sup> mg<sup>-1</sup>), SLA2; specific leaf area of the second leaf (cm<sup>2</sup> mg<sup>-1</sup>), BY; dry biomass (gm<sup>-2</sup>), GY; grain yield (gm<sup>-2</sup>).
†: Means within a row with the same letter are not significantly different according to the Least Significant Difference (LSD) at a 0.05 probability level

### 3.2. Variances, coefficients of variation, heritability and genetic advance for traits in the barley RILs

Phenotypic, genotypic and environmental variances as well as their coefficients of variation are presented in Table 2. The mean quares indicted that the significant differences present among the RILs for all the traits (P > 0.01) exept for LDW2 (Supplemental Table 2). A high genotypic and phenotypic variances were observied for biomass (BY gm^-2^); 4030.20 gm^-2^ / RIL and 1933.26 gm^-2^ / RIL, respectively (Table 2). The coefficients of variation (CV) was higth for LDW2. The highest genotypic (GCV) and phenotypic coefficient of variation (PCV) were observed for BY gm^-2^ and grain weight (GY gm^-2^), whereas the lowest GCV and PCV values were recorded for the trait of LDW1; 0.05 cm^2^ and 0.13 cm^2^, respectively. The broad sense heritability (h^2^) estimate varied from 0.01% in LDW2 to 84.65% in LL1. The studies traits LL1, LL2, LA1 and LDW1 had high heritability (>60%). The hightes value of genetic advance (GA) was observied with GY gm^-2^ / RIL (2038.06 gm^-2^ / RIL). The genetic advance as percentage of means (GA 5%) for twelve traits ranged from 0.55 in LDW1 to 1879.66 gm^-2^ / RIL for traits.

**Table 2.** Genetic parameters among twelve early seedling vigour-related traits of 185 RILs of barley

|  | LL1 | LL2 | LW1 | LW2 | LA1 | LA2 | LDW1 | LDW2 | SLA1 | SLA2 | BY | GY |
| --- | --- | --- | --- | --- | --- | --- | --- | --- | --- | --- | --- | --- |
| CV | 6.62 | 9.46 | 12.08 | 33.65 | 15.51 | 16.77 | 13.44 | 1617.03 | 13.68 | 13.23 | 22.04 | 25.50 |
| Min | 1.91 | 7.88 | 0.46 | 0.52 | 2.21 | 3.64 | 0.08 | 0.02 | 126.40 | 157.34 | 228.60 | 76.33 |
| Max | 3.42 | 14.37 | 0.81 | 2.27 | 6.08 | 9.90 | 0.19 | 70.63 | 242.17 | 310.76 | 705.70 | 296.13 |
| Mean | 2.52 | 11.45 | 0.60 | 0.69 | 3.67 | 6.28 | 0.12 | 0.70 | 175.56 | 254.28 | 499.03 | 200.59 |
| GV | 0.46 | 0.62 | 0.00 | 0.01 | 0.27 | 0.55 | 0.00 | 0.01 | 140.80 | 251.17 | 1933.26 | 420.48 |
| PV | 0.71 | 1.79 | 0.01 | 0.06 | 0.59 | 1.66 | 0.00 | 127.14 | 717.78 | 1383.15 | 14030.20 | 3037.15 |
| PCV | 3.13 | 5.22 | 0.41 | 2.92 | 5.40 | 8.80 | 0.13 | 6077.78 | 136.29 | 181.32 | 937.17 | 504.71 |
| GCV | 2.03 | 1.80 | 0.12 | 0.30 | 2.46 | 2.91 | 0.05 | 0.29 | 26.73 | 32.93 | 129.14 | 69.88 |
| $h^2_b$ % | 84.65 | 61.31 | 54.43 | 25.78 | 71.47 | 59.75 | 69.00 | 0.01 | 42.27 | 39.96 | 32.41 | 32.53 |
| GA 5% | 1.24 | 2.27 | 0.01 | 0.03 | 0.88 | 2.04 | 0.00 | 0.04 | 625.86 | 1140.33 | 9379.99 | 2038.06 |
| GA% of mean | 49.22 | 19.81 | 1.37 | 4.66 | 23.90 | 32.54 | 0.55 | 5.40 | 356.50 | 448.46 | 1879.66 | 1016.05 |

LL1; leaf length of the first leaf (cm), LL2; leaf length of the second leaf (cm), LW1; leaf width of the first leaf (cm), LW2; leaf width of the second leaf (cm), LA1; leaf area of the first leaf (cm<sup>2</sup>), LA2; leaf area of the second leaf (cm<sup>2</sup>), LDW1; leaf dry weight of the first leaf (mg), LDW2; leaf dry weight of the second leaf (mg), SLA1; specific leaf area of the first leaf (cm<sup>2</sup> mg<sup>-1</sup>), SLA2; specific leaf area of the second leaf (cm<sup>2</sup> mg<sup>-1</sup>), BY; dry biomass (gm<sup>-2</sup>), GY; grain yield (gm<sup>-2</sup>)
CV; coefficients of variation, GV; Genotypic variance, PV; Phenotypic variance, PCV; Phenotypic coefficient of variation, GCV; Genotypic coefficient of variation, $h^2_b$ %; heritability in broad sense, GA5%; Genetic advance 5%, GA of mean; Genetic advance % of mean

### 3.3. Phenotypic correlations

Correlations among traits are shown in Figure 2. The strongest correlation was found between LA and LW. There is a close relationship between the two traits of the first and second leaves. The positive correlation between leaf length of the second leaf (LA2) and LW of the first leaf (LA1) is similar to the one between LA1 and LW2. An identical pattern is found between the pairs (LA2 - LA1) and LW of the second leaf (LW22 - LW1). SLA is positively correlated with LA and negatively correlated with LDW, regardless of the rank of the leaves. However, an examination of the relationship between SLA and LDW of the second leaf revealed a significantly negative correlation, while SLA1 is weakly correlated with LDW1. The relationship between SLA1 and LA1 was similar to the one between SLA2 and LA2. An identical correlation pattern was found between LA and LL of the first and second leaves. A weak correlation was observed between LDW2 and LL2, while the correlation between LDW1 and LL1 was significant. SLA2, LA2, LL2, and LW2 were positively correlated with seed yield and biomass (Figure 2), while a weak correlation was found between the characteristics of the first leaf (SLA1, LA1, LL1, LW1) and GY, suggesting that the early seedling traits associated with the second leaf could be used to predict seed yield and biomass. A significant correlation was observed between GY and BY.

**Figure. 2.**
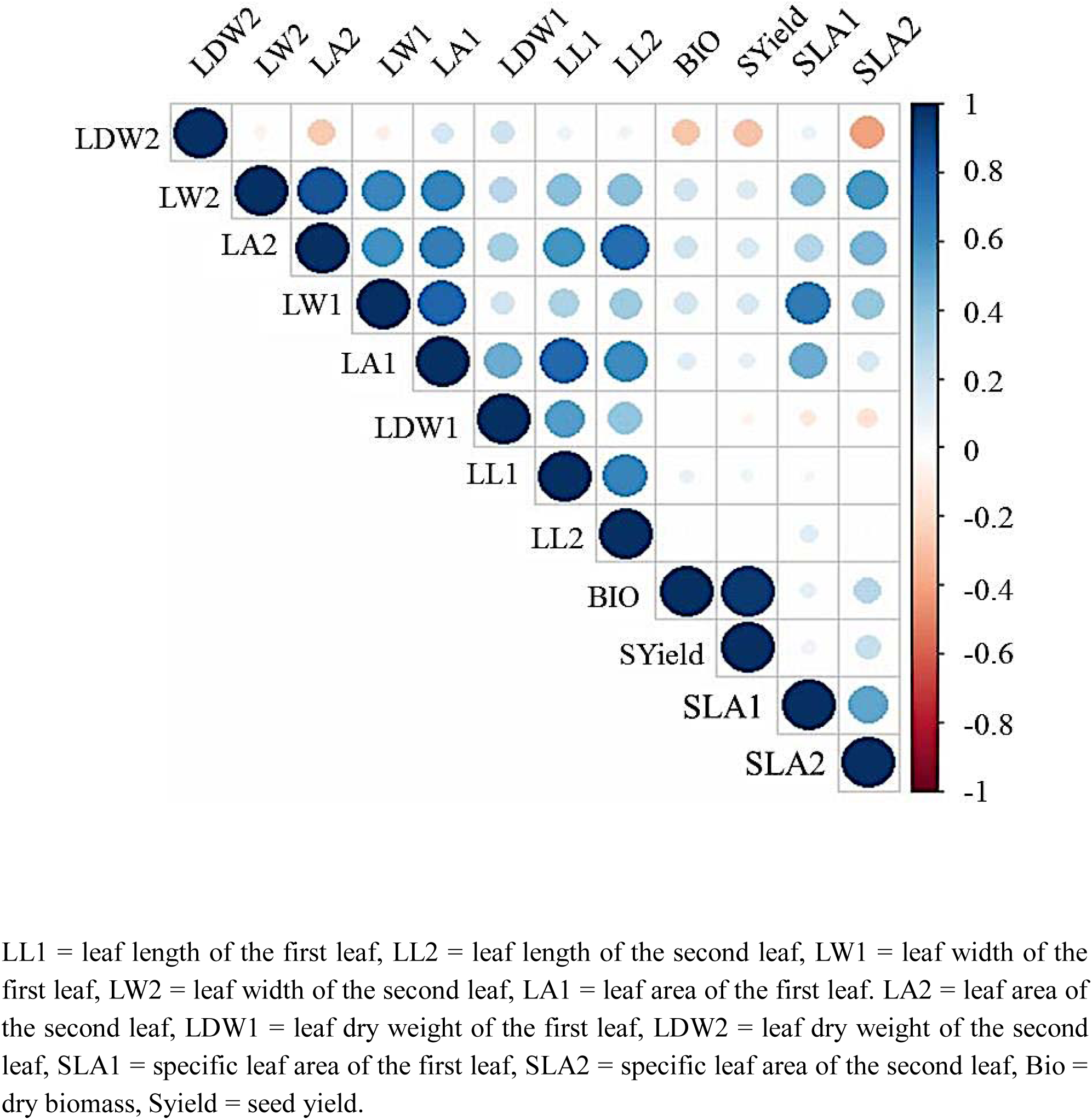
Visualize correlation matrix among variables using correlogram. Positive correlations are displayed in blue and negative correlations in red color. Color intensity and the size of the circle are proportional to the correlation coefficients. In the right side of the correlogram, the legend color shows the correlation coefficients and the corresponding colors.

The principal component analysis of these traits, shows a better insight into the relationships within and between traits. SLA2; GY; BY, LW1; LW2 and LA1; LA2 were closely correlated across the trials (Figure 3). All the points corresponding to each trait were placed in the same quadrant of the graph of the loadings on the first two principal components. These two components explained 60 % of the total variance. LDW1; LDW2 data points, however, were distributed over two quadrants, indicating changes in the direction of correlations within this trait and between traits.

**Figure. 3.**
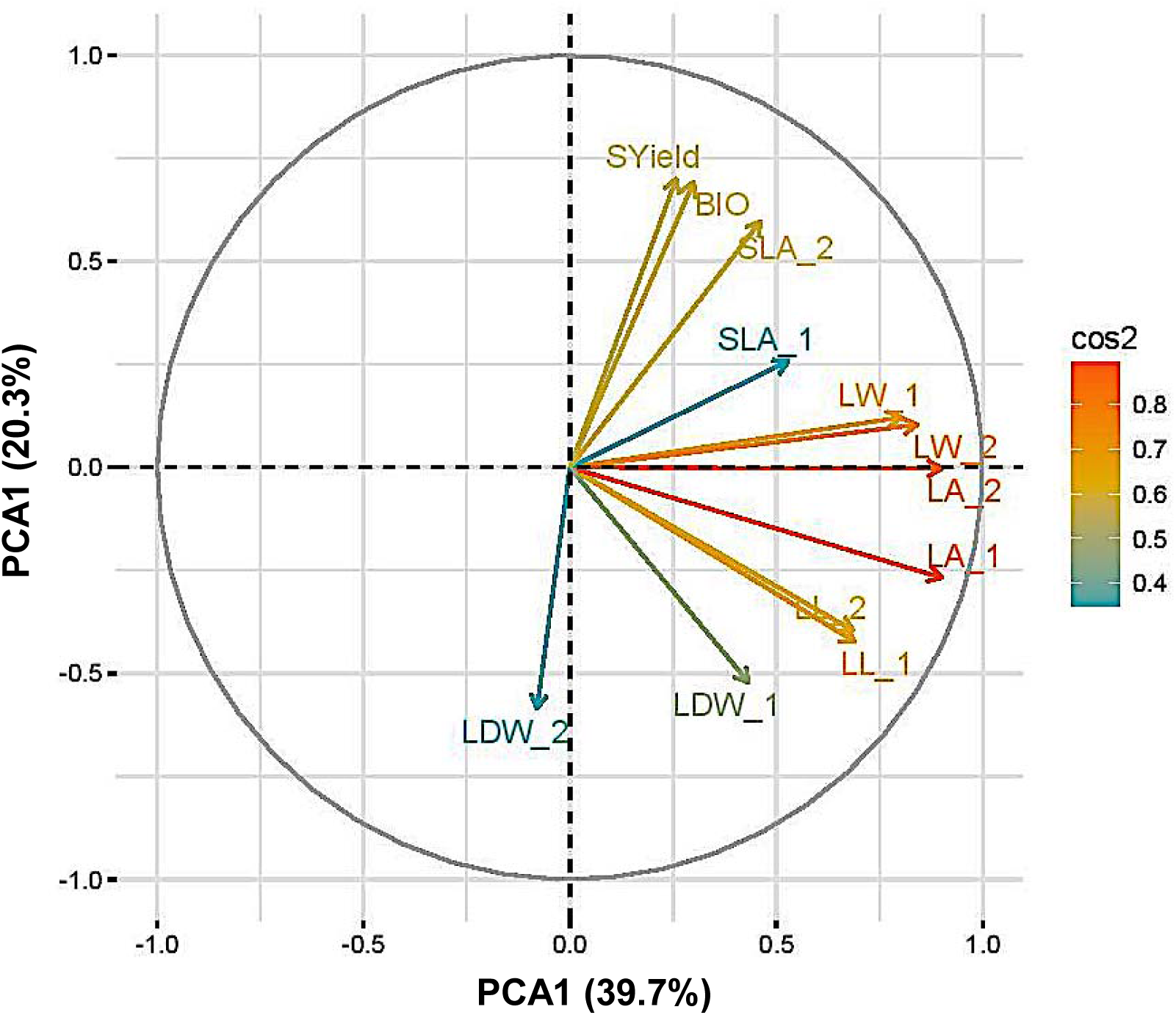
Plot of the first two axes of a principal component analysis carried out with the variables leaf width of the first leaf (LW1), leaf width of the second leaf (LW2), leaf length of the first leaf (LL1), leaf length of the second leaf (LL2), leaf area of the first leaf (LA1), leaf area of the second leaf (LA2), leaf dry weight of the first leaf (LDW1), leaf dry weight of the second leaf (LDW2), specific leaf area of the first leaf (SLA1), specific leaf area of the second leaf (SLA2), grain yield (GY), and total dry biomass (BY).

### 3.4. Construction of genetic map

A genetic map was constructed using the 328 polymorphic DArT markers on the seven barley chromosomes (Figure 4). The generated map spanned 1075.1 cM distance of the barley genome with an average marker density of 3.38 cM. Each chromosome differed from each other with respect to the total number of markers mapped, total cM distance and marker density. Variation in length varied from 104.9 cM (chromosome 4H) to a maximum length of 191.0 cM (chromosome 5H). The marker density was highest on chromosomes 3H, 5H, and 6H (2.3 cM, 2.7 cM, and 2.5 cM) which harbored 71, 72, and 57 markers, respectively. The lowest marker density was observed on chromosome 4H (7.5 cM) with 14 markers (Figure 4).

**Figure 4.**
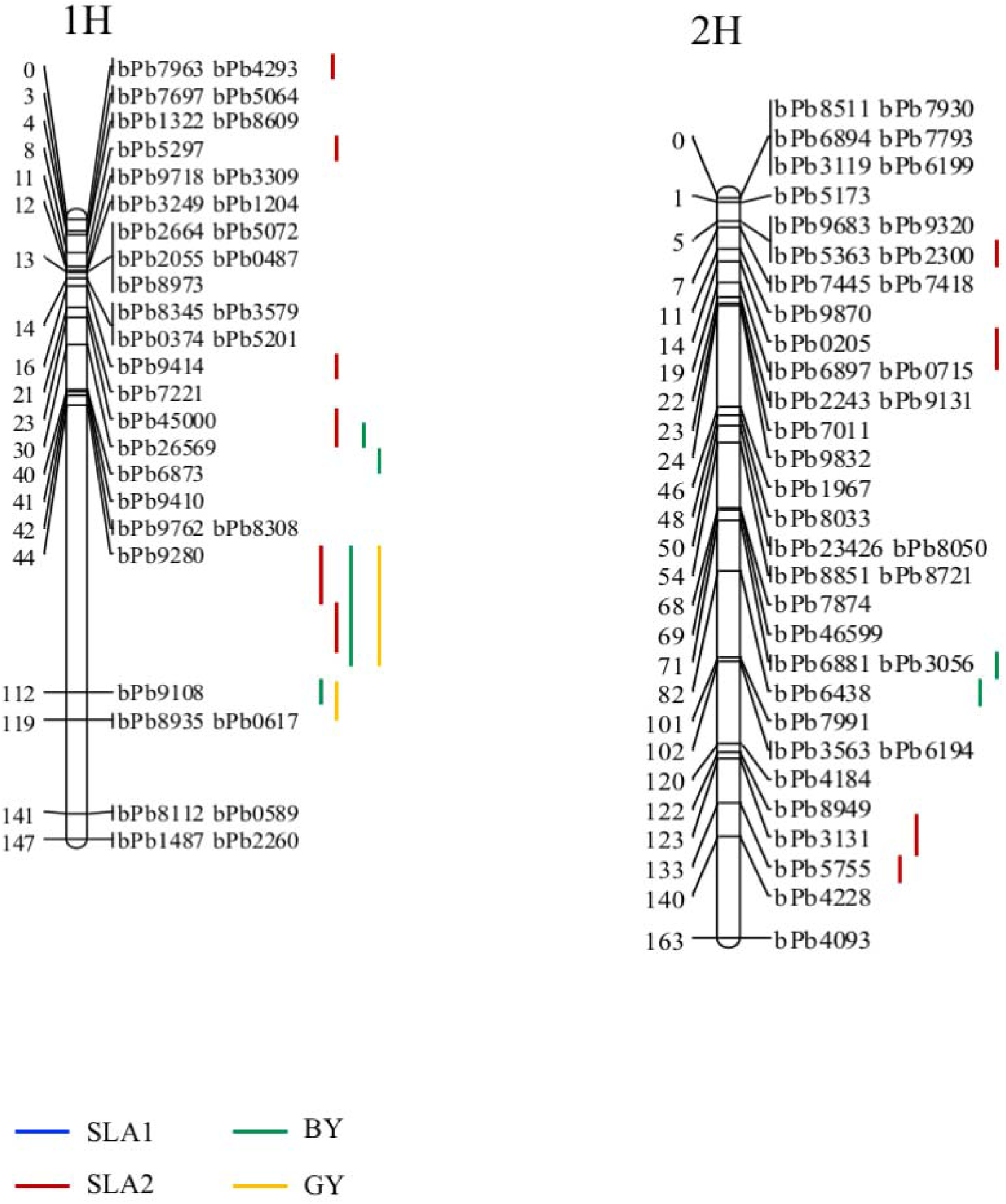
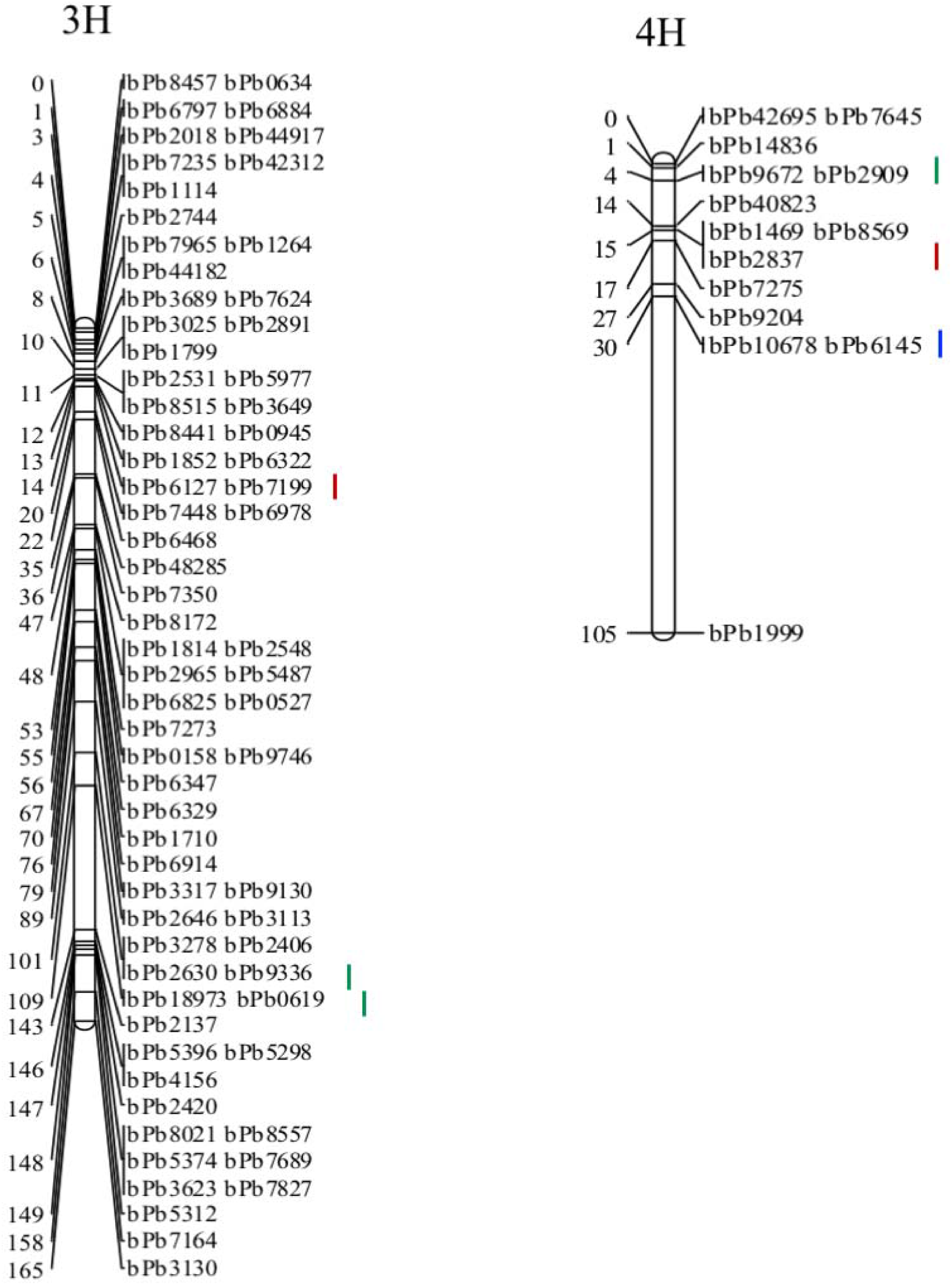
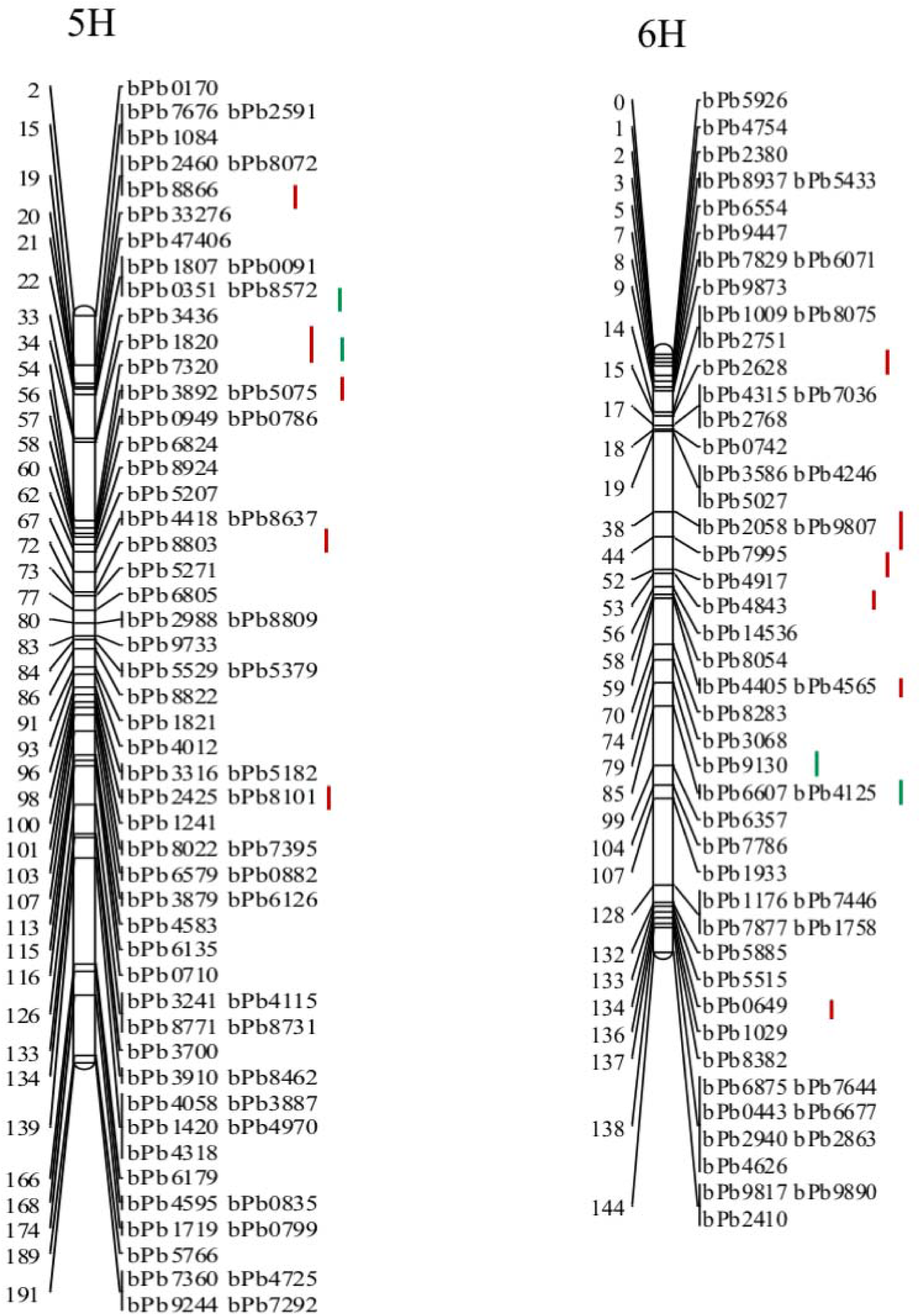
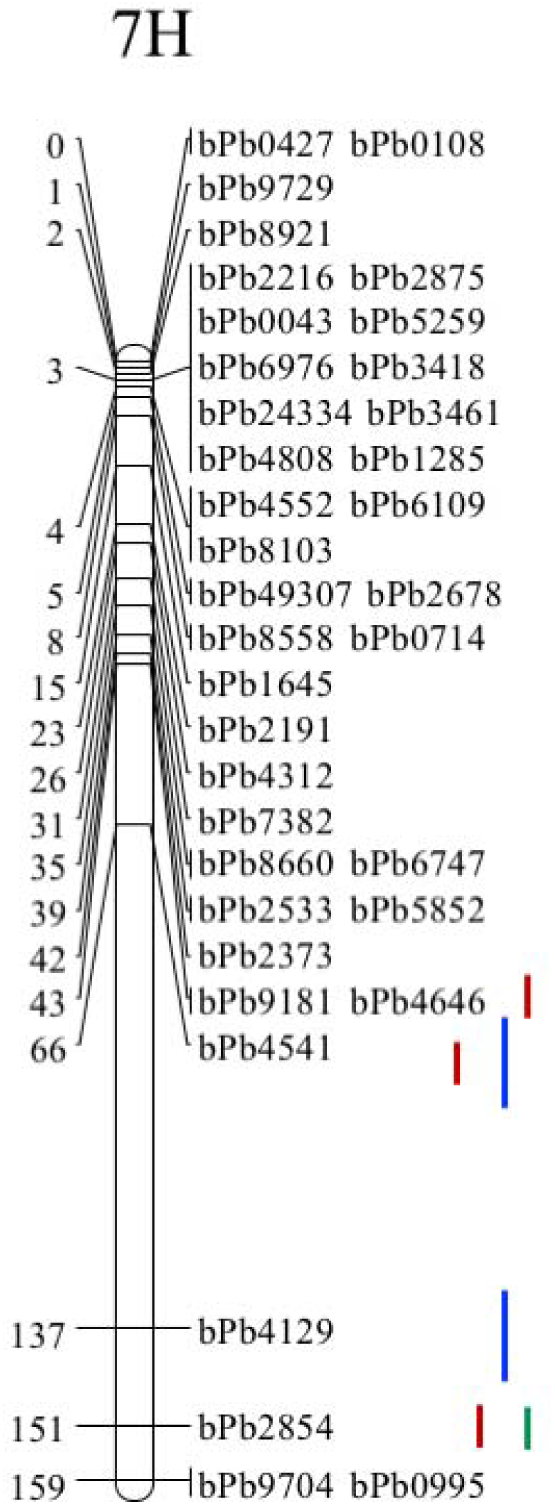
DArT linkage map for a Xena and H94061120 RIL population displaying markers with unique segregation patterns and associated consensus quantitative trait loci (QTLs) observed for early seedling vigour. The sequences of the relevant DArT makers that are associated with the QTLs in the present study presented in supplemetal Figure 1.

### 3.5. QTL detection and analysis

Forty-sixsignificant QTLs were detected for the 12 traits (Table 3). The CIM method revealed 35 significant QTLs (LOD > 3.0) and 11 tentative QTLs (2.5 < LOD < 3.0) associated with seedling vigour-related traits. Of the 46 QTLs identified, 26 were for SLA2 with LOD greater than 3.0, three were for SLA1 with LOD less than 3.0, 15 were for biomass with LOD, and two were for seed yield with LOD of 3.67 and 3.03. Of the 15 QTLs detected for biomass, 6 were significant (LOD > 3.0) and 8 were tentative (2.5 < LOD < 3.0). Five of the QTLs identified for SLA2 collocated with three QTLs for biomass on chromosome 1H, two QTLs for SLA1 on 7H, and one QTL for grain yield on 1H (Figure 3). Twenty-one of the QTLs detected for SLA2 accounted for 18% of the observed phenotypic variance (PVE), three for 19%, one for 17%, and one for 16% (Table 2). Two QTLs for SLA1 captured PVE of 13% and one for 16%. Thirteen of the QTLs identified for biomass accounted for 10% of the phenotypic variance, while the other two captured 18 and 9% of the variance, respectively. QTLs for seed yield accounted for 17.8 and 6.9% of the variance. The phenotypic variances explained by the two QTLs for seed yield were 18% and 7% (Table 3).

**Table 3.** Genetic characteristics of the QTLs (LOD > 2.5) related to the specific leaf area of the first leaf, specific leaf area of the second leaf, dry biomass, and grain yield

| Trait | Chr | Marker | Peak | Interval | LOD | ADD | PVE (%) | RecombiL | RecombiR |
| --- | --- | --- | --- | --- | --- | --- | --- | --- | --- |
| SLA1 | 4H | bPb6145 | 37.1 | 27.8 - 64.0 | 2.57 | 22.91 | 16.1 | 0.065 | 0.371 |
| SLA1 | 7H | bPb4541 | 70.9 | 65.7 - 100.5 | 2.50 | 25.43 | 12.7 | 0.048 | 0.366 |
| SLA1 | 7H | bPb4129 | 130.9 | 100.5 - 136.5 | 2.56 | 25.05 | 13.1 | 0.364 | 0.054 |
| SLA2 | 1H | bPb4293 | 2 | 0.6 - 3.2 | 6.31 | -75.91 | 17.8 | 0.020 | 0.013 |
| SLA2 | 1H | bPb5297 | 8.7 | 7.0 - 10.7 | 6.15 | -75.91 | 17.8 | 0.010 | 0.018 |
| SLA2 | 1H | bPb9414 | 18.1 | 17.1 - 18.7 | 6.54 | -75.91 | 17.8 | 0.020 | 0.023 |
| SLA2 | 1H | bPb45000 | 26.5 | 22.8 - 31.0 | 6.97 | -75.91 | 17.8 | 0.038 | 0.033 |
| SLA2 | 1H | bPb9280 | 45.5 | 43.0 - 76.5 | 7.18 | -75.91 | 17.8 | 0.020 | 0.368 |
| SLA2 | 1H | bPb9280 | 106.5 | 79.7 - 108.7 | 6.55 | -75.91 | 17.8 | 0.358 | 0.052 |
| SLA2 | 2H | bPb2300 | 6.2 | 3.8 - 7.0 | 5.76 | -75.91 | 17.8 | 0.010 | 0.007 |
| SLA2 | 2H | bPb0205 | 16.4 | 9.5 - 19.0 | 6.53 | -75.91 | 17.8 | 0.020 | 0.023 |
| SLA2 | 2H | bPb3131 | 127.7 | 122.0 - 132.8 | 7.11 | -75.91 | 17.8 | 0.048 | 0.053 |
| SLA2 | 2H | bPb5755 | 136.3 | 133.4 - 140.3 | 6.78 | -75.91 | 17.8 | 0.029 | 0.034 |
| SLA2 | 3H | bPb7199 | 16.7 | 13.9 - 20.2 | 6.78 | -75.91 | 17.8 | 0.029 | 0.032 |
| SLA2 | 4H | bPb2837 | 15.6 | 12.6 - 20.2 | 6.14 | -75.91 | 17.8 | 0.010 | 0.018 |
| SLA2 | 5H | bPb8866 | 17.3 | 17.0 - 21.9 | 3.06 | -73.41 | 16.8 | 0.000 | 0.028 |
| SLA2 | 5H | bPb1820 | 36.8 | 33.3 - 54.2 | 7.56 | -77.73 | 18.8 | 0.032 | 0.143 |
| SLA2 | 5H | bPb5075 | 55.1 | 55.1 - 60.5 | 5.48 | -77.73 | 18.8 | 0.000 | 0.019 |
| SLA2 | 5H | bPb8803 | 69.8 | 66.2 - 75.7 | 4.15 | -75.91 | 17.8 | 0.000 | 0.033 |
| SLA2 | 5H | bPb8101 | 97.5 | 94.0 - 103.5 | 5.74 | -77.40 | 18.6 | 0.000 | 0.023 |
| SLA2 | 6H | bPb2628 | 16.2 | 12.6 - 19.0 | 5.91 | -75.91 | 17.8 | 0.010 | 0.010 |
| SLA2 | 6H | bPb9807 | 41 | 19.0 - 44.3 | 6.69 | -75.91 | 17.8 | 0.029 | 0.024 |
| SLA2 | 6H | bPb7995 | 47.5 | 44.9 - 52.5 | 7.06 | -75.91 | 17.8 | 0.038 | 0.047 |
| SLA2 | 6H | bPb4843 | 53.6 | 51.8 - 58.8 | 6.21 | -75.91 | 17.8 | 0.010 | 0.020 |
| SLA2 | 6H | bPb4565 | 64.2 | 59.4 - 66.7 | 7.18 | -75.91 | 17.8 | 0.048 | 0.051 |
| SLA2 | 6H | bPb0649 | 134.1 | 128.3 - 138.4 | 4.22 | -75.91 | 17.8 | 0.000 | 0.020 |
| SLA2 | 7H | bPb9181 | 42.7 | 41.1 - 44.9 | 3.63 | -75.91 | 17.8 | 0.000 | 0.005 |
| SLA2 | 7H | bPb4541 | 67.9 | 60.7 - 73.3 | 5.79 | -72.67 | 16.5 | 0.020 | 0.373 |
| SLA2 | 7H | bPb2854 | 154.6 | 153.0 - 158.7 | 6.93 | -75.91 | 17.8 | 0.039 | 0.044 |
| BY | 1H | bPb45000 | 26.5 | 24.1 - 28.6 | 2.69 | 179.14 | 9.6 | 0.038 | 0.033 |
| BY | 1H | bPb26569 | 34.9 | 31.2 - 38.6 | 2.76 | 179.08 | 9.6 | 0.048 | 0.046 |
| BY | 1H | bPb9280 | 75.5 | 48.1 - 106.7 | 3.65 | -78.13 | 18.1 | 0.236 | 0.259 |
| BY | 1H | bPb9108 | 112 | 107.8 - 127.6 | 4.18 | 178.09 | 9.5 | 0.000 | 0.065 |
| BY | 2H | bPb3056 | 74.8 | 68.3 - 77.8 | 2.66 | 178.42 | 9.6 | 0.038 | 0.068 |
| BY | 2H | bPb6438 | 93.1 | 84.1 - 103.7 | 3.09 | 177.77 | 9.6 | 0.099 | 0.076 |
| BY | 3H | bPb48285 | 40.9 | 36.8 - 46.0 | 3.02 | 181.41 | 9.9 | 0.048 | 0.057 |
| BY | 3H | bPb9336 | 104.8 | 100.5 - 109.4 | 2.67 | 179.06 | 9.6 | 0.038 | 0.036 |
| BY | 3H | bPb0619 | 138.8 | 110.0 - 144.1 | 3.05 | 178.36 | 9.6 | 0.226 | 0.036 |
| BY | 4H | bPb2909 | 8.7 | 3.8 - 15.8 | 2.86 | 178.92 | 9.6 | 0.048 | 0.052 |
| BY | 5H | bPb8572 | 25.8 | 20.0 - 35.2 | 2.95 | 178.67 | 9.6 | 0.041 | 0.066 |
| BY | 5H | bPb1820 | 36.8 | 34.6 - 56.7 | 3.01 | 179.08 | 9.6 | 0.032 | 0.143 |
| BY | 6H | bPb9130 | 79.6 | 74.0 - 84.7 | 2.70 | 178.95 | 9.6 | 0.010 | 0.048 |
| BY | 6H | bPb4125 | 91.8 | 84.7 - 100.0 | 2.94 | 178.85 | 9.6 | 0.065 | 0.065 |
| BY | 7H | bPb2854 | 150.6 | 134.6 - 156.8 | 3.60 | 179.19 | 9.6 | 0.000 | 0.079 |
| GY | 1H | bPb9280 | 74.5 | 48.4 - 106.2 | 3.67 | -35.69 | 17.8 | 0.231 | 0.264 |
| GY | 1H | bPb9108 | 112 | 110.2 - 122.4 | 3.03 | 69.50 | 6.9 | 0.000 | 0.065 |

Chromosome (Chr), marker name (Marker), peak position in centiMorgan (cM) on chromosome, QTL interval, peak LOD, additive genetic effect on trait expression (ADD), percentage of phenotypic variation explained by an individual QTL (PVE), recombination frequency of the left flanking marker with the QTL (RecombiL), and recombination of the right flanking marker with the QTL (RecombiR)

There are six clusters of QTLs for the traits detected. On chromosome 1H, three clusters were detected: 1) QTLs for SLA2, biomass and yield were located within the same interval (44.0 to 108.0 cM) and share a common nearest marker, bpb9280; 2) QTLs for biomass and yield were located within the interval [107.8 to 127.6 cM] and share a common marker, bpb9108; 3) QTLs for SLA2 and biomass were found within the interval [22.8 to 31.0 cM] and share a common marker bpb45000. On chromosome 5H, QTLs for SLA2 and biomass were located within the interval [33.3 to 56.7 cM] and share a common marker, bpb1820. Furthermore, two clusters were detected on chromosome 7H: QTLs for SLA1 and SLA2 located within the interval [60.7 to 100.5 cM] with bpb4541 as the closest marker (Figure 4).

Positive colocations were identified for SLA2, biomass and seed yield with PVE values ranging from 17.8 to 18.1%. Three additional colocations between SLA2 and biomass were detected on chromosomes 5H and 7H with PVE values of 18.8% and 9.6%, respectively. One positive colocation was identified between SLA2 and SLA1 on chromosome 7H where the two QTLs exhibited PVE values of 16.5% and 12.7%. QTLs interval length ranged from 1.6 to 58.6 cM, averaging 15.9 cM. Most of the QTLs detected contained one or two markers. Chromosome 1H is likely to play a key role in seedling vigour and yield determination in barley.

## 4. Discussion

Seed vigour depends on the physiological and genetic potential of the seeds and on the conditions encountered during storage (Rajjou et al., 2008). It has a strong influence on plant stand establishment, and the production of high vigour seeds to stabilize crop yield is a challenge for crop breeders. So that, the amount of nutrition production essentially depends on the first few leaves of the seedling (Katsvairo et al., 2003). Quantitative trait loci mapping is an efficient method to analyze genetically complex traits and genotype-phenotype mapping (Zhu et al., 2008). The current study focused on characterizing early seedling vigour (ESV) as complementary physiological traits that could be used to improve the water use efficiency and yield stability of barley under low moisture conditions. Early seedling vigour in this study refers to an increase in seedling leaf characteristics that would help plants to form a canopy more quickly and reduce water evaporation from the soil surface as well as increase canopy transpiration. Therefore, the most crucial step in QTL mapping for seedling vigour is the evaluation and screening of the quantitative traits regard to ESV (Mangin et al., 1994). Few studies on the molecular mechanisms underlying seed vigour traits in barley have been reported. In this study, 185 RILs were evaluated for ESV under rain-fed conditions during two years.

### 4.1. Seedling vigor-related traits

Twelve seedling related traits identified as representative indicators for seedling vigour during early seedling development. The significant variation of RILs for ESV and seedling growth was observed. Specific leaf area (SLA) of the first leaf is highly correlated with SLA of the second leaf. However, their contribution to ESV is relative, with SLA of the second leaf being the most significant contributor. In this study, we found a negative correlation between leaf area and leaf dry weight content of the second leaf, suggesting that high or low SLA depends on the significance of the correlation between the two components based on the genotype, species, and environment. The analysis of this type of trait has received little attention from geneticists (Yin et al., 1999). An increase in leaf area does not necessarily translate to a proportional increase in dry matter content. This may explain the high SLA observed for the second leaf. A high SLA would result in greater water loss due to the larger leaf area exposed to ambient air. However, larger leaf areas with greater biomass allocation to the leaves are often associated with high relative growth rate which might lower SLA (Poorter et al., 2012). Considering that rapid dry matter production during early seedling growth is an important aspect of seedling vigour and the most common measure, high early vigour might coincide with a high relative growth rate in the early stages of seedling development as suggested by (ter Steege et al., 2005). On the other hand, a positive correlation was observed between leaf area and leaf dry weight content of the first leaf, suggesting that an increase in leaf area would result to an increase in dry matter content. Leaf traits may reflect the adaptation mechanisms of plants to the environment (Tian et al., 2016). Therefore, a lower SLA could be associated with smaller leaf area, which may be shown to reduce water loss due to evapotranspiration on the leaf surface. However, in some succulent plants with poor seedling vigour that are common in tropical regions, low SLA may be associated with low leaf dry-matter and high leaf thickness. As a consequence of these variations, SLA and its components are often related to each other and to productivity. Our results confirmed previous findings stating SLA as a suitable trait for the selection of plants with good early seedling vigour in cereals (Rebetzke et al., 2008). Hence, Specific leaf area, the ratio of leaf area to leaf dry mass, is a key functional trait of plants underlying variation in growth rate among species (Pérez-Harguindeguy et al., 2013; Flores et al., 2014). SLA is also a major trait in the worldwide leaf economics spectrum, which reflects the range of fast to slow returns on nutrient and dry mass investment in leaves among species (Flores et al., 2014). We found that the width of the first leaf is highly correlated with leaf area. It was suggested that leaf width of the first leaf should integrate embryo size and SLA, and this would be a simple way to screen and select for high early vigour. Wang et al., later established that seedling leaf width was highly heritable and had a high genetic correlation with total leaf area in wheat during the vegetative stage (Wang et al., 2011). Sundgren et al., also showed the importance of both embryo size and SLA in determining vigour among wheat lines (Sundgren et al., 2018).

We found that SLA2 contributed to the highest seedling vigour and seed yield compared to the rest of seedling-traits, suggesting that SLA2 is important for achieving higher seed yield. In the current study, the knowledge of dgree and nature of the variability among the RILs for each traits is very necessary for making simultaneous selection on more number of traits to make significant improvement seedling traits in barley. Consequently, the hight correspondence among genotypic and phenotypic coefficient of variation for most the recorded traits showed that these characters hight influenced by the environment. Moreover, the highest observed genotypic and phenotypic coefficient of variation, indicates that selection can be applied on the traits to isolate more promising RILs. The studies traits LL1, LL2, LA1 and LDW1 had high heritability (>60%) in broad sense (h^2^_b_), indicates that these characters could be easily improved by selection. Leaf length (LL1) and area (LA1) of the first leaf showed moderately high; 84.65 % and 71.47 % heritability respectively with a genetic advance of at 5% 1.24 and 0.88 (Table 2). Since the heritability estimates the relative contributions of differences in genetic and non-genetic factors to the total phenotypic variance in the studied population. It is an important concept in quantitative genetics, particularly in selective breeding. Given that, theses results indicating the possibility of additive gene effect for the expression of these traits. Therefore selection would be effective for improving this traits. Estimates of GA for GY/RIL was 2038.06 gm^-2^ / RIL indicating that whenever we select the best, 5% high yielding genotypes as parents, average grain yield/plant of progenies could be improved by 2038.06 gm-2 / RIL.

### 4.2 Map Chatacteristics

Over the last decade, barley has been the subject of extensive mapping studies with the DArT technology (Rodríguez-Suárez et al., 2012; Liu et al., 2015). Thus, DArT markers have proved to be very useful to detect chromosome substitutions in the breeding program (Li et al., 2014). By the end of 2012, a total of 2,032 DArT markers have been mapped to 646 unique positions (bins) in *Hordeum chilense* recombinant inbred line (RIL) population (Rodríguez-Suárez et al., 2012). Besides, the genetic map is very useful as the basis to develop comparative genomics studies with barley and model species (Rodríguez-Suárez et al., 2012). ESV is a quantitative trait which is controlled by many genes, and has been widely reported to provide improved productivity of cereal crops in semi-arid environments, and particularly those environments in which rainfall pattern are rare (Botwright et al., 2002). Mapping of QTLs related to ESV can enable dissection of their genetic control and molecular mechanism, leading to the possibility to develop new varieties with improved ESV and enhanced yield.

In the present study, the map based on segregations in progeny from a cross of the Xena and H94061120 cultivars representing separate gene defined positions of 328 polymorphic DArT markers. These markers were distributed on 7 chromosomes spanning a cumulative distance of 1075 cM with an average marker density of 3.3 cM. The high density of the genetic maps showed that there was large genetic diversity and variation in the analyzed phenotypic traits. The high efficiency of mapping markers is of paramount importance in species that have a variable genomic constitution and possible chromosome rearrangements (Wójcik-Jagła et al., 2013). Among the 7 chromosomes, each chromosome differed from the other with respect to length and marker distribution as a result of which some chromosomes were densely populated (3H, 5H, 6H), while other exhibited few markers (1H, 2H, 4H, 7H) which could be explained by the fact that DArT markers are randomly distributed in plant genomes. Most of the markers were located on the centromeric region. This may be due to the lower recombination frequency in these regions. Previous analyses of DArT sequences in other species indicated that DArT markers tend to be located in gene-rich regions (Gawroński et al., 2016).

### 4.3. QTL and DArT Marker Discovery

According to developmental genetics, different QTLs may have different expression dynamics during trait development (Wang et al., 2017). Many previous investigates focused mainly on late-growth stages, where analysis was limited to the performance of a trait at a fixed time or stage of ontogenesis (Broughton et al., 2015; Wang et al., 2016; Zhou et al., 2016; Wen et al., 2018). The present study provides insight into the genetic and physiological relationships among traits that contribute to early seedling vigour in barley. We assessed five ESV characteristics at third leaf (LW, LL, LA, LDW, SLA) at two stages of seedling growth (first and second leaf stage), to reveled the QTLs expression patterns (Wang et al., 2017).

We found that multiple loci have been identified for SLA, biomass, and seed yield explaining a high phenotypic variance. A total of 46 QTLs detected, 29 were for SLA (26 for SLA2 and 3 for SLA1), 15 for biomass, and 2 for seed yield, suggesting that by selecting based on SLA, good seedling vigour, rapid establishment and thus higher yield could be attained (Table 2 and Figure 4). (Gawroński et al., 2016) used a genetic map based on 262 SSR markers and found a 27 QTL for seed vigor in rice. In our study, we found that most of QTLs were located on chromosomes 1H, 5H and 6H [48]. It is noteworthy all seven chromosomes, QTLs for early vigour traits were detected, and, in some cases, we found that many QTLs controlling multiple traits were located at the same or overlapping marker interval on the same chromosomes. For instance, a pleiotropic QTL, detected between DArT markers bPb-9280 and bPb-9108 on chromosome 1H, was related to SLA2, biomass, and yield. It may presently be inaccurate to determine whether one gene affects a range of traits or whether there are several genes clustered in the same region that act upon different related traits. Considering all information here, we suggested that this region may be a credible region for cluster of QTLs (Herwaarden et al., 1998). This may imply an existence of pleiotropic QTL or tightly linked QTL in our study. Xu et al., 2007 suggested that if two QTL peaks are located very close to each other, and the 1-LOD support intervals completely or mostly overlapped, these two QTLs would be regarded as a single QTL having pleiotropic effects. Lui et al., reported QTL for morphological and physiological traits of flag leaf (net photosynthetic rate, stomatal conductance, flag leaf area, flag leaf length, flag leaf width, relative chlorophyll content and leaf nitrogen concentration) in the similar region (Mikołajczak et al., 2016). Moreover, clustering of QTLs for related traits has been observed in other studies (e.g. in wheat [*Triticum aestivum*] and rice [*Oryza sativa*]; (ter Steege et al., 2005), and it was suggested that these QTLs clusters represent gene clusters that are separated by regions with noncoding sequences. An alternative explanation for clustering of QTLs for traits at different organizational levels in the plants is that of a mechanistic dependency rather than a genetic dependency between traits. For example, the colocation of QTLs for SLA2 with QTLs for biomass and seed yield might be due to the fact that one or more genes in that region affect SLA2; consequently, this chromosomal region affects the resulting biomass and seed yield. The colocalization of QTLs for SLA1, SLA2 with biomass and yield is not unexpected, because these traits are associated with photosynthesis and transpiration in barley. The relationship among these traits indicates that these parameters may not be independent but interacting. They may be co-regulated for the protection of photosynthetic apparatus, an important factor in dry matter accumulation.

The 26 significant QTLs for SLA2 detected on all chromosomes explained 16.5 – 18.8 % of the observed phenotypic variance with the larger contribution from markers bPb5075 and bPb1820 (5H). In a population of fodder barley reported in previous study (Sandve et al., 2011) identified 4 QTLs for water content stress index on chromosome 2H and 5H, explaining 37.3% of the phenotypic variance with the largest contribution from marker bPb-5075 (5H) and bPb-1976 (2H). Therefore, QTL for SLA2 is collocated with QTLs for water content stress index, suggesting that the DArT marker bPb-5075 could be useful for marker-assisted selection (MAS) in barley for breeding for early seedling vigour. Similarly, two QTLs for SLA2 and biomass (5H), which peaked at DArT marker bPb-2857 collocated with a QTL for *QPSII.sthb* related to early short-time drought tolerance in barley detected on chromosome 5H (Rodríguez-Suárez et al., 2012).

QTL clustering in barley was repeatedly reported in some studies (Hori et al., 2007) (Mikołajczak, 2016; Zhou et al., 2016). In our study, we demonstrated several significant QTL clusters of ESV characteristics in barley under rainfall condition. The more QTLs of seedling vigour-related traits that are identified, they more improve breeding efficiency by marker assisted selection (MAS). Although many QTLs were identified for most of the seedling vigour traits considered in this study, we did not detect significant QTLs for leaf area (LA), leaf length (LL), and leaf width (LW) of the first and second leaves. These traits may be complex physiological traits that presumably are under control of many loci on the genome. QTLs with small effects on the overall complex traits are difficult to detect so that for such traits, usually only a few major QTLs are identified (Gawroński et al., 2016).

## Conclusion

This study identified 46 significant QTLs for ESV related traits. Of which 26 were detected for SLA2, 15 for biomass, 3 for SLA1, and 2 for yield. QTLs for yield harbor 2 QTLs for SLA2 and 2 QTLs for biomass. QTLs controlling SLA2 were distributed on all the seven chromosomes and explained on the average of 16.5 – 18.8% of the phenotypic variance. This suggests that SLA2 traits may contribute to ESV in barley, but further study using genetically different populations is required. Considering the importance of SLA2 and SLA1 for rapid plant establishment and putative influence on later seed yield, information gained in this study could be used to improve early seeding vigour, plant establishment, and seed yield. Co-localization of QTLs for different traits suggests a common genetic basis of regulation of the traits and point to the regulation of these traits either by tightly linked genes or by pleiotropic regulation. An assessment of the phenotypic relationships between early vigour and related traits indicated strong correlations between early vigour and grain yield.

## Authors contributions

AA; Conceived of the study. LC; Performed the data analysis, carried out map construction, interpreted data, and drafted the manuscript. SE; Conducted the field experiment and collected the phenotypic data. AE and TK; Performed correlation and statistical analysis wrote the manuscript, organized figures and additional files. All authors read and approved of the final manuscript.

## Confliction of interest statement

The authors declare that the research was conducted in the absence of any commercial or financial relationships that could be construed as a potential conflict of interest.

## Acknowledgment

This study was funded by Alberta Crop Industry Development Fund (ACIDF). The matching funding provided by the Brewing and Malting Barley Research Institute is acknowledged. We are grateful to the staff of Field Crop Development Centre (FCDC), Lacombe, for providing materials used in this study. In-kind support provided by 20/20 Seed Labs Inc. for seed vigour analysis is appreciated. We are also grateful to 20/20 Seed Labs Inc. for hosting the Alberta Ingenuity Industrial R&D associate who worked on the early seedling vigour related traits. We thank Professor. Calvin O. Qualset – University of California, Davis, for his critical comments and helpful suggestions.

---

**Supplemtial Table 1.** Mean performance for twelve early seedling vigour-related traits of 185 RILs of barley

| Genotypes | LL1 | LL2 | LW1 | LW2 | LA1 | LA2 | LDW1 | LDW2 | SLA1 | SLA2 | BY | GY |
| --- | --- | --- | --- | --- | --- | --- | --- | --- | --- | --- | --- | --- |
| 1 | 2.32 | 10.66 | 0.53 | 0.73 | 3.13 | 6.33 | 0.12 | 0.02 | 171.35 | 251.69 | 541.20 | 231.00 |
| 2 | 2.29 | 10.09 | 0.60 | 0.57 | 3.32 | 4.65 | 0.10 | 0.02 | 164.24 | 204.95 | 499.43 | 205.43 |
| 3 | 2.22 | 9.89 | 0.52 | 0.63 | 2.83 | 5.25 | 0.10 | 0.02 | 151.86 | 273.00 | 475.43 | 200.67 |
| 4 | 3.11 | 7.88 | 0.81 | 2.27 | 6.08 | 5.53 | 0.10 | 70.63 | 242.17 | 201.98 | 384.87 | 186.67 |
| 5 | 2.49 | 10.20 | 0.51 | 0.67 | 3.15 | 5.56 | 0.11 | 0.02 | 179.80 | 240.58 | 439.00 | 174.67 |
| 6 | 2.50 | 10.69 | 0.57 | 0.71 | 3.46 | 6.11 | 0.12 | 0.02 | 153.71 | 250.31 | 501.73 | 213.13 |
| 7 | 2.27 | 10.57 | 0.61 | 0.68 | 3.35 | 5.79 | 0.11 | 0.02 | 193.81 | 265.27 | 470.43 | 170.63 |
| 8 | 2.58 | 12.45 | 0.61 | 0.74 | 3.81 | 7.40 | 0.13 | 0.03 | 188.09 | 287.25 | 524.33 | 202.33 |
| 9 | 3.42 | 9.17 | 0.62 | 1.81 | 5.20 | 5.07 | 0.12 | 53.85 | 186.27 | 157.34 | 458.60 | 230.20 |
| 10 | 2.77 | 13.10 | 0.78 | 0.81 | 5.25 | 8.52 | 0.17 | 0.03 | 188.33 | 252.89 | 500.37 | 186.77 |
| 11 | 2.33 | 10.75 | 0.54 | 0.55 | 3.01 | 4.79 | 0.10 | 0.02 | 153.85 | 215.05 | 501.70 | 215.30 |
| 12 | 2.75 | 11.91 | 0.61 | 0.70 | 4.01 | 6.62 | 0.14 | 0.03 | 171.75 | 237.65 | 423.50 | 166.23 |
| 13 | 2.46 | 12.01 | 0.64 | 0.72 | 3.80 | 7.02 | 0.15 | 0.03 | 171.13 | 228.83 | 509.03 | 215.93 |
| 14 | 2.66 | 12.69 | 0.66 | 0.78 | 4.25 | 7.98 | 0.14 | 0.03 | 182.09 | 286.52 | 422.67 | 168.93 |
| 15 | 2.22 | 11.24 | 0.53 | 0.57 | 2.83 | 5.16 | 0.11 | 0.02 | 166.63 | 227.21 | 539.50 | 233.80 |
| 16 | 2.88 | 11.39 | 0.59 | 0.65 | 4.09 | 5.88 | 0.13 | 0.03 | 166.42 | 224.60 | 511.57 | 195.63 |
| 17 | 2.51 | 12.03 | 0.65 | 0.76 | 3.96 | 7.38 | 0.14 | 0.03 | 182.27 | 258.71 | 554.63 | 218.83 |
| 18 | 2.77 | 12.17 | 0.65 | 0.75 | 4.33 | 7.33 | 0.16 | 0.03 | 156.87 | 234.11 | 611.83 | 229.80 |
| 19 | 2.90 | 12.56 | 0.52 | 0.66 | 3.63 | 6.69 | 0.14 | 0.03 | 148.84 | 246.30 | 436.80 | 165.40 |
| 20 | 2.38 | 11.42 | 0.59 | 0.67 | 3.40 | 6.23 | 0.12 | 0.02 | 175.93 | 255.05 | 661.53 | 274.60 |
| 21 | 2.63 | 11.75 | 0.57 | 0.64 | 3.63 | 6.09 | 0.12 | 0.02 | 165.17 | 245.58 | 525.10 | 217.63 |
| 22 | 2.28 | 11.41 | 0.52 | 0.59 | 2.98 | 5.40 | 0.11 | 0.02 | 161.03 | 237.60 | 429.03 | 174.87 |
| 23 | 2.59 | 13.18 | 0.61 | 0.74 | 3.83 | 7.81 | 0.13 | 0.03 | 179.81 | 304.38 | 481.30 | 199.37 |
| 24 | 2.33 | 11.29 | 0.52 | 0.69 | 2.95 | 6.40 | 0.13 | 0.03 | 160.42 | 250.18 | 519.83 | 220.90 |
| 25 | 2.53 | 10.96 | 0.63 | 0.76 | 3.87 | 6.71 | 0.11 | 0.03 | 180.70 | 261.63 | 467.70 | 201.57 |
| 26 | 2.20 | 10.49 | 0.59 | 0.67 | 3.16 | 5.66 | 0.11 | 0.02 | 179.79 | 266.36 | 586.47 | 227.03 |
| 27 | 2.69 | 12.31 | 0.67 | 0.79 | 4.35 | 7.79 | 0.14 | 0.03 | 194.48 | 284.57 | 495.47 | 200.30 |
| 28 | 2.35 | 10.43 | 0.55 | 0.63 | 3.08 | 5.31 | 0.11 | 0.02 | 143.23 | 225.19 | 534.63 | 227.97 |
| 29 | 2.59 | 12.31 | 0.60 | 0.65 | 3.70 | 6.42 | 0.13 | 0.03 | 165.46 | 239.15 | 468.37 | 180.73 |
| 30 | 2.75 | 12.30 | 0.60 | 0.67 | 3.95 | 6.58 | 0.12 | 0.02 | 193.72 | 270.96 | 531.83 | 201.73 |
| 31 | 2.62 | 12.05 | 0.56 | 0.66 | 3.54 | 6.39 | 0.12 | 0.02 | 170.60 | 262.46 | 500.80 | 180.43 |
| 32 | 2.24 | 11.79 | 0.58 | 0.65 | 3.18 | 6.15 | 0.13 | 0.03 | 166.59 | 242.22 | 374.13 | 141.53 |
| 33 | 2.19 | 10.06 | 0.60 | 0.65 | 3.22 | 5.30 | 0.11 | 0.02 | 167.26 | 251.36 | 420.53 | 180.27 |
| 34 | 2.61 | 11.97 | 0.61 | 0.77 | 3.85 | 7.43 | 0.14 | 0.03 | 178.40 | 274.29 | 526.93 | 210.33 |
| 35 | 2.54 | 12.65 | 0.64 | 0.75 | 3.86 | 7.61 | 0.13 | 0.03 | 197.20 | 299.89 | 418.10 | 160.30 |
| 36 | 2.73 | 12.87 | 0.63 | 0.76 | 4.21 | 7.87 | 0.15 | 0.03 | 176.04 | 253.42 | 532.03 | 210.77 |
| 37 | 2.55 | 12.39 | 0.59 | 0.69 | 3.66 | 6.93 | 0.13 | 0.03 | 176.43 | 264.54 | 558.63 | 225.03 |
| 38 | 2.67 | 11.43 | 0.62 | 0.77 | 4.04 | 7.19 | 0.13 | 0.03 | 182.71 | 293.10 | 502.53 | 198.00 |
| 39 | 2.55 | 10.84 | 0.62 | 0.71 | 3.81 | 6.21 | 0.11 | 0.02 | 185.82 | 272.61 | 482.97 | 193.33 |
| 40 | 2.40 | 11.37 | 0.60 | 0.71 | 3.46 | 6.43 | 0.12 | 0.02 | 168.82 | 257.88 | 595.83 | 244.30 |
| 41 | 2.33 | 10.84 | 0.68 | 0.68 | 3.84 | 6.04 | 0.12 | 0.02 | 186.02 | 261.02 | 475.30 | 181.30 |
| 42 | 2.77 | 11.32 | 0.52 | 0.60 | 3.46 | 5.43 | 0.13 | 0.03 | 161.72 | 211.42 | 484.20 | 201.20 |
| 43 | 2.67 | 11.01 | 0.72 | 0.73 | 4.55 | 6.29 | 0.14 | 0.03 | 189.11 | 230.11 | 428.93 | 167.00 |
| 44 | 2.18 | 11.26 | 0.53 | 0.61 | 2.79 | 5.51 | 0.10 | 0.02 | 163.70 | 260.73 | 449.73 | 178.07 |
| 45 | 2.75 | 11.83 | 0.64 | 0.78 | 4.20 | 7.38 | 0.16 | 0.03 | 146.39 | 231.18 | 480.33 | 185.03 |
| 46 | 2.43 | 12.15 | 0.65 | 0.72 | 3.76 | 7.02 | 0.13 | 0.03 | 175.30 | 271.70 | 481.87 | 199.70 |
| 47 | 2.59 | 12.68 | 0.70 | 0.80 | 4.35 | 8.13 | 0.16 | 0.03 | 193.70 | 256.89 | 463.40 | 166.70 |
| 48 | 2.20 | 11.09 | 0.54 | 0.59 | 2.87 | 5.18 | 0.12 | 0.02 | 155.18 | 218.41 | 483.50 | 197.00 |
| 49 | 2.31 | 11.03 | 0.63 | 0.72 | 3.53 | 6.34 | 0.11 | 0.02 | 182.77 | 288.87 | 587.17 | 224.67 |
| 50 | 2.22 | 11.47 | 0.57 | 0.60 | 2.97 | 5.49 | 0.11 | 0.02 | 163.10 | 254.74 | 475.77 | 186.00 |
| 51 | 2.26 | 10.75 | 0.63 | 0.67 | 3.45 | 5.73 | 0.10 | 0.02 | 187.83 | 282.04 | 578.07 | 240.93 |
| 52 | 2.08 | 10.60 | 0.53 | 0.66 | 2.67 | 5.64 | 0.11 | 0.02 | 160.10 | 245.02 | 528.70 | 205.97 |
| 53 | 2.42 | 12.83 | 0.58 | 0.67 | 3.23 | 6.80 | 0.15 | 0.03 | 174.54 | 228.46 | 466.70 | 186.57 |
| 54 | 2.39 | 11.43 | 0.53 | 0.68 | 3.08 | 6.22 | 0.12 | 0.02 | 166.88 | 268.01 | 492.43 | 195.70 |
| 55 | 2.72 | 11.93 | 0.61 | 0.71 | 3.98 | 6.78 | 0.13 | 0.03 | 172.05 | 256.49 | 501.10 | 203.10 |
| 56 | 2.95 | 13.62 | 0.67 | 0.81 | 4.75 | 8.83 | 0.16 | 0.03 | 187.04 | 271.91 | 523.40 | 202.03 |
| 57 | 2.72 | 12.23 | 0.54 | 0.62 | 3.56 | 6.09 | 0.13 | 0.03 | 160.10 | 227.25 | 277.70 | 96.40 |
| 58 | 2.54 | 12.41 | 0.63 | 0.73 | 3.83 | 7.29 | 0.14 | 0.03 | 193.63 | 261.15 | 570.33 | 221.03 |
| 59 | 2.54 | 10.98 | 0.57 | 0.65 | 3.54 | 5.79 | 0.12 | 0.02 | 150.12 | 242.50 | 578.50 | 249.33 |
| 60 | 2.35 | 11.22 | 0.52 | 0.56 | 2.94 | 5.04 | 0.13 | 0.03 | 136.25 | 196.07 | 430.00 | 189.30 |
| 61 | 2.34 | 11.05 | 0.59 | 0.62 | 3.29 | 5.46 | 0.13 | 0.03 | 180.36 | 218.58 | 705.70 | 296.13 |
| 62 | 2.54 | 12.25 | 0.69 | 0.74 | 4.27 | 7.33 | 0.13 | 0.03 | 203.11 | 273.86 | 547.00 | 229.40 |
| 63 | 2.53 | 11.93 | 0.61 | 0.68 | 3.68 | 6.55 | 0.13 | 0.03 | 173.72 | 244.34 | 433.87 | 179.40 |
| 64 | 2.36 | 11.46 | 0.55 | 0.65 | 3.19 | 5.94 | 0.11 | 0.02 | 163.42 | 263.47 | 512.13 | 215.50 |
| 65 | 2.45 | 10.50 | 0.53 | 0.69 | 3.25 | 5.85 | 0.12 | 0.02 | 160.11 | 242.81 | 461.83 | 177.53 |
| 66 | 2.42 | 10.72 | 0.59 | 0.65 | 3.41 | 5.61 | 0.10 | 0.02 | 184.83 | 276.60 | 454.03 | 179.93 |
| 67 | 2.52 | 11.27 | 0.60 | 0.64 | 3.66 | 5.79 | 0.12 | 0.02 | 182.73 | 244.21 | 589.27 | 227.57 |
| 68 | 2.76 | 12.21 | 0.70 | 0.77 | 4.86 | 7.57 | 0.13 | 0.03 | 165.03 | 287.86 | 613.43 | 254.23 |
| 69 | 2.17 | 11.13 | 0.55 | 0.59 | 2.92 | 5.30 | 0.10 | 0.02 | 174.81 | 268.20 | 586.23 | 225.30 |
| 70 | 2.38 | 10.03 | 0.55 | 0.71 | 3.20 | 5.88 | 0.10 | 0.02 | 171.90 | 292.14 | 391.00 | 157.50 |
| 71 | 2.32 | 10.59 | 0.72 | 0.72 | 4.11 | 6.23 | 0.10 | 0.02 | 223.35 | 277.48 | 491.07 | 200.47 |
| 72 | 3.04 | 13.02 | 0.69 | 0.76 | 5.11 | 7.92 | 0.16 | 0.03 | 172.33 | 245.93 | 472.40 | 196.67 |
| 73 | 2.13 | 10.91 | 0.65 | 0.67 | 3.38 | 5.89 | 0.11 | 0.02 | 182.00 | 259.00 | 494.60 | 193.07 |
| 74 | 2.34 | 10.21 | 0.58 | 0.60 | 3.24 | 4.89 | 0.09 | 0.02 | 181.14 | 258.51 | 533.43 | 228.43 |
| 75 | 2.36 | 10.21 | 0.63 | 0.69 | 3.56 | 5.48 | 0.09 | 0.02 | 186.57 | 293.52 | 540.23 | 183.73 |
| 76 | 2.35 | 11.16 | 0.58 | 0.72 | 3.28 | 6.53 | 0.11 | 0.02 | 196.68 | 284.35 | 488.40 | 191.10 |
| 77 | 2.49 | 11.20 | 0.54 | 0.62 | 3.24 | 5.57 | 0.11 | 0.02 | 174.37 | 261.24 | 490.00 | 218.10 |
| 78 | 2.71 | 12.06 | 0.58 | 0.62 | 3.79 | 6.00 | 0.13 | 0.03 | 161.16 | 225.90 | 521.40 | 221.50 |
| 79 | 2.43 | 10.95 | 0.61 | 0.73 | 3.54 | 6.38 | 0.11 | 0.02 | 180.95 | 295.83 | 632.53 | 257.17 |
| 80 | 2.38 | 10.35 | 0.52 | 0.62 | 2.99 | 5.25 | 0.11 | 0.02 | 149.76 | 238.03 | 481.60 | 209.70 |
| 81 | 2.60 | 11.53 | 0.59 | 0.67 | 3.72 | 6.22 | 0.11 | 0.02 | 182.36 | 278.81 | 562.37 | 228.90 |
| 82 | 2.96 | 13.33 | 0.61 | 0.69 | 4.35 | 7.35 | 0.14 | 0.03 | 169.89 | 256.03 | 495.93 | 186.27 |
| 83 | 2.60 | 12.59 | 0.51 | 0.63 | 3.20 | 6.37 | 0.13 | 0.03 | 156.20 | 250.30 | 449.17 | 187.40 |
| 84 | 2.63 | 10.88 | 0.52 | 0.67 | 3.31 | 5.91 | 0.11 | 0.02 | 161.35 | 265.16 | 592.00 | 255.07 |
| 85 | 2.52 | 10.54 | 0.58 | 0.70 | 3.47 | 5.90 | 0.11 | 0.02 | 180.70 | 231.40 | 345.83 | 126.37 |
| 86 | 2.61 | 11.38 | 0.66 | 0.67 | 4.12 | 6.14 | 0.11 | 0.02 | 195.08 | 275.04 | 575.40 | 251.23 |
| 87 | 2.55 | 10.31 | 0.64 | 0.73 | 3.94 | 6.06 | 0.10 | 0.02 | 196.06 | 310.16 | 468.77 | 188.70 |
| 88 | 2.39 | 11.05 | 0.59 | 0.62 | 3.44 | 5.55 | 0.12 | 0.02 | 180.40 | 233.27 | 560.80 | 236.87 |
| 89 | 2.41 | 11.57 | 0.63 | 0.66 | 3.67 | 6.11 | 0.13 | 0.03 | 165.64 | 242.23 | 449.07 | 178.67 |
| 90 | 3.03 | 12.07 | 0.67 | 0.74 | 4.82 | 7.18 | 0.12 | 0.02 | 190.11 | 301.82 | 590.13 | 228.97 |
| 91 | 2.86 | 12.98 | 0.66 | 0.78 | 4.57 | 8.14 | 0.15 | 0.03 | 192.48 | 264.62 | 596.83 | 233.13 |
| 92 | 2.59 | 11.70 | 0.62 | 0.66 | 3.86 | 6.19 | 0.14 | 0.03 | 174.45 | 221.55 | 258.80 | 83.70 |
| 93 | 2.53 | 12.02 | 0.65 | 0.75 | 4.11 | 7.36 | 0.15 | 0.03 | 179.49 | 250.52 | 587.10 | 242.23 |
| 94 | 2.97 | 11.82 | 0.54 | 0.64 | 3.83 | 6.00 | 0.12 | 0.02 | 156.22 | 245.09 | 526.70 | 205.90 |
| 95 | 2.66 | 11.93 | 0.61 | 0.74 | 3.93 | 7.10 | 0.13 | 0.03 | 182.45 | 268.95 | 441.17 | 166.60 |
| 96 | 2.67 | 10.82 | 0.56 | 0.67 | 3.75 | 5.87 | 0.12 | 0.02 | 189.47 | 247.52 | 506.10 | 211.67 |
| 97 | 2.23 | 11.01 | 0.51 | 0.62 | 2.71 | 5.50 | 0.10 | 0.02 | 170.07 | 279.78 | 474.23 | 181.87 |
| 98 | 2.76 | 11.89 | 0.52 | 0.59 | 3.45 | 5.64 | 0.13 | 0.03 | 148.90 | 210.54 | 422.90 | 161.90 |
| 99 | 2.07 | 10.03 | 0.52 | 0.57 | 2.59 | 4.68 | 0.10 | 0.02 | 160.08 | 232.25 | 487.30 | 200.13 |
| 100 | 2.80 | 11.08 | 0.57 | 0.65 | 3.81 | 5.78 | 0.11 | 0.02 | 159.88 | 275.72 | 514.63 | 224.77 |
| 101 | 2.41 | 11.78 | 0.57 | 0.69 | 3.31 | 6.53 | 0.11 | 0.02 | 192.69 | 291.14 | 369.83 | 128.70 |
| 102 | 2.45 | 10.29 | 0.54 | 0.67 | 3.19 | 5.54 | 0.10 | 0.02 | 160.29 | 269.92 | 412.33 | 167.80 |
| 103 | 2.35 | 11.19 | 0.66 | 0.77 | 3.74 | 6.89 | 0.12 | 0.02 | 215.66 | 297.04 | 590.77 | 242.13 |
| 104 | 2.67 | 12.33 | 0.46 | 0.52 | 2.99 | 5.45 | 0.12 | 0.02 | 126.40 | 225.58 | 501.40 | 196.10 |
| 105 | 2.18 | 9.52 | 0.55 | 0.64 | 2.91 | 4.89 | 0.10 | 0.02 | 159.12 | 242.70 | 459.63 | 189.00 |
| 106 | 2.76 | 12.34 | 0.59 | 0.73 | 3.91 | 7.19 | 0.14 | 0.03 | 177.22 | 253.71 | 531.67 | 227.87 |
| 107 | 2.52 | 11.59 | 0.66 | 0.75 | 4.04 | 7.03 | 0.13 | 0.03 | 190.86 | 269.63 | 541.13 | 219.47 |
| 108 | 2.71 | 11.83 | 0.66 | 0.71 | 4.45 | 6.82 | 0.13 | 0.03 | 177.43 | 232.40 | 416.17 | 175.67 |
| 109 | 2.22 | 10.44 | 0.56 | 0.56 | 2.98 | 4.66 | 0.10 | 0.02 | 174.52 | 233.43 | 409.40 | 145.00 |
| 110 | 2.07 | 10.07 | 0.50 | 0.55 | 2.48 | 4.50 | 0.11 | 0.02 | 146.56 | 204.16 | 568.90 | 230.50 |
| 111 | 2.58 | 10.71 | 0.63 | 0.69 | 3.87 | 5.93 | 0.10 | 0.02 | 210.30 | 289.37 | 559.13 | 216.87 |
| 112 | 2.49 | 11.53 | 0.65 | 0.76 | 3.86 | 7.10 | 0.12 | 0.02 | 186.59 | 285.40 | 456.87 | 184.43 |
| 113 | 2.59 | 12.11 | 0.61 | 0.69 | 3.82 | 6.69 | 0.14 | 0.03 | 160.76 | 237.98 | 579.20 | 242.63 |
| 114 | 2.76 | 11.64 | 0.62 | 0.62 | 4.14 | 5.75 | 0.13 | 0.03 | 172.72 | 221.29 | 529.50 | 224.07 |
| 115 | 2.37 | 12.02 | 0.56 | 0.61 | 3.21 | 5.80 | 0.12 | 0.02 | 162.29 | 234.55 | 384.90 | 131.57 |
| 116 | 2.30 | 10.15 | 0.65 | 0.68 | 3.58 | 5.54 | 0.10 | 0.02 | 191.43 | 270.78 | 436.07 | 169.03 |
| 117 | 2.52 | 11.16 | 0.61 | 0.65 | 3.71 | 5.87 | 0.11 | 0.02 | 182.62 | 263.21 | 442.97 | 179.97 |
| 118 | 2.58 | 10.46 | 0.66 | 0.70 | 4.09 | 5.86 | 0.11 | 0.02 | 202.74 | 271.69 | 450.20 | 180.80 |
| 119 | 2.41 | 11.28 | 0.57 | 0.57 | 3.47 | 4.51 | 0.11 | 0.02 | 201.40 | 206.53 | 441.70 | 178.03 |
| 120 | 2.89 | 11.66 | 0.56 | 0.69 | 3.91 | 6.57 | 0.14 | 0.03 | 140.51 | 238.89 | 356.87 | 137.23 |
| 121 | 2.83 | 12.51 | 0.60 | 0.69 | 4.09 | 6.95 | 0.13 | 0.03 | 174.71 | 257.80 | 228.60 | 76.33 |
| 122 | 2.62 | 12.05 | 0.61 | 0.72 | 3.90 | 7.05 | 0.14 | 0.03 | 174.29 | 253.99 | 556.80 | 225.93 |
| 123 | 2.62 | 11.33 | 0.53 | 0.66 | 3.34 | 5.91 | 0.13 | 0.03 | 139.92 | 213.90 | 404.00 | 171.57 |
| 124 | 2.76 | 12.10 | 0.62 | 0.67 | 4.10 | 6.51 | 0.14 | 0.03 | 170.25 | 235.83 | 503.97 | 207.10 |
| 125 | 2.40 | 12.97 | 0.61 | 0.71 | 3.53 | 7.47 | 0.16 | 0.03 | 163.21 | 239.29 | 528.33 | 207.17 |
| 126 | 2.76 | 12.87 | 0.65 | 0.73 | 4.38 | 7.57 | 0.15 | 0.03 | 203.46 | 258.28 | 581.07 | 233.10 |
| 127 | 2.76 | 12.93 | 0.70 | 0.75 | 4.64 | 7.75 | 0.14 | 0.03 | 193.68 | 283.72 | 494.10 | 197.03 |
| 128 | 2.29 | 11.13 | 0.64 | 0.74 | 3.55 | 6.59 | 0.13 | 0.03 | 182.37 | 263.40 | 456.97 | 172.80 |
| 129 | 2.63 | 10.86 | 0.54 | 0.63 | 3.49 | 5.53 | 0.11 | 0.02 | 171.79 | 255.59 | 529.70 | 209.70 |
| 130 | 2.48 | 11.34 | 0.64 | 0.75 | 3.84 | 6.83 | 0.12 | 0.02 | 196.09 | 290.41 | 531.30 | 217.33 |
| 131 | 2.57 | 11.69 | 0.70 | 0.76 | 4.35 | 7.16 | 0.13 | 0.03 | 200.68 | 267.64 | 472.60 | 185.73 |
| 132 | 2.36 | 10.95 | 0.69 | 0.71 | 3.92 | 6.30 | 0.13 | 0.03 | 204.16 | 239.36 | 432.43 | 177.83 |
| 133 | 2.39 | 9.87 | 0.59 | 0.63 | 3.35 | 5.06 | 0.11 | 0.02 | 182.33 | 234.41 | 611.10 | 249.50 |
| 134 | 3.36 | 14.37 | 0.73 | 0.86 | 5.83 | 9.90 | 0.19 | 0.04 | 168.96 | 263.39 | 449.07 | 162.57 |
| 135 | 2.43 | 10.08 | 0.56 | 0.62 | 3.23 | 5.01 | 0.10 | 0.02 | 168.07 | 247.32 | 550.97 | 216.83 |
| 136 | 2.81 | 13.35 | 0.61 | 0.73 | 4.11 | 7.80 | 0.15 | 0.03 | 175.53 | 264.31 | 492.53 | 194.20 |
| 137 | 2.68 | 12.11 | 0.55 | 0.65 | 3.57 | 6.26 | 0.13 | 0.03 | 167.46 | 241.53 | 345.90 | 133.67 |
| 138 | 2.64 | 10.57 | 0.52 | 0.63 | 3.32 | 5.33 | 0.11 | 0.02 | 162.92 | 252.99 | 572.67 | 243.03 |
| 139 | 2.24 | 10.58 | 0.56 | 0.60 | 3.01 | 5.06 | 0.13 | 0.03 | 140.24 | 202.40 | 464.70 | 192.50 |
| 140 | 2.77 | 11.66 | 0.78 | 0.84 | 5.23 | 7.86 | 0.13 | 0.03 | 221.30 | 310.76 | 621.00 | 254.20 |
| 141 | 2.59 | 11.73 | 0.57 | 0.60 | 3.52 | 5.55 | 0.12 | 0.02 | 166.93 | 231.78 | 589.53 | 241.73 |
| 142 | 2.54 | 11.93 | 0.66 | 0.66 | 4.07 | 6.36 | 0.12 | 0.02 | 198.03 | 271.42 | 528.97 | 216.33 |
| 143 | 2.46 | 11.24 | 0.69 | 0.69 | 4.06 | 6.24 | 0.13 | 0.03 | 153.09 | 248.68 | 585.63 | 254.73 |
| 144 | 2.23 | 10.98 | 0.57 | 0.68 | 3.09 | 6.07 | 0.11 | 0.02 | 185.59 | 289.03 | 491.43 | 196.13 |
| 145 | 2.59 | 11.30 | 0.59 | 0.71 | 3.71 | 6.47 | 0.12 | 0.02 | 163.73 | 267.29 | 563.20 | 226.57 |
| 146 | 2.59 | 13.04 | 0.76 | 0.68 | 4.73 | 7.10 | 0.13 | 0.03 | 226.91 | 268.39 | 442.60 | 176.40 |
| 147 | 2.06 | 10.47 | 0.59 | 0.57 | 2.91 | 4.83 | 0.12 | 0.02 | 183.57 | 207.42 | 386.03 | 149.10 |
| 148 | 2.87 | 13.70 | 0.66 | 0.66 | 4.57 | 7.24 | 0.15 | 0.03 | 189.61 | 234.25 | 505.80 | 222.40 |
| 149 | 2.65 | 12.28 | 0.55 | 0.69 | 3.82 | 7.12 | 0.13 | 0.03 | 178.47 | 226.82 | 483.70 | 198.37 |
| 150 | 2.48 | 12.80 | 0.64 | 0.69 | 3.84 | 7.07 | 0.13 | 0.03 | 197.29 | 287.99 | 505.27 | 208.57 |
| 151 | 2.44 | 10.82 | 0.68 | 0.64 | 4.01 | 5.63 | 0.12 | 0.02 | 187.16 | 243.74 | 604.10 | 243.80 |
| 152 | 2.27 | 10.55 | 0.53 | 0.63 | 2.91 | 5.43 | 0.10 | 0.02 | 169.74 | 256.86 | 330.57 | 115.27 |
| 153 | 2.75 | 12.25 | 0.68 | 0.70 | 4.46 | 6.89 | 0.14 | 0.03 | 182.49 | 251.94 | 481.10 | 199.07 |
| 154 | 2.44 | 11.41 | 0.62 | 0.70 | 3.64 | 6.52 | 0.12 | 0.02 | 181.71 | 277.80 | 602.73 | 245.53 |
| 155 | 2.53 | 11.95 | 0.60 | 0.72 | 3.69 | 6.94 | 0.12 | 0.02 | 180.78 | 284.36 | 598.60 | 251.57 |
| 156 | 2.40 | 12.14 | 0.53 | 0.63 | 3.07 | 6.11 | 0.12 | 0.02 | 147.47 | 244.76 | 482.40 | 197.73 |
| 157 | 2.68 | 12.27 | 0.67 | 0.69 | 4.29 | 6.83 | 0.14 | 0.03 | 195.96 | 241.77 | 631.73 | 272.50 |
| 158 | 2.79 | 13.95 | 0.60 | 0.72 | 4.07 | 8.09 | 0.15 | 0.03 | 168.00 | 265.58 | 544.93 | 213.60 |
| 159 | 2.92 | 13.43 | 0.70 | 0.79 | 4.96 | 8.47 | 0.15 | 0.03 | 203.62 | 292.05 | 556.30 | 219.93 |
| 160 | 2.62 | 11.88 | 0.60 | 0.71 | 3.80 | 7.09 | 0.11 | 0.03 | 168.40 | 265.48 | 570.33 | 199.40 |
| 161 | 2.92 | 11.28 | 0.56 | 0.70 | 3.93 | 6.35 | 0.12 | 0.02 | 156.68 | 259.77 | 482.50 | 187.00 |
| 162 | 2.56 | 11.12 | 0.60 | 0.57 | 3.69 | 5.10 | 0.12 | 0.02 | 177.02 | 216.42 | 613.40 | 245.00 |
| 163 | 2.39 | 10.89 | 0.52 | 0.65 | 3.03 | 5.62 | 0.12 | 0.02 | 159.46 | 234.04 | 538.57 | 221.43 |
| 164 | 2.99 | 12.31 | 0.54 | 0.70 | 3.92 | 7.02 | 0.14 | 0.03 | 169.16 | 239.95 | 520.97 | 209.27 |
| 165 | 2.70 | 11.47 | 0.61 | 0.73 | 3.95 | 6.83 | 0.14 | 0.03 | 163.96 | 235.89 | 509.63 | 198.10 |
| 166 | 2.33 | 10.61 | 0.59 | 0.69 | 3.30 | 5.81 | 0.12 | 0.02 | 180.58 | 255.30 | 526.67 | 210.97 |
| 167 | 2.25 | 12.03 | 0.61 | 0.69 | 3.30 | 6.73 | 0.12 | 0.02 | 208.62 | 276.33 | 436.00 | 179.20 |
| 168 | 2.35 | 8.42 | 0.50 | 0.54 | 2.82 | 3.64 | 0.08 | 0.02 | 127.42 | 224.67 | 349.50 | 102.00 |
| 169 | 2.53 | 11.58 | 0.64 | 0.74 | 3.91 | 6.90 | 0.13 | 0.03 | 178.81 | 270.56 | 489.57 | 186.23 |
| 170 | 2.35 | 10.79 | 0.54 | 0.62 | 3.05 | 5.42 | 0.11 | 0.02 | 162.31 | 253.21 | 424.70 | 180.40 |
| 171 | 2.67 | 11.25 | 0.53 | 0.59 | 3.39 | 5.31 | 0.11 | 0.02 | 152.75 | 250.88 | 654.03 | 276.73 |
| 172 | 2.41 | 10.96 | 0.48 | 0.58 | 2.79 | 5.16 | 0.11 | 0.02 | 155.75 | 232.19 | 455.03 | 181.13 |
| 173 | 2.63 | 12.61 | 0.69 | 0.71 | 4.40 | 7.19 | 0.14 | 0.03 | 190.67 | 260.58 | 644.20 | 258.77 |
| 174 | 2.20 | 9.80 | 0.69 | 0.67 | 3.67 | 5.34 | 0.10 | 0.02 | 192.91 | 263.56 | 238.70 | 82.60 |
| 175 | 2.16 | 9.67 | 0.56 | 0.66 | 2.96 | 5.21 | 0.09 | 0.02 | 184.45 | 290.51 | 589.27 | 223.47 |
| 176 | 2.34 | 11.73 | 0.53 | 0.54 | 3.02 | 5.17 | 0.12 | 0.02 | 156.12 | 215.28 | 528.40 | 205.00 |
| 177 | 2.73 | 11.76 | 0.62 | 0.70 | 4.15 | 6.69 | 0.13 | 0.03 | 191.90 | 265.05 | 481.63 | 202.10 |
| 178 | 1.92 | 9.59 | 0.50 | 0.63 | 2.32 | 4.86 | 0.09 | 0.02 | 170.20 | 271.54 | 499.90 | 199.77 |
| 179 | 2.18 | 11.37 | 0.64 | 0.65 | 3.36 | 5.86 | 0.11 | 0.02 | 202.80 | 264.95 | 599.90 | 249.70 |
| 180 | 2.18 | 11.72 | 0.55 | 0.65 | 2.91 | 6.12 | 0.12 | 0.02 | 161.25 | 247.65 | 444.03 | 175.67 |
| 181 | 1.91 | 10.00 | 0.48 | 0.66 | 2.21 | 5.23 | 0.11 | 0.02 | 151.75 | 241.25 | 493.10 | 181.00 |
| 182 | 2.41 | 11.47 | 0.62 | 0.75 | 3.57 | 6.86 | 0.12 | 0.02 | 196.19 | 293.37 | 528.17 | 210.20 |
| 183 | 2.66 | 12.13 | 0.61 | 0.70 | 3.93 | 6.82 | 0.12 | 0.02 | 182.17 | 278.25 | 505.47 | 205.33 |
| 184 | 2.38 | 9.84 | 0.59 | 0.70 | 3.35 | 5.55 | 0.11 | 0.02 | 159.73 | 262.10 | 557.17 | 231.23 |
| 185 | 2.30 | 10.49 | 0.56 | 0.65 | 3.14 | 5.52 | 0.11 | 0.02 | 169.35 | 251.44 | 434.87 | 180.37 |
| LSD 5% | 0.80 | 1.74 | 0.12 | 0.37 | 0.91 | 1.69 | 0.03 | N.S | 38.57 | 54.02 | 176.59 | 82.13 |
| LSD 1% | 1.06 | 2.29 | 0.15 | 0.49 | 1.20 | 2.23 | 0.03 | N.S | 50.78 | 71.13 | 232.52 | 108.14 |

## References

Argyris, J., Truco, M.J., Ochoa, O., Knapp, S.J., Still, D.W., Lenssen, G.M., Schut, J.W., Michelmore, R.W., & Bradford, K.J. (2005). Quantitative trait loci associated with seed and seedling traits in Lactuca. Theor Appl Genet 111: 1365–1376.

Botwright, T.L., Condon, A.G., Rebetzke, G.J., & Richards, R.A. (2002). Field evaluation of early vigour for genetic improvement of grain yield in wheat. Aust. J. Agric. Res. 53: 1137– 1145.

Broughton, S., Zhou, G., Teakle, N.L., Matsuda, R., Zhou, M., O’Leary, R.A., Colmer, T.D., & Li, C. (2015). Waterlogging tolerance is associated with root porosity in barley (Hordeum vulgare L.). Mol. Breed. 35.

Burton, G. & Devane, E. (1953). Estimating Heritability in Tall Fescue (Festuca Arundinacea) from Replicated Clonal Material. Agron. J. 45: 478.

Burton, G.W. (1952). Quantitative inheritance in grasses. In Proceedings of 6th International Grassland Congress, pp. 277–283.

Chen, G., Krugman, T., Fahima, T., Chen, K., Hu, Y., Röder, M., Nevo, E., & Korol, A. (2010). Chromosomal regions controlling seedling drought resistance in Israeli wild barley, Hordeum spontaneum C. Koch. Genet. Resour. Crop Evol. 57: 85–99.

Elakhdar, A., EL-Sattar, M.A., Amer, K., Rady, A., & Kumamaru, T. (2016). Population structure and marker–trait association of salt tolerance in barley (Hordeum vulgare L.). C. R. Biol. 339: 454–461.

Flores, O. et al. (2014). An evolutionary perspective on leaf economics: phylogenetics of leaf mass per area in vascular plants. Ecol. Evol. 4: 2799–2811.

Gawroński, P., Pawełkowicz, M., Tofil, K., Uszyński, G., Sharifova, S., Ahluwalia, S., Tyrka, M., Wędzony, M., Kilian, A., & Bolibok-Brągoszewska, H.. DArT Markers Effectively Target Gene Space in the Rye Genome. Front. Plant Sci.

Heneen, W.K. (2011). Cytogenetics and Molecular Cytogenetics of Barley: A Model Cereal Crop with a Large Genome. In Barley (Wiley-Blackwell: Oxford, UK), pp. 112–121.

Herwaarden, A.F. van, Farquhar, G.D., Angus, J.F., Richards, R.A., & Howe, G.N. (1998). “Haying-off”, the negative grain yield response of dryland wheat to nitrogen fertiliser. I. Biomass, grain yield, and water use. Aust. J. Agric. Res. 49: 1067.

Hoffmann, A., Maurer, A., & Pillen, K. (2012). Detection of nitrogen deficiency QTL in juvenile wild barley introgression lines growing in a hydroponic system. BMC Genet. 13.

Hori, K., Sato, K., & Takeda, K. (2007). Detection of seed dormancy QTL in multiple mapping populations derived from crosses involving novel barley germplasm. Theor. Appl. Genet. 115: 869–876.

Johnson, H.W., Robinson, H.F., & Comstock, R.E. (1955). Estimates of Genetic and Environmental Variability in Soybeans. Agron. J. 47: 314.

Katsvairo, T.W., Cox, W.J., Van Es, H.M., & Condon, A.G. (2003). Spatial Growth and Nitrogen Uptake Variability of Corn at Two Nitrogen Levels. Agron. J. 95: 1000.

Li, X. M., Chen, X. M., Xiao, Y. G., Xia, X. C., Wang, D. S., He, Z. H., & Wang, H. J. (2014). Identification of QTLs for seedling vigor in winter wheat. Euphytica 198: 199–209.

Liu, L., Sun, G., Ren, X., Li, C., & Sun, D. (2015). Identification of QTL underlying physiological and morphological traits of flag leaf in barley. BMC Genet. 16: 29.

López-Castañeda, C., Richards, R.A., Farquhar, G.D., & Williamson, R.E. (1996). Seed and seedling characteristics contributing to variation in early vigor among temperate cereals. Crop Sci. 36: 1257–1266.

Lu, X. L., Niu, A. L., Cai, H. Y., Zhao, Y., Liu, J. W., Zhu, Y. G., & Zhang, Z. H. (2007). Genetic dissection of seedling and early vigor in a recombinant inbred line population of rice. Plant Sci. 172: 212–220.

Mahender, A., Anandan, A., & Pradhan, S.K. (2015). Early seedling vigour, an imperative trait for direct-seeded rice: an overview on physio-morphological parameters and molecular markers. Planta 241: 1027–1050.

Mangin, B., Goffinet, B., & Rebai, A. (1994). Constructing confidence intervals for QTL location. Genetics 138: 1301–1308.

Mansour, E., Casas, A.M., Gracia, M.P., Molina-Cano, J.L., Moralejo, M., Cattivelli, L., Thomas, W.T.B., & Igartua, E. (2014). Quantitative trait loci for agronomic traits in an elite barley population for Mediterranean conditions. Mol. Breed. 33: 249–265.

Mikołajczak, K. et al. (2016). Quantitative Trait Loci for Yield and Yield-Related Traits in Spring Barley Populations Derived from Crosses between European and Syrian Cultivars. PLoS One 11: e0155938.

Pérez-Harguindeguy, N. et al. (2013). New handbook for standardised measurement of plant functional traits worldwide. Aust. J. Bot. 61: 167.

Poorter, H., Niklas, K.J., Reich, P.B., Oleksyn, J., Poot, P., & Mommer, L. (2012). Biomass allocation to leaves, stems and roots: meta-analyses of interspecific variation and environmental control. New Phytol. 193: 30–50.

Qin, D., Dong, J., Xu, F., Guo, G., Ge, S., Xu, Q., Xu, Y., & Li, M. (2015). Characterization and fine mapping of a novel barley Stage Green-Revertible Albino Gene (HvSGRA) by Bulked Segregant Analysis based on SSR assay and Specific Length Amplified Fragment Sequencing. BMC Genomics 16: 838.

Rajjou, L., Duval, M., Gallardo, K., Catusse, J., Bally, J., Job, C., & Job, D. (2012). Seed Germination and Vigor. Annu. Rev. Plant Biol. 63: 507–533.

Rajjou, L., Lovigny, Y., Groot, S.P.C., Belghazi, M., Job, C., & Job, D. (2008). Proteome-wide characterization of seed aging in Arabidopsis: a comparison between artificial and natural aging protocols. Plant Physiol. 148: 620–41.

Rebetzke, G.J., Botwright, T.L., Moore, C.S., Richards, R.A., & Condon, A.G. (2004). Genotypic variation in specific leaf area for genetic improvement of early vigour in wheat. F. Crop. Res. 88: 179–189.

Rebetzke, G.J., Van Herwaarden, A.F., Jenkins, C., Weiss, M., Lewis, D., Ruuska, S., Tabe, L., Fettell, N.A., & Richards, R.A. (2008). Quantitative trait loci for water-soluble carbohydrates and associations with agronomic traits in wheat. Aust. J. Agric. Res. 59: 891– 905.

Rebetzke, G.J. & Richards, R.A. (1999). Genetic Improvement of Early Vigour in Wheat. Aust. J. Agric. Res. 50: 291–301.

Richards, R.A. (2000). Selectable traits to increase crop photosynthesis and yield of grain crops. J. Exp. Bot. 51: 447–458.

Richards, R.A. & Lukacs, Z. (2002). Seedling vigour in wheat - Sources of variation for genetic and agronomic improvement. Aust. J. Agric. Res. 53: 41–50.

Richards, R.A., Rebetzke, G.J., Condon, A.G., and Van Herwaarden, A.F. (2002). Breeding opportunities for increasing the efficiency of water use and crop yield in temperate cereals. Crop Sci. 42: 111–121.

Rodríguez-Suárez, C., Giménez, M.J., Gutiérrez, N., Ávila, C.M., Machado, A., Huttner, E., Ramírez, M.C., Martín, A.C., Castillo, A., Kilian, A., Martín, A., & Atienza, S.G. (2012). Development of wild barley (Hordeum chilense)-derived DArT markers and their use into genetic and physical mapping. Theor. Appl. Genet. 124: 713–722.

Sandve, S.R., Kosmala, A., Rudi, H., Fjellheim, S., Rapacz, M., Yamada, T., & Rognli, O.A. (2011). Molecular mechanisms underlying frost tolerance in perennial grasses adapted to cold climates. Plant Sci.

Sbei, H., Sato, K., Shehzad, T., Harrabi, M., & Okuno, K. (2014). Detection of QTLs for salt tolerance in Asian barley (Hordeum vulgare L.) by association analysis with SNP markers. Breed. Sci. 64: 378–388.

Singh, U.M., Yadav, S., Dixit, S., Ramayya, P.J., Devi, M.N., Raman, K.A., & Kumar, A. (2017). QTL Hotspots for Early Vigor and Related Traits under Dry Direct-Seeded System in Rice (Oryza sativa L.). Front. Plant Sci. 8: 286

ter Steege, M.W., den Ouden, F.M., Lambers, H., Stam, P., & Peeters, A.J.M. (2005). Genetic and Physiological Architecture of Early Vigor in Aegilops tauschii, the D-Genome Donor of Hexaploid Wheat. A Quantitative Trait Loci Analysis. Plant Physiol. 139: 1078– 1094.

Sundgren, T.K., Uhlen, A.K., Lillemo, M., Briese, C., & Wojciechowski, T. (2018). Rapid seedling establishment and a narrow root stele promotes waterlogging tolerance in spring wheat. J. Plant Physiol. 227: 45–55.

Tian, M., Yu, G., He, N., & Hou, J. (2016). Leaf morphological and anatomical traits from tropical to temperate coniferous forests: Mechanisms and influencing factors. Sci. Rep. 6: 19703.

Tyrka, M., Bednarek, P.T., Kilian, A., W dzony, M., Hura, T., & Bauer, E. (2011). Genetic map of triticale compiling DArT, SSR, and AFLP markers. Genome 54: 391–401.

Ullmannová, K., Středa, T., & Chloupek, O. ich (2013). Use of barley seed vigour to discriminate drought and cold tolerance in crop years with high seed vigour and low trait variation. Plant Breed. 132: 295–298.

Wang, J., Sun, G., Ren, X., Li, C., Liu, L., Wang, Q., Du, B., & Sun, D. (2016). QTL underlying some agronomic traits in barley detected by SNP markers. BMC Genet. 17: 103.

Wang, Q., Sun, G., Ren, X., Wang, J., Du, B., Li, C., & Sun, D. (2017). Detection of QTLs for seedling characteristics in barley (Hordeum vulgare L.) grown under hydroponic culture condition. BMC Genet. 18.

Wang, S., Basten, C.J., & Zeng, Z.B. (2011). Windows QTL Cartographer 2.5. J. Infect. Dis. 204 Suppl: 198–199.

Wen, D., Hou, H., Meng, A., Meng, J., Xie, L., & Zhang, C. (2018). Rapid evaluation of seed vigor by the absolute content of protein in seed within the same crop. Sci. Rep. 8: 5569.

Wójcik-Jagła, M., Rapacz, M., Tyrka, M., Kościelniak, J., Crissy, K., & Żmuda, K. (2013). Comparative QTL analysis of early short-time drought tolerance in Polish fodder and malting spring barleys. Theor. Appl. Genet. 126: 3021–3034.

Yin, X., Kropff, M.J., & Stam, P. (1999). The role of ecophysiological models in QTL analysis: the example of specific leaf area in barley. Heredity (Edinb). 82: 415–421.

Zhou, G., Panozzo, J., Zhang, X. qi, Cakir, M., Harasymow, S., & Li, C. (2016). QTL mapping reveals genetic architectures of malting quality between Australian and Canadian malting barley (Hordeum vulgare L.). Mol. Breed. 36.

Zhu, C., Gore, M., Buckler, E.S., & Yu, J. (2008). Status and Prospects of Association Mapping in Plants. Plant Genome J. 1: 5.

