## Supplemental for "Major QTLs for seedling traits in barley using a DArT-based linkage map"

**Ch. 1**

bPb-7963

tgcagtaccatactttcctccaaatagttcttcttttgcatgtccttgaaattccttgtttccaaaagacactaatgttg

gaataaaccacgatgcgtggccttgtccggctgggactcggtgcaatcacctaatagtactaggattaagaattggtaac

cgtataagtttacttgtttgatgatagctgctttatatacgtaggtatcctgtaatattatgaaccaaaatcaaagcaat

acaaagcacaggaccgacacagccgtgttcgccatcacctcgagttttcgcgtgtatgttctgttgtgaatgaattagcg

ccggcgagatgatcgctaacttagctgattaagctagctactcgtgtgtgggtgtccagcagcactcgatccacctatgt

atctgcttgCttagtactactacgtgtgtatttgattagtgtggtaatatatTgcacgtacgtgtcgactcgtcaaggaa

ccgacgtgcaacatctgatcgggtctgtgacaacaattggtatctagaacttgatttgtttgcgtgattttcggaaagta

atcgaagaatcatgtcaAacccgggcaaggagattgtcgtggcgggagcgggagcaccgatgccgatggtgggggtgtct

caattcccccgactggatcgagaggactacgcactatgagcgatgaacatggaggtcgcgatggagggcgctgagttctg

gtaagctatcgatcccggtggagcccagtacacgaagggtgcggcgaagtaccggacgggtcgtcaggcgttaacggtga

tctactccgccatgccgaaggatgtgctgca

bPb-4293

TGCAGGACGAGGGGCACCGGGAGGGGAAGGGGGTGGAGCCCCCGCAACTGGTCGCGATGGCATAATATGTCCGCCGTCTC

CGAATGTTGGCAAGTATGTCGTCGGATGATCCCATGGCGCTTGAAGCCACGGCCACGGGAACATATGTCCACCATCACAT

GGCGGATGCATCGGCTGGGGAGGGCACCATTGTGGTTGCATAGGTTGGGGCATTGCCGGCGGTAGCAACCATGATGGTGG

ATGTGGCCAGCCCCCACCCCCAGCCCCCCCTTGTGATTCTAAGTCTTCCGCTGATGAGGGGTCTTCATTGTGATTGATCC

AATCACCAGTTCCATTGCCCGGATCTTCTGTATCACTGGATTCCTCTTTTTCATCTGGACCTTCTTTGCCAATCTCAAAC

GGGCCAACATGCTCTACCATAATCCATACGATGGTAATTAGGTCACCGCCACGGCCGAGTTGCCTCGCATGGGATTCTCT

GCTGCATCGGGTGGCTGCATACAACAGCTTATCCACCCAAATCTTGAAGACGAGCTCCAAAAGTTTCGGCATCTTCAAAC

CATCAGCCTTGAGCAGCTCCAGAGCAATATTAGCACCGTCTGAGACCAGACGAGTTCTACTCTTACCTAAACCCCACTCG

TCGTCTCCCTTCTTGTTCTTTGCCAGCATCCTTGCAAGCTTCTCCTCTGCA

bPb-7697

tgcagttcttggatgctttgcaacttaccaaaagtttttggtattgatcccgatatttggttctgaaacaagcctaaata

ttcgaggttcatcagatcgccaatttcctcggggatggaaccggttatttgattttcaaagaggtatagtactactagct

tggtaacatttcctaagctgtcaggaatagagccagatatttggttgatggacaagtctaaattctggagattcattaga

ttgccaaactctctggggatggaaccagatatcttattttcaaagaggtgtagtagtaatagcttggtaatatttcctaa

gctgtcaggaatagagccagatatttggttgtcagccaagtctaacttctggagattcagcaagtagCccaattctgaag

gtattgtgcctgtgatatgatttgtatagagaagaagttcattgagcatagtgaggttgcctacttctgccggtattgaa

cctgtgatttgatttctatagagaccaagttggtttagcatagtgaggttgcctattgctgggggtattgaacctgtgat

ttgattttcaaagagaaaaaattggttcaactttgtaagattggttatgaagactggaattggacctgacaaatcatttc

cagcaagatgaagaTtttgcaaatggactagcctgcctagttcttggggtataggccctgaaagttgattatgaaacacg

tacaaagtatttagttgggtcaGaTttccaagggtttttggtatcatgccgcctaaggtgttgttgcttagctgtagaag

ttgtaggttgacaagccttccaatctcttcaggaatgggacctgataccatggtttggtgaatgataagatcagttaaca

ttgttaggttacacagagacacagggatatgtcctgtgagtttgttaaatgagagaccaagttgggtgagactctgca

bPb-5064

tacgggtcgatcaagttagggttcgcttgtccttgtttggttctcggccatggcggcgctaaagggcctcgtcgccagca

cccacacggcaccatgctaccgagacatctcgtgcagggctgccgaacgacatcgtcgccggtggccagtcggactgctc

gaccatggCagccgctcgccatgttgttggccgggcgcactcctcgacggtggctattcatgcgccaacatgtcatcgcg

ggcagcgggccaactcctCgtggacgaacggacgcgcaccaagacccttcgaagtgctccattttgtcgcacttgctcaa

gccggcgccggtgtactccatgagaccatgacacttacgtgatcaggctgattgtaagacgcatgcttacatattcaGca

tatacatgtgcattgTctagtttggttgatcgattcctttagtctcatgctaGcccttgattaGattaGTatgatGcttg

GCcaaactgttatatcaTtaGttcaatctgtAaatacttaaGaTaatatcttttttttttgcaggtgaagataatattgt

tttcaaaacactattcggcgataggattttaatgaaggaactagtacaagcaggctccagtgatgacttggagcacttca

aaaaggaggcaactgca

**bPb-1322**

bPb-8609

TGCAGCAGATGTATCAAAGGGCAGTGAGCATATGTGAATGGTAGAACAGAAATGAACTAACAAAAGAGAATACGGTGTAG

CCATCTCAGCAACATGCACCACTAGGAGTAAAGAAGGGAGGCTTCATATATTATTGAGAATTCGGCATGAATGCATTTAT

CATGAAAGACTTGCTACATTAGTTTGCCTCGTCAGGAAACTTATTTGTTCAGTACCACAAAGTTGGAGCTACCAATGACA

TGTTTGCTAGATCAGTTTGTCGTCACGCAACATGGGTCAGAGCATACCTCATAAAGCTTCGTCAAGTAAATCCTTACGAT

TGGTCATTCCAATAAGCAACACGTTATTCAACGCCTCACACCGTCTATCTGTAAATTTGA

bPb-5297

ccnctagtaacggccgccagtgtgctggaattcgccctttggatccagtgcagcatgccagccagataaaaatggtttat

ttcctgtacacattgctgccatgatgaatagtagaattgccatccatgttctgctcacactataccctcgttgcatcgga

tttcctgataaccagggaagatccttccttcatattgctgcccagaacaagaactcccTtgtagtaagatatgcttgcag

tgtatcgtGtcgggattttggatcgataatgaagtcgataatgaacgctcaggacgacgatggcaacaccgcactacatc

tagctgtggatgctggggatctatcttctttCtgtgcactgctggggaaccgagaagtattattaaatatcagaaataac

aagaatcaaactccactggatttggcatggagcaagaggaaaggttttgcgtacggatgggtattaatctatatttcctc

tctcatttcctcatgttagttcgtcatatagctagaagatgtatcccagtatcccacttcatttttttcagtaaatatat

atcccgcatcttgatgattttcttgcagaatccggaacatgtgatgtacagaactcttgtacgtgccggctctagtcatg

ctagcttttggagagatcgtcttcaacaactttgcaatttagtgccactagaaagtcaagaggatgaagggaaagatcca

gcaaaagaaaaaaaagaggatgaagagaaagaagagtcagctaaagtgacagattcaactcggacacttggcattggctc

ggtactcatagcatcagtgacattcactgcaaccttcaccgtacctgcactggntccatcaagggcgaattctgcanata

tccatnnnncctggcnnnnnntcgagcatgcatctagagggcc

bPb-9718

tgcagcgatagcccttacgggagctatggcggcaagacatcttttacgtggtactggtccggggaggaggatgaggatga

ccttcctaatgggttccaatggagggacgagcctcgaccgaacaaatcaaaggaaagagtttggaacgagagcgatgtgg

atgaggaagaggcccctcgtcgcgatgatctgaaaagccataggatatcccttggcttgccagccttgggtcccttaaag

cttgatcatatcaagtccgcgtaagtctttgagcatctgcatagcatttgcgaggattctgataatcgggctcctgattg

gtgttctctttctcgatgctacgtttctgca

**bPb-3309**

bPb-3249

tgcagaggtgttcgttagctatttgcagttagagagtgctcaatataagacctgcatgccttttctgaatatagcagatt

ttttttctcagcaacgaggtaaatgaagtgcactgccaaataaagcatttaaactaattttagtgctgaccacactgagc

tggtggttagtaccagttgaaggttgtgtttagacttatatccttgggacagaatggaaggcaaatggaaaacaaatttt

gggttaaaccggacagagggaaaagaaaaatgaacagcccacaagacaatgagtatctgttctttttttaggcaattggc

taagaattgtgcattgtctacaactgcaagcaaagcaaagactatactaagagtctcgcatcgtctccggctgcaagcaa

agcaaagggaagctagcagaaacttggcatgccctgca

**bPb-1204**

**bPb-2664**

bPb-5072

tgcagaaatgcacgcacaaagaaatcaagttacatgctaatccaaacatcagctctagctcttgtcaaacggagttcatt

catagaagagtcttacaaattgaaggcagacatcacatttcgggtggtgtagctccttgtagcacaacttgtggtatggc

tctgcgcctagcaaggtgaactggaaacacaaagatagagaaaacgggtgacggtcactttctatttcaatggatctgta

gctcaaggcttaaagggagtgtaggatggtcacctcagtctcacggatagggtgggtgcaggaagagcaacggaagcact

gaggatgccagtacatccccatgcagctcaagtaatggccatgtccaacctcatgcttgcagccaccacagaccctgca

bPb-2055

tgcagcataccaacatgataagaatggtttgttacctgtacatgtcgctgccttgatgcataggaaagttgccatcctta

ttctcctcgaaagatgcggtggttgcctcgctcttcctgataagcagggtagatcctttcttcacattgctgtccagagt

gaggcctatggtatagtatgatatgcttgtcaagaaccngtttttgggccgatactcaacgctcgggacaatgatggcaa

caccgcattacacctagctgtggaggttgaggatttactcatcgtctgttatctcttgcggaatccaaaagtattattaa

atgtaagaaacaacaaggatcaaactccgctggatctagcaaggaacaagactcatctcacagatttttcctatggactg

gtattaaatctagcttccgttctcatgcatgcttcatgctagttgatcttataaatggaaagataaaactatatttcttc

attattgtgttttcagaatccggagaacgcaatatacgagacactcagggatgtcggagctaggcatggtagcttctgga

gtgatcatgttcaacaactctgcattcagaatcagttgtcaaacctaAagggtgaagagAatcaAtcagaaGaagtggat

gattttgagatgcataAGAagaaAAaggatgaaGagaatcaAtcaGaagAagtggatgagttggaGaagcatacGgAaaa

aaaggataaaaagaaaaaggaagcgaaccgtaagaaaaaagagtatgaaaagaaaaaagaggtgaaccgtaagaaaaaag

aggaagaagagaaaaaagactcagatcaaatgaatgattcaacccggacacttggcattggctcggtactcatcacaacg

atgacattcggtgcaaccttcacagtacctgca

bPb-0487

ccagtGcagggcatgccaagtttctgctagcttccctttgCtttgcttgcagccggagacgatgcgagactCttagtata

gtctttgctttgcttgcagttgtagacaatgcacaattcttagccaattgcctaaaaaaagaacagatactcattgtctt

gtgggctgttcatttttcttttccctctgtccggtttaacccaaaatttgttttccatttgccttccattctgtcccaag

gatataagtctaaacacaaccttcaactggtactaaccaccagctcagtgtggtcagcactaaaattagtttaaatgctt

tatttggcagtgcacttcatttacctcgttgctgaGaaAaaaaatctgctatattcAgaaaaggcAtgcaggtcttatat

tgagcactctctaactgcAaaatagctAaCgaacacctctgcactggatccntca

bPb-8973

tgcagcataccaacatgataagaatggtttgttacctgtacaTgtcgctgccttgatgcataggaaagttgccatcctta

ttctcctcgaaagatgcggtggttgcctcgctCttcctgataagcagggtagatcctttcttcacattgctgtccagagt

gaggcctatggtatagtacgatatgcttgtcaagaaccagtttttgggccgatactcaacgctcgggacaatgatggcaa

caccgcattacacctagctgtggaggttgaggatttactcatcgtctgttatctcttgcggaatccaaaagtattattaa

atgtaagaaacaacaaggatcaaactccgctggatctagcaaggaacaagactcatctcacagatttttcctatggactg

gtattaaatctagcttccgttctcatgcatgcttcatgctagttgatcttataaacggaaagataaaactatatttcttc

attattgtgttttcagaatccggagaacgcaatatacgagacactcagggatgtcggagctaggcatggtagcttctgga

gtgatcatgttcaacaactctgcattcagaatcagttgtcaaacctaaagggtgaagagaatcaatcaGaagaagtggat

gaTtttgagatgcataagaagaaaaaggatgaagagaatcaatcagaAGAaGtggAtgagttggagaaGcatacggaaaa

aaaggataaaaAGaaaAAggaagcgaaccgtaagaaaaaagagtatgaaaagaaaaaagaggtgaaccggaagaaaaaag

aggaagaagagaaaaaagactcagatcaaatgaatgattcaacccggacacttggcattggctcggtactcatcacaacg

atgacattcggtgcaaccttcacagtacctgca

bPb-8345

tgcagggcatgccaagtttctgctagcttccctttgctttgcttgcagccggagacgatgcgagactcttagtatagtct

ttgctttgcttgcagttgtagacaatgcacaattcttagccaattgcctaaaaaaagaacagatactcattgtcttgtgg

gctgttcatttttcttttccctctgtccggCttaacccaaaatttgttttccatttgccttccattctgtcccaaggata

taagtctaaacacaaccttcaactggtactaaccaccagctcagtgtggtcagcactaaaattagtttaaatgctttatt

tggcagtgcacttcatttacctcgttgctgagaaaaaaaatctgctatattcagaaaaggcatgcaggtcttatattgag

cactctctaactgcaaatagctaacgaacacctctgca

bPb-3579

TGCAGGGCATGCCAAGTTTCTGCTAGCTTCCCTTTGCTTTGCTTGCAGCCGGAGACGATGCAAGACTCTTAGTATAGTCT

TTGCTTTGCTTGCAGTTGTAGACAATGCACAATTCTTAGCCAATTGCCTAAAAAAAGAACAGATACTCATTGTCTTGTGG

GCTGTTCATTTTTCTTTTCCCTCTGTCCGGTTTGACCCAAAATTTGTTTTCCATTTGCCTTCCATTCTGTCCCAAGGATA

TAAGTCTAAACACAACCTTCAACTGGTACTAACCACCAGCTCAGTGTGGTCAGCACTAAAATTAGTTTAAATGCTTTATT

TGGCAGTGCACTTCATTTACCTCGTTGCTGAGAAAAAAAATCTGCTATATTCAGAAAAGGCATGCAGGTCTTATATTGAG

CACTCTCTAACTGCAAATAGCTAACGAACACCTCTGCA

bPb-0374

tgcagaggtgttcgttagctatttgcagttagagagtgctcaatataagacctgcatgccttttctgaatatagcagatt

ttttttctcagcaacgaggtaaatgaagtgcactgccaaataaagcatttaaactaattttagtgctgaccacactgagc

tggtggttagtaccagttgaaggttgtgtttagacttatatccttgggacagaatggaaggcaaatggaaaacaaatttt

gggtcaaaccggacagagggaaaagaaaaatgaacagcccacaagacaatgagtatctgttctttttttaggcaattggc

taagaattgtgcattgtctacaactgcaagcaaagcaaagactatactaagagtcttgcatcgtctccggctgcaagcaa

agcaaagggaagctagcagaAActtggcatgccctgcactggatccntcaaggggcnaattctgcagatatccatcanac

ctgggcggcangctcgagcatgcatctngannnccaattcgccctatagtgagtcgtantacaattcactggccgtcgtt

ttacaacgtcgn

bPb-5201

tgcagatgtacaaaatgccgtcatcaaccatatgtgtgggcagcacggtcacatcatagtggcaaggcaagagcagcagg

ccagcacccagaaaacacagtagttccaccttccctgtctaatcgcggcaaacacggtcacagtcgtagaagtgatggat

aatttgatccttttaagaatgtattttgtctgatccctccctcgacattaccaaccggcaaggcagcttgTttttAtccg

agagattattagtctttgacgcaagttggtgattatttgaagagatgaacacagctagctgccgtcgatgatccatagct

gcggtagattatttttaaaaactcaatctaaaacctaatcccagtcggcactgctactgtgcttgtttatagtttttcaa

gcgaggccggatcttggacattgattttcttgcatcaccacttcaccagtatattacttgtatggcattagtcattagaa

accgtaactatacatgcatacgtaccggtgcagttgtgagcttgacgggcaaccgctccaaatgatgattagagggagag

gacgatgctgcactgctgcttccgcctctgtttgtcgcgtacttagcgtatgtacgtacgttcttcttcaagttaaaaac

acaatctcacagattcgggtcttggtctcttgcaataagcatattctaggaacacataccacgacacgacagatagcagt

tccaaaaaatatattaaaactattattctgacttcgtacttaacttgctaacgaactcaaactgattcgtctcttttcta

tatttcacctgaTttcggacgcccggctacaacccgctcggtggtggcagcctgca

bPb-9414

tgcagcataccaacataataagaatggtttgttacctgtacnngtcgctgccttgatgcataggaaagttgccatcctta

ttctcctcgaaagatgcggtggttgcctcgctcttcctgataagcagggtagatcctttcttcacattgctgtccagagt

gaggcctatggtatagtacgatatgcttgtCaagaaccagtttttgggccgatactcaacgctcgggacaatgatggcag

caccgcattacacctagctgtggaggttgaggatttactcatcgtctgttatctcttgcggaatCcaaaagtattattaa

atgtaagaaacaacaaggatcaaactccgctggatctagcaaggaacaagactcatctcacagatttttcctatggactg

gtattaaatctagcttccgttctcatgcatgcttcatgctagttgatcttataaatggaaagataaaactatatttcttc

attattgtgttttcagaatccggagaacgcaatatacGagacactcagggatgtcggagctaggcatggtagcttctgga

gtgatcatgttcaacaactctgcattcaGaatcagttgtcaaacctaaagggtgaagagaatcaatcagaagaagtggat

gattttgagatgcataagaagaaaaaggatgaagagaatcaatcagaagaagtggatgagttggagaagcatacggaaaa

aaaggataaaaagaaaaaggaagcgaaccgtaagaaaaaagagtatgaaaagaaaaaaggggtgaaccgtaagaaaaaag

aggaagaagagaaaaaagactcagatcaaatgaatgattcaacccggacacttggcattggctcggtactcatcacaacg

atgacattcggtgcaaccttcacagtacctgca

bPb-7221

TGCAGCCGTAGCCGCCGTCTTGTCTGCCTTCATGAGCAATTTCTTATCACATTATTTGCACTTTTGCAAAAAGTTCGGCG

AGACCATCCTCGTCTGTAACTAATTCTTCATCGTAGTACCTCCAAATGGTAGAGCATCTCCAACGAGTGTTTTGATTTGG

ATTAGACATGTGTGTGGAAAATGCGATGGAGTTTGAGTGTGAGAAATGAGAAAGTGAGCATCCTTTTTTATAGCAAAAAA

TGGCACGTTAGGCACATTAGACAACAATAATGCATAAAATAAAAAATGAGGTAATGTATCATCTGGGATTAATCTTGAGC

TTTATTCGTGTCTACCTCATTAAATCCTCCCATTCCTGGGAAGTAAAAGAATACAAAGCCACGCTCTAGAATTACAAAAG

GGAAATAAAATTACAAAAAGATAAGAAATTTACAAAAGAAAAAGAAAAGGACAACACTACTATTCTTCCCGTTCTTCTTC

TTCAAGGTTCATTTCTTCAAACATGGGATCTTCTGCA

**bPb-45000**

**bPb-26569**

bPb-6873

tgcagaaagttaatatcacgtattctgttaaaaactaaatcatttctgcaattctatagagcccacaaaagagcacatgc

tccaacacgaatatgtttggccaaataaacatccactccctgtaaaaggcaaatgattattatgattgttccttatttat

ctttcggcatcagctagtatgtaaattgaacatcattctttttggcaacagacgctagtgaggaaaatactctaaactgg

gagcaacggtacaacatcattcttggaattgccaagggaatactgtatcttcacgaggactcaatcccaaggataatcca

cagggaccttaaagctaataatattcttctagacgaggagatggatcctaaaatcgcagactttggattggcaaggctgc

tacaagaaggtcacacccatactcaaaccactagagctgctggaacactgtaagtcaaatgcctataattaggtttgcaa

aaacgaaaaagttacttgtaattatgatctatatcacttagtgaaggcccgtttccaagcctaatttcccatcctctgca

bPb-9410

tgcagatgtacaaaatgccgtcatcaaccatatgtgtgggcagcacggtcacatcatagtggcaaggcaAgagtagcagg

ccagcacccagaaaacacagtagttccaccttccctgtCtaatcgcggcaaacacggtcacagtCgtagaagtgatggat

aatttgatccttttaagaatgtattttgtctgAtccctccctcgacattaccaAccggcaaggcagcttgtttttatccg

agagattattagtctttgacgcaagttggtgattatttgaagagatgaacacagctagctgccgtcgatgatccatagct

gcggtagattatttttaaaaactcaatctaaaacctaatcccagtcggcactgctactgtgcttgtttatagtttttcaa

gcgaggccggatcttggacattgattttcttgcatcaccacttcaccagtatattacttgtatggcattagtcattagaa

accgtaactatacatgcatacgtaccggtgcagttgtgagcttgacgggcaaccgctccaaatgatgattagagggagag

gacgatgctgcactgctgcttccgcctctgtttgtcgcgtacttagcgtatgtacgtacgttcttcttcaagttaaaaac

acaatctcacagattcgggtcttggtctcttgcaataagcatattctaggaacacataccacgacacgacagatagcagt

tccaaaaaatatattaaaactattattctgacttcgtacttaacttgctaacgaactcaaactgattcgtctcttttcta

tatttcacctgatttcggacgcccggctacacccgctcggtggtggcagcctgcactgntccatcaagggcnaattctgc

nnanatcatcncncnntgncngnnnntcgagcatgcatctagagggnccaatt

bPb-9762

NACGGCCGCCAGTGTGCTGGAATTCGCCCTTGATGGATCCANTGCAGGAGAAGCAAGAAGAAGCCCAACACGACTTCCGG

TACAAGCGACGGTGAAAAAAGGCCACGCCCGCACCCACGCCCTACAAAAAAACATGCGTCGCTCGACCTGAGGCCCACCT

CATTCTCCATGACAATGGTGACCCACACTGGCACCACACGCTGACCAAGGGCTTCACTTGGTGAATGACCCAGACAAAAT

TTAGAGACACAATCCAAAATCAACAACCATAATTGGCTGTGAAGCCTTAAAATTTAGGGAGCTTAGTGTTATAGAAGATG

TAGCACCTCCGATCAATATAAACAAAGGTTTATTCTCAACTAGGTAACAAAGTGTTTCATCCTACCTCGTCATCAGTACG

TAAAACGGAAATTTTCACTCCCAGACTCAAAAACCATTCTTGCTGATCCAGTTGCATTACATGCGGCAAGTTAGCATATA

TCCGGATCAAGATTACCAGCTACTTTTGCTTGGGGTGCTGCA

**bPb-8308**

**bPb-9280**

bPb-9108

tgcaggtttaactcaagaagacaccaatgccaaagtatctatgtcaaccaagactaattctacagaactattagcaatct

ccagacctttaggcagctcacattattacgcgtgcatcaacaagttcagttctactcccaatcctgctacaagccgggta

gtaaacgggcgttttanctttttttcattttgcaacgagtgcgttttacgctctgtcccgtgcttgcagtacataaatga

gaTttcctcaattAtctgccttatTTttacTttgtaaaggccctttatCtaCttaaaccctttagcttaggaccagatta

attaggatatagtgaacaatgttatgcacgccattttaggttttgctgcccggtaggagactagctaaagctctagtaaa

atcagtagtgcttgatgaagattcagagaacaccagaatcatctttaagcatagtaactgca

**bpb-8935**

bPb-0617

tgcagtaattccgaagatttgttatttacgaagtaaaatgaacaagccttctcatattgggggattccttcctatttggg

gagtaagtttttcccatattccaatttttagttttttttatctcaattttcattttttctatatattttttgactgatTg

agtttttattttggaatgccaGatcatttatctgtattttgttgattttgatgttattgagaatcatcaatcaactgtgg

attattctCttcCtcggataactcatattaagaatgatgatttccagtatcttgcattggtcgatatgaattatgagtct

aagaaagcttttgcagttctatcagtaagcatcgtataatatatttcatgtgtttacttatactttcagctccggtattt

tttcattgttcattcttctttcatttgtagcttcgagacatatctcaaacaccctatgctaatgttgggcttgtgaacaa

tgctcctgttaatccttccgagtttctttcagttcctcatgtggaggtatgtcttttccttcattcagatttcattgaat

acatgattttcaactagtctcttgtttttttgcactatagtttaatatttatagataatgtcacttttaggtcattaatg

ccatgaatgttgacttagatggcaatgtgaagccagatcctcattccaagaaacaagtaaatttgaatattTtccccctc

gtacaatattgttcatcatttaaagcaaatttgttctgca

bPb-8112

tgcagatgtacaaaatgccgtcatcaaccatatgtgtgGgcagcacggtcacatcatagtggcaaggcaagagcagcagg

Ccagcacccagaaaacacagtagttccaccttccctgtctaatcgcggcaaacacggtcacagtcgtagaagtgatggat

aatttgatccttttaagaatgtattttgtctgatccctccctcgacattaccaaccggcaaggcagcttgtttttatccg

agagattattagtctttgacgcaagttggtgattatttgaagagatgaacacagctagctgccgtcgatgatccatagct

gcggtagattatttttaaaaactcaatctaaaacctaatcccagtcggcactgctactgtgcttgtttatagtttttcaa

gcgaggccggatcttggacattgattttcttgcatcaccacttcaccagtatattacttgtatggcattagtcattagaa

accgtaactatacatgcatacgtaccggtgcagttgtgagcttgacgggcaaccgctccaaatgatgattagagggagag

gacgatgctgcactgctgcttccgcctctgtttgtcgcgtacttagcgtatgtacgtacgttcttcttcaagttaaaaac

acaatctcacagattcgggtcttggtctcttgtaataagcatattctaggaacacataccacgacacgacagatagcagt

tccaaaaaatatattaaaactattattctgacttcgtacttaacttgctaacgaactcaaactgattcgtctcttttcta

tatttcacctgatttcggacgcccggctacaacccgctcggtggtggcagcctgca

bPb-0589

tgcaggctgggcccagccgccaccgagcgggttgtagccgggcgtcccaaatcaggtgaaatatagaaaagagacgaata

ctgattggtacttcagtttgagttcgttagcaagttataagtatgaagtcagaaaaaaaattatttcttttttgcaactg

ctatctgtggtgtcgtggtatatgcttattgcaagagaccaagacccgcgcgaaccagtgagattgtgtttttaacttga

agaagaacgtacatacgctaagtgcgcgacaaacagaggcggaagcagcagtgcagcatcgtcctctccctgtaAtcatc

gtttagagtggttgcccgtcaagctcacaactgcaccggtacgtatgcatgtatagttacggtttctaatgactaatgcc

atacaagtaatatactggtgaaatggtgatgcaagaaaatcaatgtccaagatccggcctcgcttgaaaaaatataaaca

agcacagtagcagtgccgactgggatcaggtttcagattgagttttaaaacataatgtatgggttgcactgagatattcg

aaggcccacaagctagctactgtgctcggccgtctggatgctcatgcatgaccgccgtgatggtgacggtgcagctatgg

atcatcgacggcagctagctgtgtttatctcttcaaataatcaccaacttgcgtcaaagactattttcaaataatctctc

ggataaaaacaaGctgccttgccggttggtaatgTggagggagggatcaGacaaaAatacAttcttaaaaggatcaaatt

atccatgacttctacgactgtgaccgcgcttgccgattagacagggaagggggaactactgtgttttctgctggcctgct

gctcttgccttgccactatgatgtgaccgtgctgccaacacatatggttgatgacggcattttgtacatctgca

**bpb-1487**

bPb-2260

tgcagatatctgaacttgaaaaagtgaagtccccaagagaggcacatcgtataaaattagtggaaaaacataggctcaag

gacttgaaattggagtggaccagaggtgttgagagatttgtctatgacaaaatgttgttgaaaaatctagtgccgccaag

cacattgaagaatttggagatatgtggttacaatggtgtgagctttcctgcgtggctagtgggacaactaccaaacctag

tctcattggttctcagagatatggaaaatgtggaagagtggaacatgtcatactccattgatgaggaccatgtgattcaa

acattggaaatacatggttgttccatgttaaggatgaaacgacctcttcctaaagctaaatcatgggtgatagcacacag

tgataatgtattatcatcattggatgagtgcactgtgtcacacaccagtgctttcccctgcacttctccgataactaccg

tgttgttgtcggttaaatactgcaaggggccttatcggtggaggttacttcaacacttcactggactctcttctttgtca

attcacaactgcggtgatctaactggatcaccagcgatcaccCatcatctatcgtctctcgaggcactatgcctagaaga

cgaacatatggaagaactgccaaaatggatgggtgagctcagatcactacacagtctggaaataaggtggagcaatgttg

taacagagttaaatgagacaatgaggaaactcacaatgctgca

**-------------------------------------------------------------------------**

**Ch. 2**

bPb-8511

tgcagcgaagaagatgtagcatagagtcatgcagttcgctggaagtgtgttgaagccatccacacccaatatcgccatgt

ttcggcacacaatggcatatggtgctgggtattcggtgccgcttgtaccgatgttatgtatatcgaatctgattagttat

tatggacgaacaatgcatattagagccaatgtaaattcagtgaaagcaagtagattataagtacaatgaaatttgtcaca

tgctcatgcaatcaaatggtagagcaacaactagtccactaatcacaagtaatagacataacttatttaaaaaggtcagt

tcttccttcctaacaccaaaaccctttgaccgagcatatgcattatagtaaatgtaaccttcttcctcgctgccaaaaat

ttgactgatcaccatgtcatattctgcaatgttctcgagggtggcactcccatattccatgtcaaaccctgcaattaaaa

agaatctaatctaagttatttgaattccacccatgaaactaaaacacgtatcagtaacaatgcttgtgtacataatcaaa

cagaaacattagatatactgca

bPb-7930

tgcagcatgttggcatgggccggcatactagcgtgctgcccacttttattcaatttgttattttcttgtgttcatccagt

agaaaaaaatcgcagattttttatttgttacgaaaataaaaatatgaaataaaaaaaggtttacgaatttaaataaaaag

gttcatccatttgaaaaaaatacttaaaaggttcatcaattttaaaaaaagttcatcaatttaaaagaaaaaaatcattg

aaactggaaaaaagtcaactgacgcttagatgggccagctcatttacgaacaccacacgagttaacaaaaatgcacgata

actgaataggaaatgccacaggacggcctgca

bPb-6894

tgcagtccatgggcgataacccctccaccggcaggtgctcgccccggtcgatcatcctccaaacctcgttcatcgccacc

gtggtctgcgtcgtcgtgggtacgtagtttgcattaattgtttcagccagtacatatatagaacattCggtttaagtgat

atatatcaatCgaattacacagatagaatatataAcattcagttcaagtgaattataatcacatgcatataccgtgcgta

ccggattgttggcgaacatgccggcgtcaattaggttaaactcctccaattcgaaatttagtggacacaagtttataatg

tagaacttgtgggcagggaagaatgttggcgcggcagttgcggcgatgcaaatgtccgagaggttggggttcatcacccg

atcctgttgcgcctgtcaatggttcagcagctcgatcagtacatccGatgacgatcagttggttcagaagcagcgcagta

attaataagtagctagcttagctagggtaccttagaggtggagaagaCcacaagctgattgcgcttgatgtcgaaCgtgg

gcacgatgacgttggtgacggtttccttaagcttgcggtcgccgagtttccctctgaccaccttgcggagcccctcgccg

tcgtacttggggcggagcagcgcgtacctgacgaaatccgggttcagcagcgtcatgcgcaggccctccacgaactcctt

gatgcactcatcgaaaggcgtcgagtccagcgccagaccctgcgcacggcacatcgcctgca

bPb-7793

TGCAGTAAGAAAAGTTGCATCTTTCAGGCAACACAAACACACTGTCCACTTCTCTGTCAGTTCATTGCAATTGCGTTCTT

CAATCTTCAGTGATAACATCCCCTCTGAACAGAGATCATCATAATCTGCGTGTTACTCCCCTACGGGTTTTGGTTAAATC

ACTCTTCTAGTTTGACCTTTGGATAAATCTGCATATTCCTACTAGTTTGGCCTTTGGACAAGTCTGCATATTCCTATTAG

TCTGCGGGTAGTACTAAATTGTTTCTGCTTAAATCACTCTTCTATGGTGCTGGTTCTGTTGGTGCAAATCAGGAAGTGCT

TACTGGAGTATATCTAATGTTTCTCTTTGATTATGTATACAAGCATTGTTACTGATACGTGTTTTAGTTTCATGCGTGGA

ATTCAAATAACTTAGATTTGATTCTTTATAATTGCAGGGTTTGACATGGAATATGGGAGTGCCACCCTCGAGAACATTGT

AGAATATGACATGGTGATCAGTCAAATTTTTGGCAGCGAGGAAGAAGGTTACAATTACTATAATGCATATGCTCGGTCAA

AGGGTTTTGGTGTTAGGAAGGAAGAACCGACCAGGAAGTCGGGCACGAACATATCTTTTCGTCGCCTTTATGTCTATTGT

AAAGAAGGATATCGGGCGAGGAAGCACTTCAAAAAAACTGAACGAGTGTGAACTCCTAGGCCGCTGTCACGTTGTGNATG

TGGCGCCCGGATGGAGATCGAGCTTCGTATGGATAATGGTGAATGGTTTGTAAAANATTTTGTGGACNAACATAACCATC

CACTCGCTAAGCCTGACCAGACAGCTTTCATAANATCACATCGCGGACTGAGTGATGTGCAAAAGGNNNNNNNNNNNNNN

NNNNNNNNNNNNNNNNNNNNNNNNNNNNNNNNNNNNCAAGTACAAGAAGAAAAAAATGTTTAGCACAAGTGATGCATCTC

CTGGAAGTTTTTTTTGGTTGCATTGTGTGACAGAACATTTAGTATGCTTAACAAGATGACTTTTCAGAAAATATATATGC

AGTACANNATGAAACACAAGGAGTACTAGTTAATAAATNGCAACCTTTTTAAATAAGTTATGTNTATTACTTGTGATTAC

TGGACTAGTTGTTGCTCTACCATTTGATTGCATGAGCATGTGACAAATTTCATTGTACTTATAATCTACTTGCTTTTACT

GAATTTACATTGGCTCTAATATGCATTGTTCGTCCATAATGACTAATCAGATTCGATATATATAACATCGGTACAAGCGG

CACCGAATACCCAGCACCGTATGCCATTGTGTACCAAAACATGGCGATATTGGGTGTGGATGGCTTCAACACGCTTCCAG

CGAACTGCATGACTCTGTGCTACATCTTCTTCGCTGCA

**bpb-3119**

bPb-6199

TGCAGCGGTTCAGCAGCGCAAACTCTCGTTGTTCCTCTCGGTCTCTCCTGATCCACCCATCACAACAGCAACAGTATATG

CTTATATATATGGGATGTGTCGTGGTGTGTGCATGAGCAACTGAAATTTTCTAATAAGTTACATGGATAAGTTGAGTATG

TTGTAAAATGCATTAACCACTGAAAGATTGGATAAACATGTGCACCACATAAACNAGCATGCTATAAATTCCCATCCNAT

TACCTTTGAATGCTGAAGGAAGGAAAGCTTCAAAAGCTGAGTAATGTACATCTCGCCGAGCTCCCAAGCGGTGCATGTTG

GTGTTGCCCCAATAAAAGAAAGAGAAATCCACTGGTGAATTAGATCACTTTTAGATATCCTGTATCCTTTTGGAAAGATT

GCCCAATAGGCAAAGCAGAGCTTCAAATACCGAGGCATAANAGTATAGCTTAACTTCNAGGATGCAAGCACCCGTGTTGC

AGCTTTATCTTTGGGATCTGATGAAGAAGAAGATTCTTTCCNGATATCACTGTCTCTCANTTTCTTCCACTGATTATAGG

TCATAGTTTCCAAAATGGGTCCNNTTGATTGAGCAGCTAANCCACACCTCCNCACTTCGCTGCNNNNNNNNNNNNNNNNN

NNNNNNNNNNNNNNNNNNNNNNNNNNNNNNNNNCATTGCGTGTGGTTGCTATAACANCCACCTTGCCACCCCTTCCANCC

TTAAGCATGNCAGTCAGACTACGCAGTTGGGAATCATCCTCTGCCCACAAATCATCTAGAATAATAAGANCCTTCTTATT

GGCGAGTAGCTGTGCAAGTGATCTGTGTATCATCTGCGTTTCAGTGTAGGTACTCTCATTCTGTGACAGTTGACTTATTA

AAGAATTGCCAATTTTGTTCAAATCAAATGTCTCGGACACATAAATCCACACACGGACGTATTCTTCAAATTTTGTATTA

TTATTGTAAATCATTCTCGCCAAGGTCGTCTTGCCAAGGCCTCCAATGCCGTGCATAGCAAGGACAACGAGGTCTTGGAC

TGTGTTCTCAGATAAAGAAACCCATATTTTATTTTCCTCTTCGGTCCTCCCAATGATTGAATCCTCAATGGTTGAATATG

TTTCTCGCATTTGTATGAATTTCAGCTCATCAGCATTAGTGCCCGGCACTAACATGAATTCCCGATACTGATGTGTGATC

CCCTGCA

bPb-5173

tgcagtggttggtttggagcaagcactttgagtggcgcatatgggcttcgctggtcccgatagtcggggggatcctcctg

acctcaatgacagagcttagtttcaacatttttggtttctgtgctgccatgataggctgcctggccacgtctaccaagac

catcttggcagagtccctgctccacggatacaaatttgacaggtacatgcttatctcgttgtgtaatatctgatgcttta

gcgactggtgttgtatcgcattctcaattttaaagcattttatcttctttcttactaaaatgatactagatcaaaagatt

tagttcttactgtagtgctaccgtcaggttcagtcctgtaagttcttgcctcggtcagatgatttattaagtattctctg

tttctgcactggatccattcaagggcgaattctgcagatatccatcacactggcggcAgctcgagcatgcatctagaggg

cccaattcgccctatagtgagtcgtattacaattcactggccgtcgttttacaacgtcgtga

**bpb-9683**

**bpb-9320**

**bpb-5363**

**bpb-2300**

bPb-7445

TGCAGGTAGCACCATGTCCTCTGCATCAATCCATTTCGATCTGACTGTAATTCTGTTCCATTCCTTGCTTTGCATGTGTG

TGTGTGCGTGCGTGATCCTGCTGCATCTGTTTGCAGGTGGCCGTCAAGTGCAGGCCGCTGACCGACAACGAGAGCTGGCG

CTCGCGGCACATCATACAAGTCATTGACGATAAGGTAAACATTCAGAGCCTTGGCTTGATCCTGCTTGATTTGGCTGGTG

GTTTGTGTCATGGGTGGTTACAATTTTGCCATGGATGATAATGGACTAATGGTATGGTAGCAAGGTAGTAATAATGTTGT

GCTTGTATGTTTCTCCAGAATGTGGCCGTGTTGGACCCTGATGTCTCCAAGGACCGCCTTTTGTAATGATAACAGCTGTA

ATCAGCAATTGATTCAGCATCAGTTGGAAAAATAAGCAGGGATATTGTCATTAAAAAGATTACAGTTTGTGAAGCTTAGG

CGCTTTGCAACATCAGATTCGAGTTGATGTAGATATGCCATGCTTACTATTTGACATGAATACTTTGTGTTGTGCTTCTT

AGCTCTAATAGTACTTTGTGTTTACTTATGTAAATTAAAGGCCAACTTGTGAATTGGAAGGCTAGCTTGGATGGTGCTGA

TGAGTTCCTTGGATCATGATCGATGGCCAACTCCACCAGCCTCTGCA

bPb-7418

tgcagagcagagcgggcagtaggggtagtagcgataggacttcctctgtcgcatcagtagagtagaggccttgctctggg

gagcatcgcagcgtcgtctctgttgtaggaggagagaggggtacccaaaagggaaaccccctcctctttgtcaagtctcg

tagagtagcgcagtggtttgcatgtgatgtgctaatcaggcggtcgtgggttcgagaccccacggcgccaaccccccttt

ttcgtgttttgtttcgcagagtagcataagcaacaacagagacaatttatcctgttagccatccatgttctgtcgagcat

atggacagtagcagagtggtggcttgttcgttggtgaaacacggggtcttgggttcgaatcccttccccatccttttatt

tcttttgcttttcttttactgttttgaatatgtttgcttgtatagttactgca

bPb-9870

TGCAGTGACTAGTGCGGATCACTCGCTGCTGCTGCCTGCTATGGGAGGTGCTTCTTAATTTTTATATAGCGGTGCCATTG

CGCGCGGCGTTGTGGTTGTGGCGAGCACCTCGTTGTGGGCGGACCACTGACAAGGTACAACGCGTAGGCGTAGCATTATA

TTTGTATGCTGTATATAATAACCTTTATATGCACAAGCAATGCATGCTCCGCGTGTTTGAAATCTGAGACCACTTTTTCA

ACAACCAACCATGGATCGCTAGCTTATTCCAAAGTTCCAGCTCAAAGGTCCTTGATCGATGGCCATTTTTTAATCCTCCG

TTCATACAGGCTTCTTTAATCACCACTAGTTCAACCGACAGGCTCCTCAATTTGTGCCATTTGCTTTTCCTCTTTGCACA

AACAATAGTAACGCTTTTTACTTGATTCTTTGCAGGTCTTTTTGTTGGGGGGGGGGGGNCCNNNNNNNNNNNNNNNNNNN

NNNNNNNNNNNNNNNNNNNNNNNNNNNNNNNTTGGGGNGGGGGGGGCCCTCCTTATCAGTTAACTAATTATTTTTGCCCC

ATTAATTTTCACCCGTCAAAAAATTGCCAGAGTCAATTACACCCGCAGGTCTGTGAACTTTACAAGAAAAATCAGTTTGA

TGCTAGAACTTACGGCATACATTGAACCGGTAATAAAACTTGACTCGGGCATGCATCTATGATTCTAATCATGTTTTGTA

TACATGTAACATACCGAAAAGGCACACCAACACGGCGTGGGACCCAATGTAAGTGACGGATGGCGGTGTCGTGCGTGTGG

CATGATTTTTTTGCAAAAAGAACACCTGATATTTTACCTGTTCACACAAAAAAGCATATCAATTGTTTTTCTTAAGTAAC

CTTGAAACATAACTTTGACAATTGGGCCCCACCCACCAGAACACAATACAACAAAGGAAAACGAGGACGCAAAATTAATC

ACTGCCAAGTTCAAACACACGACCTTAAACGTCAGTGCGGAAAGGCTAGCCACCAGCCGACCACACCATATATCACTTTG

TATAAATGGGAGCTCTATCCGTAATGACCTACAGACAAGAAGGCACATAGGCCTTAAATGGCCTGCA

bPb-0205

tgcagaggagaagcttgcaaggatgctggcaaagaacaagaagggagacgacgagtggggtttaggtaaaagtagaactc

gtctggtctcagacggtgctaatattgctctggagctgctcaaggctgatggtttgaagatgccgaaacttttggagctc

gtcttcaagatttgggtggataagctgttgtatgcagccacccgatgcaacagagaatcccatgcgaggcaactcggccg

tggcggtgacctaattaccatcgtatggattatggtagagcatgttggcccgtttgagattggcaaagaaggtccAgatg

aaaaagaggaatccagtgatacagaagatccgggcgatggaaCtggtgattggatcaatgacaatgaaCacccctcatca

gcggaagacttaGaatcacaagggggggctgggggtgggggctggccacatccaccatcatggttgctaccgccggcaat

gccccaacctatgcaaccacaatggtgccctccccagccgatgcatccgccatgtgatggtggacatatgttcccgtggc

cgtggcttcaagcgccatgggatcatccgacgacatacttgccaacactcggagacggcggacatattatgccatcgcga

ccagttgcgggggctccacccccttcccctcccggtgcccctcgtcctgca

bPb-6897

tgcaggttacaaagtcttggaggtcggtggctcggaggaggtgtacgacaagatcatatttggtgctcatgcacctgatg

ttctgagaatactaggggatgaagcaacacacgaggagttgagaattctgggtgctttccaatatgtccataggtatgaa

atttgtgtctgctataaaatttaatttactcaattgataagtgccgtacgtgtggcaccaaacactttagtcctcaagac

aatgttttaaatagcgggctatgacaaactagcggcaggccttaaaatcagctgtagcaggctataacgacatatttgca

tacgataccatttgacggcaccctgttgaaatggttatagcggggcttcagccggttatttaaaactacggaaaggtgcc

atagcacccgggtctcccgacacccgtttatctggtgcgtccagtttttttgaagattcgagctgaggactagacaacag

caggtcatttgcgtctgcatgttaattaaaaAgttattttacatagcagaacatggtcggtcgtaggctgcacgagacag

agtaggtgtgacagcaccgtgctgca

bPb-0715

tgctcgagcggccgccagtgtgatggatatctgcagaattcgcccttgatggagccagtgcaggttacaaagtcttggag

gtcggtggctcggaggaggtgtacgacaagatcatatttggtgctcatgcacctgatgttctgagaatactaggggatga

agcaacacacgaggagttgagaattctgggtgctttccaatatgtccataggtatgaaatttgtgtctgctataaaattt

aatttactcaattgataagtgccgtacgtgtggcaccaaacactttaatcctcaagacaatgttttaaatagcgggctat

ggcaaactagcggcaggccttaaaatcagctatagcaggctataacggcatatttgcatacgatatcatttgacgacacc

ctgttgaaatggctataacggggcttcagccggttatttaaaactacggaaagatgccatagcacccgggtctcccgaca

cccgtttatctggtgcgtccagtttttttgaagattcgagctgaggactagacaacaacaggtcatttgcgtctgcatgt

taattaaaaagttattttacatagcagaacatggtcggtcgtaggctgcacgagacagagtaggtgtgacagcaccgtgc

tgca

bPb-2243

TGCAGGAGATCGAATAAAAATAAGAAAAAATGGCAACGAAGCGGCAGCCACCAATTGCGCACACGCGCACAAAGAGAGCA

AACAGGAGAGAGAACGTTGGATGGACGGTACCTCTCACACTTATACCACCCTCTACAGTTGGAAATGCAGACTATGGACT

AAGAATAGAACAAAAGTGAATCAAAAAATATTAGCTTAATCTCGATGAATATAGATCTTGATGCCTTTTGAAGGCCAACT

ACTAAGGTAGTAGTGCACACCATGGTTCCCATGGCTTTTTCAAAAGGAACGCAACACTATAATAGCGAAAACCTGATGTA

TATAAGCTACAGAAACAATATAGATAGAAGAACCAAGCATGTCCCTTTCATGAGAATACAGTAGGGTGGCTGTTTCTGCA

CATGAAAATAAAATCCACCAAAAAGGGAGAAAGAAGAGTAAAACCATAGCTGGTAACTTGGACACACATCANATATAGTA

CTGGTATAATTTAAAGCATAGTAAAGCTTGATAAATTAAAAATAGAACTTTGCAAACAACATGTAGATTGAAGTATCATT

TAAAATGAGTGAACTATTCATAGATACGTACTTTGTAGGTACAAATATCTTTCTCGAAAATACATTCCATCAAAG

bPb-9131

tgcagtcctgaaggtagaggatacacgatctttactatatttttatgggtgtctacattgttgtacacactttgatggaa

tgtattttcgagaaagatatttgtacctacaaagtacgtatctatgaatagttcactcattttaaatgatacttcaatct

acatgttgtttgcaaagttctatttttaatttatcaagctttactatgctttaaattataccagtactatatttgatgtg

tgtccaagttaccagctatggttttactcttctttctccttttttggtggattttattttcatgtgcagaaacagccacc

ctactgtattctcatgaaagggacatgcttggttcttctatctatattgtttctgtagcttatatacatcaggttttcgc

tattatagtgttgcgttccttttgaaaaagccatgggaaccatggtgtgcactactaccttagtagttggccttcaaaag

gcatcaagatctatattcatcgagattaagctaatattttttgattcacttttgttctattcttagtccatagactgcat

ttccaactgtagagggtggtataagtgtgagaggtaccgtccatccaacgttctctctcctgtttgctctctttgtgcgc

gtgtgcgcaattggtggctgccgcttcgttgccaTttttttcttatttttattcgatctcctgca

bPb-7011

tgcagaaaaataaataaataaacagcaatggatgacagtatgcttaggcgtgaacaccatgatctgtgtggcacctagct

tcctcaccttccctagcaaattttctgatcaattggcacgcgatggacgaacccccgggtgtgtccaacaaggacttgaa

cgtcgcgttcagcgccgaactgtcggtccgacaggctagcgcctttgacgaagtatcttgttcctgcatcacactggttc

tgtttctagcatacatctctatcatagaaaccattgttgaactcttcatggtcactccaccattattcatctctttcagt

atcttatccgcatcggtgatcatcccacttttgcagcaaacttccattacagtcatgcagagggcctcatcaacctggat

atcttcttttctcatgttcagaatgaacgcacttgccttctctagctggcctagcttgaaatacaatctcagtaggtcat

tgcagcagaacacatcagggaggccatatttgcacagagctctgaaggcctcttcagctgca

**bpb-9832**

bPb-1967

tgcagatgccggaggaatataatcttttcaggaaaaatctctgtcaaatttacattatttctgagtgtacgatcgaacac

cataactcaactaggacctgatgagcaactaacatcagtaacagcctgttcctatgttgcagaaggcggataattcaaga

acaaagaaagaaaataacacaatgcaggaatcagaaggcaggtaactctacatacaagaaagcagatggcacaacacatc

tgatagagaaaaaatgcatctgacaaagttaggaaagtagttctacaagagtagcaggaggattgagaaaactctctatg

cattattaagctggtaagcaaatggctaattatgcagaaagaatattatttacagctagcgtagctgcttgaaaacctct

agggtatgcagatagcatatttaatgtcgcaagatcggcactatgactattcaagaaaccttggtaaatttagcaagaac

attcacctactgttgaaatcagtgcccgaaagcatcagaagggataggagtaaaagcagacaaactgttatgatagagaa

taagAagggaattgaccacactttgattcagccataggcaGagaataTAaGtacaggttggagccActcctagccTagtg

gaGatcAtgggttgCccactactctcggacgcgtgcangggtacaaactgcactggatccntcaaagggcgaaattctgc

agnatntccatcacactggcggccgctcgagcatgc

**bpb-8033**

bPb-23426

tgcaggagcatagaaataatgggatggaatctcgcacaggaccaaaggttgttgtttttatttagctgaagtatttgttc

tttcacaactattttaatgtgtatagAataatttagcttgtccccttgttttgcaggaaccacCGGAgaatgctgttata

acaacatgCtgccatattttctgctatgaatgtgcacaagagagcttaagtgaagaggaagtctgccctgtttgcaaaca

gaaattatgttCtgaattgcttttttcacgcccagtactaaggctcTgtatctctgatgagttggagtcatatgcaacag

cagcaggagatagttctgca

bPb-8050

tgcaggagcatagaaataatgggatggaatctcgcacaggaccaaaggttgttgtttttatttagctgaagtatttgttc

tttcacaactattttaatgtgtatagaataatttagcttgtccccttgttttgcaggaaccaccggagaatgctgttata

acaacatgctgccatattttctgctatgaatgtgcacaagagagcttaagtgaagaggaagtctgccctgtttgcaaaca

gaaattatgttctgaattgcttttttcacgcccagtactaaggctctgtatctctgatgagttggagtcatatgcaacag

cagcaggagatagttctgca

**bpb-8851**

bPb-8721

catgcgctcctttgttgctcttgttgcaataaaacaaaatcaatctactactgtattacgagtaacttattatacggcgg

cgcacaagtaaccatgcggtactccttgccaaacatttgagcagtagaagtaccttcatcgatccggatacgcagtctac

gaaagggcgaattctgcagatatccatcnnntggcggccgctcgagcatgcatctagagggcccaattcgccctatagtg

agtcgtattacaattcactggccgtcgttttacaacgtcgtgatnggaaaannnnggcgagtgaattgtaatacgactta

ctatagggcgaattgggccctcnagatgcatgctcgagcggccgccagtgtgatggatatctgcagaattcgccctttcg

tagactgcgtatccggatcgatgaaggtacTtcTactgctcaaatgtTtggcaaggagtaccgcatggttacttgtgcgc

cgccgtatAaTAAgttacTcGTaatacagtagtagattgattttgttttattgcaacaagagcaacaaaggagcgcatg

bPb-7874

tgcaggaccactgccattggcgttctgctctccggcatccgccatggctctgaaattcctttncaaatctctgncttttc

tctaaagattgtagcctatatgtgggtnggcaagactgagatgattgagcggataacaatnncacacaggaaacagntat

ncnanncaaggccaagctnncnncgagctcggatcccttagtaactgccgccagtgtgctggaattcgcccttgagtagt

gccagaacggtccatgtagttttgtggtgtaattgcctagttggtcactaaatctgcactggatccggatacgcagtcta

cgaagggcgaattctgcagatatccatcacactggcggccgctcgagcatgcatctagagggcccaattcgccctatagt

gagtcgtattacaattcactggccgtcgttttacaacgtcgtgactgggaaaacaatcat

**bpb-46599**

**bpb-6881**

**bpb-3056**

**bpb-6438**

**bpb-7991**

bPb-3563

tgcaggcgcactttaggggaacattgttctgtggaatgcaggtgtaccttcaagttgccttatgcgcaatatgtgcgtca

accatgttatattatgttcccttagagttgggtcacgcgctcatttgttttagggcataaatacgtatgtgtgactaccc

taaacagaacaaaaatggaaaggggtcaagggacctttgcttggtttgattaattcacgtgttaagattgaagtgtgatc

ttgcattgatgttctacaaagtgatttgctaggaaagagtgaattacagaaagatacaagtttaatatatgatatatcgc

taaactacagattgaagaacagatatttacaactattgattcaagtgaacaacatgtagttttgtggtgtaattgcctag

ttggtcactaaatctgca

bPb-6194

tgcagatttagtgaccaactaggcaattacaccacaaaactacatgttgttcacttgaatcaatagttgtaaatatctgt

tcttcaatctgtagtttagcgatatatcatatattaaacttgtatctttctgtaattcactctttcctagcaaatcactt

tgtagaacatcaatgcaagatcacacttcaatcttaacacgtgaattaatcaaaccaagcaaaggtcccttgaccccttt

ccatttttgttctgtttagggtagtcacacatacgtatttatgccctaaaacaaatgagcgcgtgacccaactctaaggg

aacataatataacatggttgacacacatattgcgcataaggcaacttgaaggtacacctgcattccacagaacaatgttc

ccctaaagtgcgcctgca

bPb-4184

tgcagtgcaaaaccacgggtgtacaatgatcctttcaacaatccgttagtcaagatcgatgaaggtacttctactgctca

aatgtttggcaaggagtaccgcatggctcctgtccggcttacaaatgaacaacagacaatgcatcagaaaaggcgatcac

atgcatatcagtggaaaaggcccaccgtgtttcttaaagaaggtgatcccctgccaccagatgtcgatccagagacagtt

agatggattccagtaaaccacccatttgctgctggttcagttgaagtagatgaggaggctgccaaaaggaatgtttatca

aaaggatggtgtcccgtcccgtgttaaagccgaacatgaagctttgagggcaaggctagaggcttctaatgatgtgagtt

tttgcaataaacatgatactctgaaccggccaaatgctacaatgcagaaagacttgtgctgctacttatttttaaatgta

gttctatatatgctgtttgcagacctctctgca

bPb-8949

tgcaggcggctaatgggaaccatcccgaaaatcagactatagAccaAtccaaggtacgtaacagataagtacttcccgat

tcccgatttctacaattgcccttgcttttttcaggtcccaaacaggccatttgcgtattctgaacattgattgtttccat

caagcgtgcagctcgtctccctgtctcggaattgtacactcttatagccttgtcgaattccctccatctctggacagttg

gcgcatagatctatgcacatagatctggtaagtaaatgcaagatggattttgctatttttacttatttttgttgtttcat

atgttgtaggttgccattgtacgttctgtatatatatatcttttccactttggatataggtaattccagcccaaacaaat

tcatgacaccttttctatgtgaatttatagttacctatttattttatttgtctgacaaaaattctaccaattctcttaat

ttaaatcactgctcttgagtgcataatttttgttaccagtataaatgatcagcactccaaatagtgtgtttctagcagaa

taatacgcagaagtgattttttccccgcctgtgcaacattatccttgttagagcttatcttgttttctttttggcatagc

acatgactagttcttgtttccatatattagatggcgcctcatgcacagataactttattgataacctctaactagTccac

tactatgaTtttagtttttctttttcttctttctatccctagccatatgatgtattcaccatttgttttggcatgatctg

ca

**bpb-3131**

bPb-5755

tgcagcagccatggaagtccaTgCtgttacacggaacagcaacgtcgggatttattgccatatagccgtgcagtttgaat

agtagtacagtatgaagctgtgcattatttgtgggcaagcggtccggCttttctgataattggtcaGctttctcttgacg

gaggtttggtttcctgtaatcaatgcccttaattcggcaagctgcaaccaatccgatctcatgagtttagagtatatact

tctgccaacaagatcggagagagttccttttgcaggctgtcagttaggctgtcgagcactggcagcTgttccattgtgct

ggaaataagttcgggttaTtttctgca

bPb-4228

tgcagtacagcagttgctagtaacgaatcaaacagtgtttcaagaacctaaaacactgcctcctcatagagtattggatc

actctattcacttcgtacttgatggtatgcctgtcaattgcagaccatatagatactctcatgtgtagaaggatgaaata

gagagacaggtggcagaaatgttgaaatctgGtctgattacacccagtgtgagtccttttgcttctcctgttttactggt

caataaaaaagatggaacatagaggttttgtgtggactacagaagaataaatgctttaaccattaaaaaaatcctatgcc

agagattgatgacttgcttgatgaattacgggacactaagtggttttccgaactggtccttagagctggatatcatcaaa

tcaaaatgagggaagaagatgaatttaagactgcttttaaaacacaccatggacaccttcagttcagggttgttccattt

ggtctgaccacaacccgaaacacttttcaatgtgtgatgaattttgtgtttgcagggccaaacagaaaatatgtgttggt

tttcatggatgacacattggttttcaatcaaacatttgaggaacatttagaacacctgca

**bpb-4093**

**-------------------------------------------------------------------------**

**Ch3**

bPb-8547

NNGTGTGCTGGAATTCGCCCTTGATGGATCCNNTGCAGGCGTGAGGATTGCCTCATACACGACAAGGAAGAAGATCGTGT

CTGGTATGGCGAGGAGGTGAAGCGCATTGATGGTGAGGAGCTGAAGCTCATACTCCATGCAGGTTCGGAGAGCAGCACAT

GACGCCAACCACTGCGATTGCTTTCTGGCCCATTGGAGCTACCAATGTTCATGTCAGTTGCTAACGACACCATCAGCGGT

CAGTGGGTTCACCCTCCATAGTTGGTTCGACTCTTTCACACCACTCATATTGGAATGCGCCAAGAATATTTGTTTCACTT

CAACATCTCATAAAGTACTTTTGGGAGTATTAAACTACTGTCCGAATGTATTGCTACTAACTAGTCCCGGACGGACTGCA

TAGATATAGCTGATTAATTTACTACACTTGCAGCTGCA

bPb-0634

tgcagtggagtgggacattgctgctcgccacgagccaagtatagcggtgctgcaatggagcgggcgctggttccgacggt

gcgaccagggcgcttcaatggcatgagcgttgccgtcgtcggcacgacaagtcaagagcgacagtgctgctgccggtagc

gttgtaatggaacgagcgatggcgctgttgccattggatcattggtgctgcaatggagtatttgcgcatggtggtgcctg

cgcgacgatgaagcttcaatgcatggctggtgttgctatggagcatcgccggtgttgctatggagcattccggagaatca

ccggtgttgctatgtagcatcctcggccgtcggggcatcatcaatgctggaatgaggcatcccggagagttgccggtgtt

gcgctagagtgttgtcgggccgccggagcatcactggtgcttcatgggatgatcgcaagcctgca

bPb-6797

TGCAGTAGCTGCTAATTTTTTTTGATGGTGGAGGGGCACGTCCATCTTTGATGGTTGCCTGTGATTTTAGGTTCTAAGTG

ATATCTGATTTGACATAGTACTGATCCTAGTTCCCCCTATGTGTTTTTATATAGGCGTTCTGCAAAGAGTATAGACAGGT

TGCTCCTCCTGTTCAGTCGAGGTGTCTTTTTGGATCCAGAGATGGCTGGCAAGTAGACAGACAAGGAGCGCGAATGACCT

TTTTAAATTTTGTCGTCATTACTTGTATCAGTTCGTTTTTTTTTTTTTTGCT

bPb-6884

TGCAGAATGGTGCCGTAACTTTTTGATGCAGAGGCCGGTGGCTGTGTTCCTCCTTTAAAAAAAAGGGTGTCATAACTTTT

TATTTGCTCCCTACATTCCCTATTAGTTGTTTCGCTATGACCAAGGAGGAGCTTCGCGCGAGGGAGGTGTACGGGTAGTT

CAATAACTTCCCAGTGATGGGATAAGCACCCGTGGATGTGTAATTAACGAGTGGGGCAGAAATCAGCAACTCGGGTGCAA

AATGATAAATGGGACTTAGAATTCGAAAATTTGCTAAATACAATACCTTAAAGAATGGAATGGAAGCAGTAACGTTAAGT

GAAACTTTAGCGACAAACGCAAAGGTGAATTCAGGAGAAGTCGCGAGCAAATGAAATCAAATGCATGAATGGAGAGGTGG

CTAACCTTTGATTTTCNNGCCTTTTCTTGCATACGCTTTGACAAAAGTTTTATGATAACTTATCCGACTGAAAAAGTATT

TAATTTTCTTATGAATTCTCGACAACTCATCTTCCGGACCACTCACTGTGGGTCTTTTCCTTCTTCTTTTCTTAGCAGTT

GGTTTCACTAGAGTGTCCACTGAGTTGCTAGGTGGCTTCTTCGCGTCTCTCGAACGAAGAATCCTGACCTGAGTCTTCTG

GATAACGTTNTTGGTCTGAGTAACGTCCTTAACAGAGGCAGAATTATTTCTATTGTNNNTCGGCGGCGAGAAAATATATT

ATTGTAGGAAAAC

bPb-2018

TGCAGAGTTACCGAGATCTATTACAGGCACCGTGTCATCAACATCATCCCACTTGTGGAACCATCAACTCAGTGCCCATT

TCAGAAGCTCATCTCAGCAAACTTAGCAACTGATGTGTACAAACAACCTCAATCATCACAACAAGTTATGACCATGGTAC

GTTGTTCAAGAGATTTCACACCAGTAGATCAATACCGTATAGTAGGCCCAGCCTCTTGTGTCAGTAACAACGCTAGCCAA

TTCTGGTATATGGCGGTATCTTATGCATACATATCTGATCTTCCACGAGACTGCATGGCTGTTTCTAAAGGCATCCCCGT

ACCCTTCACTTATGATAATCATGGCCCCAACTATGATTACACCTTNNAGGCAAATACAGTCCTCAACTCTGGAGAGACAA

bPb-44917

tgcagctgtactggtactcggaagaacagatacggcaggaggtcaaattcttagcttgtagttttatgCtggccttccaa

gtcatttgaatgttatttcattttttgcgaaagcatctgaatgttatttatatggctaaaattaacgcagattaaatggg

aatgccgtagcaatttgttctcggtctacaagtcgttttgacgtttacatgtctggaattattgctgtagaaattcggat

gacagatactgaacttttcttgccctgagagcgacgttgctgctgctgtccagttatttatagtggttcatatgaggcag

aggactcgtcgtccttgcagtgcttgtcaatcacactcttaattgcaggaacgcatacggcaggaggttgcattcttggc

ttgtacagtagttttctgctgaacttcaaagtcatttggatgatttttatacggttgaaactgccgacgattaaatcgga

atgcgtgtgacagcaaaaaatcagcttaatctttctcaccctgagctgacggtgaggttgctgctcctgtccactagttt

ggtccatatgcaacaatggactgca

bPb-7235

tgcagctattcatcaatgcaatcaaaacccatttacatatgacctagctgtccaccataaaggcttcaggaatcctccgg

tgggaaagttgcctaagagcatggttaatagtatagccaactgctggctataagcagtcttatagttcatcttatagcta

gtttgtataatagttagctataaaaaaaatacaacatttgtcatatatggcccacctttcattctcacaaaacacctagg

aacacgtgctagagctggctcttcacgaagagtccgcttcccttctctctcctcttctctcacattcaactcagcaaaaa

tatagtattttaattcttataacctgctgactgtaccttactgtacttgctctaaaaaaaacgtctcgtttcacgaggag

tccatcagattgggtcgtgcctctcttcttgttactgctcctgactcgcacgcgtcatcctcacccttgaagccccatgt

ggcgtctctgca

bPb-42312

tgcagtccgttgtgttataatagggctaagctggctctgttatggaggagatgctttctctaccacagaaaccttttata

ctcataggcctcttggaggctctagcagcagcgtaagggatggctgctgggggtgattttgtttcttctaaaaagcaaat

acaacatgttcctcatatatcttcacttgtacaatgtacttgatatgtgacataaggaactcaaaatcaaactattatat

gcttacttttgctggggcagctattctttttgggacttcaataccaatattatcacatgttggtgggtcatcagtataat

catatgacagttttcactttgtcacctcacctgcaatcacatttatataattatcttactaccttgctgca

bPb-1114

TGCAGCAAGGTAGTAAGATAATTATATAAATGTGATTGCAGGTGAGGTGACAAAGTGAAAACTGTCATATGATTATACTG

ATGACCCACCAACATGTGATAATATTGGTATTGAAGTCCCAAAAAGAATAGCTGCCCCAGCAAAAGTAAGCATATAATAG

TTTGATTTTGAGTTCCTTATGTCACATATCAAGTACATTGTACAAGTGAAGATATATGAGGAACATGTTGTATTTGCTTT

TTAGAAGAAACAAAATCACCCCCAGCAGCCATCCCTTACGCTGCTGCTAGAGCCTCCAAGAGGCCTATGAGTATAAAAGG

TTTCTGTGGTAGAGAAAGCATCTCCTCCATAACAGAGCCAGCTTAGCCCTATTATAACACAACGGACTGCA

bPb-2744

tgcagaagttacaattgggacaatggtgctgtggagacgaccatgaccactacgtgcgtggcaaacggggaagaggagca

tggacctccactacgtgcatacaccggtggcgtgccccgggatttagacgggagggagaggggcagaacatggccgggtc

atggtcttgcaccaagcccacacagcgcccacacttacttacatggaagcttcataacttaataatgtaattaatacttg

acggatcattgtttctggatttttctttcagcgtacgtaccagtaaaataaattgtaacttagaacagtactgcactacc

tttttcatgagtatactagatattgcttccttcgtttaaaagtatttgtcatagaaatgaatgtatctagatgtatttta

gttttaattatttttatttctatacatttgcgtgacaggtaattccggacggagggggtatttttttagcaaagtatagt

aagtgttgtttgtttaactgaaggtcgtttttgcgatttgtcctctttgtaaccgactgca

bPb-7965

tgcagccacagcgtttccaaggccctcggtacagaccccagcagctcatcagccccaaggccacgttcatgtcaccaaga

acgagcttgtctgtggcgtcgtcgggttcctcgcggcatgtttcctcctctctctgctgatcgtcatcgcaaagaaggcg

aggaatggggcggcgccggcgaccgatccggagccgcgagacgatgaagataccgtgctgaaggttgctgagcggtgcct

cggctgcttctttaaaCttCtgttatgcgcgtgatgggctcggtcgattgttagcagtgcaatttaccttgttgttggat

tgtatcattgtacgcactgctactgtagcagtggtctggtacactgccccttttattctttcaccttgtgtttgtagtat

cgctttataatccttgagcttacacctcggagaaatctgtaactgtggttgttgttgaataaatcttactgcactggcaa

tactactaatattgTttctatattaatcgtaTttgcagcttgTgcttttcttgGTgtaaacagtcgtgcactgtcctact

gTttgcatcgcttttatctctatatagccTatgctaatttccacGTtatcttcctaaaaattactcGtatctGggattct

attacactgtggagatccagactccttaaccatgttcaccaaccctgatcaaatcctgaatttgtctgtccttcatcact

acaataatcttattgtttttggatgtaaaagcaactcttaactttttcttaattaataaatgaggtaaatcttttgcctc

cgttttttttaaagctgttattgtgcgctgctaagccttagcaccgggcgcccggtgctgctctgtgtttctgaccggtg

gccggtgctagcg

bPb-1264

ttgggccctctagatgcatgctcgagcggccnccagtgtgatggatatctgcagaantcgcccttgatggnnccagtgca

gTccattgttgcatatggaccaaactagtggacaggagcagcaacctcaccgtcagctcaGGGtaagaaagattaagctg

attttttgctgtcacacgcattccgatttaatcgtcggcagtttcaaccgtataaaaatcatccaaatgactttgaagtt

cagcagaaaactactgtacaagccaagaatgcaacctcctgccgtatgcgttcctgcaattaagagtgtgattgacaagc

actgcaaggacgacgagtcctctgcctcatatgaaccactataaataactggacagcagcagcaacgtcgctctcagggc

aagaaaagtccagtatctgtcatcctcgaatttctacagcaataattccagacatgtaaatgtcaaaacgacttgtagac

cgagaacaaattgctacggcattcccatttaatctgcgttaattttagccatataaataacattcagatgctttcgcaaa

aaatgaaataacattcaaatgacttggaaggccagcataaaactacaagctaagaatttgacctcctgccgtatgtgttc

ttccgagtaccagtacagctgca

bPb-44182

tgcaggaaatactgatacagagataaatcaccagaaaatcagaataaggtaaccaacagattgctgtatttttaactatt

attacaatccaagtaaagtttaatgtgattctgattacaaatatctagtactagtattacatggcgccggatcattattc

atacatcagatgatcatgtagtggaaagaccgaaatatattgcatgctagtgatatgcattgatgtgcaacagatggacg

gagcagtacatgcccccagtatttactaaattgtggcagcatgttttccatgatttacaccacagggtaattttcagatg

tgatgaatggctttgggggcatctgtagattctgcaacctcgcggttaacatgtttaccacctttgtcattgatggctga

ttcttcgggttccactggatacaccatagtcccacaatggcaagttgtctgaccttttcttgctcttcctgtgcccagtc

caaagtaactactaagtcctgcccactgcttactttctgatagatccactccgggagatacacatcgttctggctaccaa

cacttgggtccgagttcctccttctactcaccatttctaacactagcatgccaaaactgtatacatcagatttgtatgat

actcccccaaagttccgtgagtatagctccagggcaatgtagcccatcgtgcctcttgctgca

bPb-3689

tgcagagcttttcacagagtggaaggtacatgtaattaagggtcggctggcaaagggcacggtataccgacccgacatag

aagagcccgaagagggtaggtggcatgccacaacactcgctgaccaagctaatgtttccttggcgccaagtgttcgtttg

gatcaggaTgttctaattgccatagtgggccgctaagacaaccaaaagcatcggatatggggaaatcccgaaaatggaca

ggaattcagcgtgcgatgagttttgatcgagccacgtgtcaagccagaagagggtaccgacaacatcacgtatgaagaaa

gtgatacccaagtgaatcccattgttgatcgactggggaaatttctagaaccgatacccctcccacggatcacatgccag

aagagggagaccctgtgggtatttggcttttataagctgca

**bpb-7624**

**bpb-3025**

**bpb-2891**

**bpb-1799**

**bpb-2531**

**bpb-5977**

bPb-8515

tgcagtacaacggtagggacgctgacggtaagattgatgaagaattcgaaGataaaccgataaataatttttatatagaa

caaagctaatctagggttaatatgtactccctccatgacaaaaatataagagcagttagatcactaaagtagggatctag

acgctcttatatttttgtacacggggagtagtagaatcctacgaccagtcaaggactatgtatcgtattaccttatttat

tcattctatccgtgtcactaaattattctgtttttggtagatgcaacatcgttaaatttgtgaacagaaagtagatgggt

gttgggccttctgctctttttggcactagaaaatgtaatattaaatatttgtttgctgtgcttggatatctattgctctt

gtcatccttcttgtaaatttttgtattaatgcccttatatatataaaggacacaccctctccgaaaatggcgtttggtgg

ccgagcaccctgca

bPb-3649

TGCAGAAACGAAACGATTAACTGGGTCACAATGCTCATAACTTTTAACATAAATAAGATGGCGAAAACAGAATATACTAC

ATAACAAGAAGATAGGTTAGTGCTACACATTTCATTATTTTATCAGCTACCATAGGTATTACCCCTACAGAAGAAACTCT

CTTGGCACGTAGAACTACATTCAATGATAGTTCATCAAAGTTTTTCACTTGCTCAAGTTTAAAGACAAGTCAATATGAAT

ATGTACATATAATAATTTGATAAAAAAAAAGTGTACATATACTTAGTGCATTTACCTGATAGATCTCAGTCGTCAAGTGT

TCGAAACAGATGGTGCAGTCAAGGGTCTCCTTCAATATTCTGACATCCATCCNAGTGTCTCCCTCTCTGATAGCTTCAGC

ACTTCTTTTGTTGGAGCTGCTGCCGTTCATCGCCNAAAAAACTCAAGGGTACCAGAACTAANTTGCTGGCTGTGTGCTGC

TGGTACTACNGACCGAAACCAAGATCCNAACCTCTTGATCCACACAACCTCACCGCTCCNTTCTCCGCCCTGTACCACAA

GAGCACATGANATACTGATAAAATTC

bPb-8441

GATCCAGTGCAGCGCAAACATTTGCGTTAGATTTACCAAAAAATCATTGGAAAAAGTGCTAGAAAATCCAAAAAATTGAG

TGCAACGCTAAAGAAATCATGCGGATACGCAGTCTACGAAAGGGCGAATTCCAGCACACTGGCGGCCGTTACTAGTGGAT

CCGAGCTCGGTACCAAGCTTGGCGTAATCATGGTCATAGCTGTTTCCTGTGTGAAATTGTTATCCGCTCAA

bPb-0945

TGCAGCGAGTGACCATCCCAACCGCCCGGATCTTCGTATCCGATACATCGAAAAAGGTACACAACTGCACCTCCACGCAT

CAAATCATGCGCTTCCTCCCATTCCTGTGATCCACCACGTGGAAAATTCAATGCTCATTCCTCTGATCCTCTCTCTATTT

CTCAATTTAGGCAAAAAGAACTTGCAGGCGCTGAGGTTTGCTTCTTGTGCATGGGATTCCTCTGTTCTACACGCATTGTT

TTGTTTGCCGCCGATTGTTCAGATTTTCTTTTCATCAAGCAAGTGTTTAGCATTCAACTTCCTTTTTCCTTTCAATTTCA

GATCAAGGGCTCGAGTTATCGAAATTATCCTGCA

**bpb-1852**

bPb-6322

catggatgtagaagcAcTacAAatccatggcacacggcgacaaccatacaGgATTgatGtcactgccAagaagctgcact

ggatccggatacgcagtctacgaagggcgaattccagcacactggcggcagttactagtggatccgagctcggtaccaag

cttggcgtaatcatggtcatagctgtttcctgtgtgaaattg

bPb-6127

tgcagaaacgaaacgattaactgggtcacaatgctcataacttttaacataaataagatggcgaaaacagaatatactac

ataacaagaagataggttagtgctacacatttcattattttatcagctaccataggtattacccctacagaagaaactct

cttggcacgtagaactacattcaatgatagttcatcaaagttcttcacttgctcaagtttaaagacaagtcaatatgaat

atgtacatataataatttgataaaaaaaaagtgtacatatacttagtgcatttacctgatagatctcagtcgtcaagtgt

tcgaaacagatggtgcagtcaagggtctccttcaatattctgacatccatccaagtgtctccctctctgatagcttcagc

acttcttttgttggagctgctgccgttcatcgccaaaaaaactcaagggtaccagaactaaattgctggctgtgtgctgc

tggtactacagaccgaaaccaagatccaaacctcttgatccacacaacctcaccgctcctttctcCgccctgtaccacaa

gagcacatgatatactgataaaattcatggatttggaagtgacactgcactg

bPb-7199

cttccaaatccatgaattttatcagtatatcatgtgctctTgtggtacagggcggggaaaggggcggtgaggtTgtgtgg

atcaggaggtttggatcttggtttcggtctgtagtaccagcagcacacagccagcaatttagttctggtacccttgagtt

tttttggcgatgaacggcagcagctccAacaaaagaagtgctgaagctatcagagagggagacacttggatggatgtcag

aatattgaaggagacccttgactgcaccatctgtttcgaacacttgacgactgagatctatcaggtaaatgcactaagta

tatgtacactttttttttatcaaattattatatgtacatattcatattgacttgtctttaaacttgagcaagtgaaaaac

tttgatgaactatcattgaatgtagttctacgtgccaagagagtttcttctgtaggggtaatacctatggtagctgataa

aataatgaaATgtgtagcactaacctatcttcttgttatgtagtatattctgttttcgccAtcttatttatgttaaaagt

tatgagcattgtgacccagttaatcgtttcgtttctgcactggatccntcaagggcnaattctgcanatatccntcacan

ctggcggcngctcgagcatgcatctngagggcccaattcgccctatagtga

**bpb-7448**

**bpb-6978**

bPb-6468

tgcagcacgacacatgttgtcaaaccatctcttctttagttttctgtaacaagactcatgttgcacaaccttgttccata

attttctacgacaacggtgttgttgcaaaatttacaataacatcTccattaaatctaattttgtgtaacaagaTccatgt

tacaaaaattacaacattattttcagctgcccacgcagttgcgtagtcgtcgttcggtgtcatcattctgtaacaacatc

gttgttacaaagataagatgagagagagatatgtggtCcaataggacacatgtcaagcaacgcaggcggtggatgccaca

tttttgatcggccggttgatcataatcatttctcatatttttattgatctgatgtatgaccagcataaaaccattttcct

gaaaatgttcatcaatatcgttgcgtgatggtctagcatacaccatcatgtgcccattctataacaggcaataaattggt

catcgtctgtcctctgtgcgccgtccatgcatgatgatgtaccattttaatccttttatggacctactgctgcgccatac

gcgagaacgaactagaagttgagaattagtggtcctagtatggatgctcatcagtcatcagcggcacgatcgaccatatt

ccagcagctcatgagtcatgagaagctcctgtcctactcccgagccaatatatagttgaatgcatacgcatgacatctct

cgactctagttgattcgccctctgtgaatcatcacggtcggtaaagttcacctgaatttcacgggaccagcgtgtccgct

acggccggtgtcatcactcacggcgagtcCgGagctggcgaggaacctaactgtcgcaatcttccaccgacctgatcgat

cagtcagtcacaccggttcttctgtggttcagatgcgccttcgcctctgca

bPb-48285

tgcagctcgtgtccacaggaagtaaggaaaccactaggatgcgtctccagcacctcacctgcccgatccctctcgtatgc

attgcgctcggatgaccagcatcaccaaatcagtggctgagctattggtgggtggcgcgctatgctcgtatctccggtgg

ttcacagaggaggacgtcaactcccaaatttcttctctgcataagcaggtgcgctttagtttcgcgccttggggacacga

cagcgcaatgttcttggtaacagtgccttgcgatcctcccgttatgattgacagcatgagcaaagagtagtacgttgtga

tgcccgcataataactgaatgccttttttttagacaaataactgcatgcttaaaaaccctcatatgccaccaccactctt

gaaaccagtggcggatgcaggcacaatttcctggtgtggcggaggtaactagcatatgtgtctcagctcaaatcatcaat

gaagaatcatgtagacctagcatacatgacacgagttaggtatggcagcagccaaagcttgccagattgctggcttcgcc

actgcttgaaactcccaatgtggaagttgtgcttataacaccatttatagtttttagttgaagtttggggataggggatg

caatgattggtaatccaccactgactggactgagtccttaaaactgca

bPb-7350

tgcagttaccaaggttgggcctgaatttgttggttgtaagaagggtgatcctgtatgtaatgagttggttgcttttccca

aacttgaacggttgattttcattgatatgcccaactgggaggcgtggtccttttttgaggaagaagttgttgctgctgat

gcaaggggagaggatggagccgctgaGattcaaaatgaggatgcccaatctccaaggttgcaactgctgcctcgtttggt

actgttgacactcgatggttgcccgaagctgagggctctcccgcgacaacttggaaaggaCaccacccgcctgaaggagc

tcaaattaattggtacaaacagcttgaaggCcgtggaggacctcccgttgctctctgaactccttaacattcagaaatgt

gaaggcctgca

bPb-8172

TGCAGTCGCAAATCTTACCATGACTTCGAGGCATATGCTTTCAGCAATCTGGAGGTACTCATCCCACAAATCTATAGTCA

TGACATAGTCAAGCATCCTTACAGGAGGCGTGCACTTCCCATAACTAGAGAACACAATGTTTCCAATAACATCCCGCTTG

ATGCAATCAACGTAGGCCCTGACACCTTGATATATGCGTTAGAACACATGTTCTCGCATCTAAACGACGGTGGAAATAGG

GTTCAAACAACACATCAGGGTAAAAGTAGTCGTCGTACAACATAATACGGCCATGTGTCATCTGTCGGTTGAACCAGTGT

AGTTGGGAACATGCTCCGCCACACGTTCCACATCTGCA

bPb-1814

tgcagggaaggaacagagaagcagccgccaaactaagtacgccacgggcttctgaaggtgaagaggaaggtgaaggtatg

aggaagtatcgatcaccccttcttgcaattgttacttatagaacacaatatccatgtatgcagttcatcatgactggagt

gtgtgctttggtgtctctgagatttactacttacttatttgggtggcatgcgaatgaaatgtttaaatgcagccgagcat

gtttctggggacaaccacatacagattggaagcgggctgcttgagatccagaacaaagaaggtcaaggtataaagaagta

ttgatcacacaaaccttgttgcaattgctacttatagaacacattttccatgtatgcactttcatcatgagtagaatgtg

tgcctcattatctgtgatttatttgttacatttcgagcaatgtgcaagtgaaatgttcaaatgcagacaagtatgtttct

gcggacaagtatgttcagattggaggctgggagcaaggaaggaagaggaaagcaactgtgagtgtgacaccgaacggagt

gcaaccaattggaattgaaatccaGaacaaaacaagtcacggtataGaaaaaataatcactagctcttagtacccgcaca

tttcaaaatacacaatgagtttcttgataatcttttgtattattgcAgGtgagtgtctgaGaGctgaGaatcaGatggtg

aggctgcactggntccatcaagggcgaattccagcacactgg

bPb-2548

ttttcccagtcacgacgttgtaaaacgacggccagtgaattgtaatacgactcactatagggcgaattgggccctctaga

tgcatgctcgagcggccgccagtgtgatggatatctgcagaattcgcccttgatggatncagtgcagggaagGaacagag

aagcagccgccaaactaagtacgccatgggcatctgaaggtgaaggtatgaggaagtatcgatcaccccttcttgcaatt

gttacttatagaacacagtatccatgtatgcagttcatcatggctagagtgtgtgctttggtgtctctgagatttactac

ttagttatttgggtggcatgcgaatgaaatgtttaaatgcagccgagcatgtttctggggacaaccacatacagattgga

agcgggctgcttgagatccagaacaaagaaggtcaaggtataaagaagtattgatcacacaaaccttgttgcaattgcta

cttatagaacacattttccatgtatgcactttcatcatgagtagaatgtgtacctcattatctatgatttatttattaca

tttcgagtaatgtgcaagtgaaatgttcaaatgcagacaagtatgtttctgcggacaactatgttcagattggaggctgg

gagcaaggaaggaagaggaaagcaactgtgagtgtgacaccgaacggagtgcaaCCAattggaattgaaatccagaacaa

aacaagtcacggtatagaaaaaataatcactagctcttagtaCCCgcacatttcaaaatacacaatgcgtttcttgataa

tcttttgtattattgcaggtgagtgtctgagagctgaGaatcagatggtgaggctgca

bPb-2965

tgcagggaaggaacagagaagcagccgccaaactaagtacgccacgggcttctgaaggtgaagaggaaggtgaaggtatg

aggaagtatcgatcaccccttcttgcaattgttacttatagaacacaatatccatgtatgcagttcatcatgactggagt

gtgtgctttggtgtctctgagatttactacttacttatttgggtggcatgcgaatgaaatgtttaaatgcagccgagcat

gtttctggggacaaccacatacagattggaagcgggctgcttgagatccagaacaaagaaggtcaaggtataaagaagta

ttgatcacacaaaccttgttgcaattgctacttatagaacacattttccatgtatgcactttcatcatgagtagaatgtg

tgcctcattatctgtgatttatttgttacatttcgagtaatgtgcaagtgaaatgttcaaatgcagacaagtatgtttct

gcggacaagtatgttcagattggaggctgggagcaaggaaggaagaggaaagcaactgtgagtgtgacaccgaacggagt

gcgaccaattggaattgaaatccagaacaaaacaagtcacggtatagaaaaaataatcactagctcttagtacccgcaca

tttcaaaatacacaatgagtttcttgataatcttttgtattattgcaggtgagtgtctgagagctgaGaatcagatggtg

aggctgca

bPb-5487

tgcagcctcaccatctgattCtcagctctcagacactcacctgcaataatacaaaagattatcaagaaacgcattgtgta

ttttgaaatgtgcgGgtactaagagctagtgattattttttctataccgtgacttgttttgttctggatttcaattccaa

ttggttgcactccgttcggtgtcacactcacagttgctttcctcttccttccttgctcccagcctccaatctgaacatag

ttgtccgcagaaacatacttgtctgcatttgaacatttcacttgcacattactcgaaatgtaataaataaatcacagata

atgaggcacacattctactcatgatgaaagtgcatacatggaaaatgtgttctataagtagcaattgcaacaaggtttgt

gtgatcaatacttctttataccttgaccttctttgttctggatctcaagcagcccgcttccaatctgtatgtggttgtcc

ccagaaacatgctcggctgcatttaaacatttcattcgcatgccacccaaataagtaagtagtaaatctcagagacacca

aagcacacactccagtcatgatgaactgcatacgtggatattgtgttctataagtaacaattgcaagaaggggtgatcga

tacttcctcataccttcaccttcctcttcaccttcagatgcccatggcgtacttagtttggcggctgcttctctgttccT

tccctgca

bPb-6825

tgcagggaaggaacagagaagcagccgccaaactaagtacgccatgggcatctgaaggtgaaggtatgaggaagtatcga

tcaccccttcttgcaattgttacttatagaacacagtatccatgtatgcagttcatcatgactagagtgtgtgctttggt

gtctctgagatttactacttagttatttgggtggcatgcgaatgaaatgtttaaatgcagccgagcatgtttctggggac

aaccacatacagattggaagcgggctgcttgagatccagaacaaagaaggtcaaggtataaagaagtattgatcacacaa

accttgttgcaattgctacttatagaacacattttccatgtatgcactttcatcatgagtagaatgtgtacctcattatc

tatgatttatttattacatttcgagtaatgtgcaagtgaaatgttcaaatgcagacaagtatgtttctgcggacaactat

gttcagattggaggctgggagcaaggaaggaagaggtaagcaactgtgagtgtgacaccgaacggagtgcaaccaattgg

aattgaaatccagaacaaaacaagtcacggtatagaaaaaataatcactagctcttagtacccgcacatttcaaaataca

caatgcgtttcttgataatcttttgtattattgcaggtgagtgtctgagagctgaGaatcagatggtgaggctgca

**bpb-0527**

**bpb-7273**

bPb-0158

tgcagctggaggatggtcacctgctcctccgacggctacgtctccgcgctgtacgtccttctccccgcccatccttgctc

acgccggtctctctctccgtcgaggagcgtccgtcgtcggtgtcctcgtcttatttccggactaatcttctcatgtgatg

gtctgtgctcacagcggtctgcccagccaaaggctctccggtaaactgtcgcccggcatcgggaacctcacgaggctgca

atctgtgtaagctcgtcgctcagatttatcagtacagtttctttgaaaaatgtattggacagatgagagattgcaaagtg

ttcaacttcaacatctctcctgttcttggacaggctactgca

bPb-9746

tgcagttttagccaaattaagtttaaccactcagttctgcttgcagttcatacgcatgtacagttgcaagtgctatcaaa

cttggatctgggggtagagagaggggaaactcacttgcacaagaggaggggagtaggaagtagcactggaaaggctcatc

tgtcctgaccacgtgaggaggacgcagccatcagtcagctcgtccatacacgtgctcatcaggatcaccctcctctccca

gcatggcctagaacccgaccctcagtatctagcttgtccactgggacgattcaacaaaatgatttttctactataataac

gattgttgtagccatattttttttactttatataaactttgcccctttcatgaacaaacttgcccccatatgcttgcatg

ctggcttcgttcctgcttgtccatgcaagaagacttgcaccaactgtattacaaagtaatgataataaatatgcatattt

tggcacgagtatttgctaactaatgaactcacatctttactctactgatgactattcaaaagtgtgcatctccatctact

gtatgaaagccttattttaatctgttggaggttgagaccgactaactccgtgcctgCctaccaaccaagcttcaaccgat

cattatagtcagtcaactctgataattttgagtcatccgaaagaccgaaattgaacaaaaacgtgattattttacatgca

atGggatgcacaaacagagacacacccagttgactcgggccaattttaagtcatccgaaattgaacaaaaacgtgttgct

cacacgcacagtttgctgcttcagtttcagtcatggcatccacaattcttatgcatccaaaactgaaggtacaaatagca

ggcaggtgatattcagatctacctgctgcacctacagaggcgagatgaggtccaagaacccgcaacagcttctccattgc

cttccttgacacaacagcaggcactcacgcatcagagaaagaagaaagccagaggatgaagaaaaccagaggaagaaaag

gtgaggagagatccataccgaggctggctcttcttggaagggatgctgacctgcggccccatgggtggggagcagtgggg

ctagggttccgggcggtggttgaggttggagggggtggaaggtggaggaggaggagaactcctctgccgatgcgtcgtgg

tggagaaggaaggagctccagccgagccggcggtggccttctgctggtggctgca

bPb-6347

tgcagttttagccaaattaagtttaaccactcagttctgcttgcagttcatacgcatgtacagttgcaagtgctatcaaa

cttggatctgggggtagagagaggggaaactcacttgcacaagaggaggggagtaggaagtagcactggaaaggctcatc

tgtcctgaccacgtgaggaggacgcagccatcagtcagctcgtccatacacgtgctcatcaggatcaccctcctctccca

gcatggcctagaacccgaccctcagtatctagcttgtccactgggacgattcaacaaaatgatttttctactataataac

gattgttgtagccatattttttttactttatataaactttgcccctttcatgaacaaacttgcccccatatgcttgcatg

ctggcttcgttcctgcttgtccatgcaagaagacttgcaccaactgtattacaaagtaatgataataaatatgcatattt

tggcacgagtatttgctaactaatgaactcacatctttactctactgatgactattcaaaagtgtgcatctccatctact

gtatgaaagccttattttaatctgttggaggttgagaccgactaactccgtgcctgcctaccaaccaagcttcaaccgat

cattatagtcagtcaactctgataattttgagtcatccgaaagaccgaaattgaacaaaaacgtgattattttacatgca

atggatgcacaaacagagacacacccagttgactcgggCCaattttaagtcatccgaaattgAacaaaaacgtgttgctc

acacgcacagtttgctgcttcagtttcagtcatggcatccacaattcttatgcatccaaaactgaaggtacaaatagcag

gcaggtgatattcagatctacctgctgcacctacagaggcgagatgaggtccaagaacccgcaacagcttctccattgcc

ttccttgacacaacagcaggcactcacgcatcagagaaagaagaaagccagaggatgaagaaaaccagaggaagaaaagg

tgaggagagatccataccgaggctggctcttcttggaagggatgctgacctgcggccccatgggtggggagcagtggggc

tagggttccgggcggtggttgaggttggagggggtggaaggtggaggaggaggagaactcctctgccgatgcgtcgtggt

ggagaaggaaggagctccagccgagccggcggtggccttctgctggtggctgca

bPb-6329

tgcaggggccggcagggccgggcaagaaacaacatatgtgaactggggataagcgctggagagaccccacggccaccatt

ggctggccgacgactccctcaaggggggagtaacaagataatgaggttttgttggcgaggagagcgacatgggaggcgta

tgatcggaatcaaaacaaatagtcacatgtgtgctgttataattacgagagttgatcctgaatacagaaaataatttatt

gtgacattcagtacgaacatttgatatacattgttgctaaagatctgtgatctctgca

**bpb-1710**

bPb-6914

caTgatttgttttgttctggatttcaattccaattggttgcactccgttcGgTgtcacactcacagttgctttcctcttc

ctttCTtGCTCccagcctccaatctgaacatagttgtccgcaaaaacatacttgtctgcatttgaacattttacttgcac

attacttgaaatgtaataaataaatcacagataatgaggcacacattctactcataatgaaactgcatacatgcggatac

gcagtctacgaaagggcgaattccagcacactggcggccgttactagtggatccgagctcggtaccaagcttggcgtaat

catggtcatagctgtttcctgtgtgaaattgttntccgctca

bPb-3317

TGCAGATATACGTGGACGGTTGCGCTGTGTGCTGCCTGCGAGTCCCACATCGGCTGGCTGTTCAGGGCTGACAAAAGGAA

CCTTCTTCCGATATCCTTCTGGGGGATCCGCATTTCCCAAACCTCAGACGGTACACAATCGGCTCAGGACCGACGTTCCG

TGTAAGGCAGCGGCCGCGGTTCTTGGCGCGATTCTCGTCATCCTTCAGCAGACCCGTGACCTATCTTCGGTGGCCTGCTC

GTTGTAGACCTTGCTGGATTTGGCCCGACGGTTTCCTGCTCGTTGTAGACCATGCTGGATTTGGCCTGACGGTTTCATCT

AGTATTCCCACTGCA

bPb-9130

agtgtgatggatatctgcagaattcgcccttgacccagtgcagtatgtaggtgcagaactgattgcatgaatggaccgca

aagagataaaacatgccgcaaatttgttctgaagcacaagagtaaaacaaattgcacttcgaaattgcgttgttccGTtC

AacaaaaggaaaatatgcatggAAccatcaaaagaaactcaacaggtgccgccgcagtgaacagcaaaagcacttattta

tccaacatactcctacacaaccttggctcaaattgcgcacatctatcatcaaagatgcacaggaaagataacaacgaata

gtggttatattcatggaagaaattctgatatgaatcaagttttctgtatctatgcagatatgaatgaaagaattccagat

atcaatctgtttggaggtcgacaaaaattctgacaccaaagatctccatccaatctactctaagattatctgtctgttgt

atcatccgctggctctctttggcatggatgtgcctgagcttcactccagactccagagtctcgggtgatagtgtagctcc

tctgcgctacatcatggtgatgtctgtgaaatgaaacagcaaattaaatccaattagtgacaatatcacagccattaact

tgattaagattcgaaggatgcaggtacggataacaagaacaatcaaaggctgca

bPb-2646

tgcagccaccagcagaaggccaccgccggctcggctggagctccttccttctccaccacgacgcatcggcagaggagttc

tcctcctcctccaccttccaccccctccaacctcaaccaccgcccggaaccctagccccactgctccccacccatggggc

cgcaggtcagcatcccttccaagaagagccagcctcggtatggatctctcctcaccttttcttcctctggttttcttcat

cctctggctttcttctttctctgatgcgtgagtgcctgctgttgtgtcaaggaaggcaatggagaagctgttgcgggttc

ttggacctcatctcgcctctgtagatgcagcaggtagatctgaatatcacctgcctgctatttgtaccttcagttttgga

tgcataagaattgtggatgccatgactgaaactgaagcagcaaactgtgcgtgtgagcaacacgtttttgttcaatttcg

gatgacttaaaattggcccgagtcaactgggtgtgtctctgtttgtGCATccattgcatgtaaaAtAatcACGTttTtGt

TCAATTtCGGTCTttcggatgActcaaaattatcagagttgaCTgActataatgatcggTtgaagcttggttggtaggca

GgcacggagttagtcggtctcaacctccaAcagattaaaataaggctttcatAcagtagaTggagatGcacacttttgaa

tagtcatcagtagagtaaagatgtgagttcaTTAGttagcaaatactcgtgccaaaatatgcatatttattAtcattact

ttgtaatacagttggtgcaagtcttcttgcatggacaagcaggaacgaagccagcatgcaagcatatgggggcaagtttg

ttcatgaaaggggcaaagtttatataaagtaaaaaaaatatggctacaacaatcgttattatagtagaaaaatcattttg

ttgaatcgtcccagtggacaagctagatactgagggtcgggttctaggccatgctgggagaggagggtgatcctgatgag

cacgtgtatggacgagctgactgatggctgcgtcctcctcacgtggtcaggacagatgagcctttccagtgctacttcct

actcccctcctcttgtgcaagtgagtttcccctctctctacccccagatccaagtttgatagcacttgcaactgtacatg

cgtatgaactgcaagcagaactgagtggttaaacttaatttggctaaaactgca

**bpb-3113**

bPb-3278

tgcagaggcgaaggcgcatctgaaccacaaaagaaccggtgtgactgactgatcgatcaggtcggtggaagattgcgaca

gttaggttcctcgccagctccggactcgccgtgagtgatgacaccggccgtagcggacacgctggtcccgtgaaattcag

gtgaactttaccgaccgtgatgattcacagagggcgaatcaactagagtcgagagatgtcatgcgtatgcattcaactat

atattggctcgggagtaggacaggagcttctcatgactcatgagctgctggaatatggtcgatcgtgccgctgatgactg

atgagcatccatactaggaccactaattctcaacttctagttcgttctcgcgtatggcgcagcagtaggtctataaaagg

attaaaatggtacatcatcatgcatggtcggcgcacagaggacagacgatgaccaatttattgcctgttatagaatgggc

acatgatggtgtatgctagaccatcacgcaacgatattgatgaacattttcaggaaaatggttttatgctggtcatacat

cagatcaataaaaatatgagaaatgattatgatcaaccggccgatcaaaaaagtggcatccaccgcctgcgttgcttgac

atgtgtcctattggaccacatatctctctctcatcttatctttgtaacaacgatgttgttacagaatgatgacaccgaac

gacgactacgcaagtgcgtgggcagctgaaaataatgttgtaatttttgtaacatggattttgttacacaaaattagatt

taatggagatgttattgtaaattttgcaacaacaccgttgtcgtagaaaattatggaacaaggttgtgcaacatgagtct

tgttacagaaaactaaagaagaggtggtttgacaacatgtgtcgtgctgca

bPb-2406

tgcagaggcgaaggcgcatctgaaccacaaaagaaccggtgtgactgactgatcgatcaggtcggtggaagattgcgaca

gttaggttcctcgccagctccGgactcgccgtgagtgatgacaccggccgtagcggacacgctggtcccgtgaaattcag

gtgaactttaccgaccgtgatgattcacagagGgcgaAtcaactagagtcgagagatgtcatgcgtatgcattcaactat

atattggctcgggagtaggacaggagcttctcatgactcatgagctgctggaatatggtcgatcgtgccgctgatgactg

atgagcatccatactaggaccactaattctcaacttctagttcgttctcgcgtatggcgcagcagtaggtctataaaagg

attaaaatggtacatcatcatgcatggtcggcgcacagaggacagacgatgaccaatttattgcctgttatagaatgggc

acatgatggtgtatgctagaccatcacgcaacgatattgatgaacattttcaggaaaatggttttatgctggtcatacat

cagatcaataaaaatatgagaaatgattatgatcaaccggccgatcaaaaaagtggcatccaccgcctgcgttgcttgac

atgtgtcctattggaccacatatctctctctcatcttatctttgtaacaacgatgttgttacagaatgatgacaccgaac

gacgactacgcaagtgcgtgggcagctgaaaataatgttgtaatttttgtaacatggattttgttacacaaaattagatt

taatggagatgttattgtaaattttgcaacaacaccgttgtcgtagaaaattatggaacaaggttgtgcaacatgagtct

tgttacagaaaactaaagaagaggtggtttgacaacatgtgtcgtgctgca

bPb-2630

tgcagaggcgaaggcgcatctgaaccacagaagaaccggtgtgactgactgatcgatcaggtcggtggaagattgcgaca

gttaggttcctcgccagctccggactcgccgtgagtgatgacaccggccgtagcggacacgctggtcccgtgaaattcag

gtgaactttaccgaccgtgatgattcacagagggcgaatcaactagagtcgagagatgtcatgcgtatgcattcaactat

atattggctcgggagtaggacaggagcttctcatgactcatgagctgctggaatatggtcgatcgtgccgctgatgactg

atgagcatccatactaggaccactaattctcaacttctagttcgttctcgcgtatggcgcagcagtaggtccataaaagg

attaaaatggtacatcaccatgcatggacggcgcacagaggacagacgatgaccaatttattgcctgttatagaatgggc

acatgatggtgtatgctaGaccatcacgcaacgatattgatgaacattttcaggaaaatggttttatgctggtcatacat

cagatcaataaaaatatgagaaatgattatgatcaaccggccgatcaaaaatgtggcatccaccgcctgcgttgcttgac

atgtgtcctattggaccacatatctctctctcatcttatctttgtaacaacgatgttgttacagaatgatgacaccgaac

gacgactacgcaactgcgtgggcagctgaaaataatgttgtaatttttgtaacatggatcttgttacacaaaattagatt

taatggagatgttattgtaaattttgcaacaacaccgttgtcgtagaaaattatggaacaaggttgtgcaacatgagtct

tgttacagaaaactaaagaagagatggtttgacaacatgtgtcgtgctgca

bPb-9336

tgcagatggcgatgtgtattcctttggaggcaaccagttcgggcagctggggattggttctgatcaagctgaggtgggaa

ctttgagtcctttttccattcttattacataacataatccgattcagggatactaagtgtgtagctgtctcttcacttgc

ccaaatacataagaatgttacttctgttttgatgcagactataccaaaactggtggatgcccccagcttggaaaataaga

atgcaagatcagtgtcgtgtggagctcgtcatagtgcaataataacaggtatagcggtcgtattttcttctggaagtaat

gttttattttatttcttgattctaacatttttttacgaaatttctgca

bPb-18973

tgcagattcagtggctgttgcgagggaatgagctcgcagtggtgatggcgctcggtgtattttgtgaaaaggagatggaa

gtgtgtgggtgggtgtgagctaaaagtgacggcgagtcagcaagtgtggccgagaatccagagcaattgattggcgtcgt

gtcacggtcagtactgtacttagctagtaatgcggggttggacgggaatttaataagcacacgcttaagtttgccggtca

aagagggtttgatgaatccggatgcagagatggctgcggaagcatcgtgaaccagattaattccccatctgtgtggactg

atgctgca

bPb-0619

tgcagattcagtggctgttgcgagggaatgagctcgcagtggtgatggcgctcggtgtattttgtgaaaaggagatggaa

gtgtgtgggtgggtgtgagctaaaagtgacggcgagtcagcaagtgtggccgagaatccagagcaattgattggcgtcgt

gtcacggtcagtactgtacttagctagtaatgcggggttggacgggaatttaataagcacacgcttaagtttgccggtca

aagagggtttgatgaatccggatgcagagatggctgcggaagcatcgtgaaccagattaattccccatctgtgtggactg

atgctgca

**bpb-2137**

**bpb-5396**

**bpb-5298**

bPb-4156

tgcagttacacaatgtcgacaagtatgctccgctgcaccttgtgttctggaactgattgtgcaaccgtgccgggcagttt

cttctgatggatgcgacttctcaggtttggttggggataggatagagattacccatgtttcgttcgtacagctcctattg

tttttgtgtcgagtcgcgggatggtcatttatttgtattagctttgacgcacttcttctgcttgtatcttattagtgcga

acatggtttagcatgcagcaaaagtatcaggttattcagaatttcagagCccatatattattgttctcattttgttccta

gaagtggaaaccatgttgttcatgtatatactgctatttttgtcagtcatgcacccatcttGcaaatcagctcaaaaCat

tttcaaaccgatctcatttgattgtgggagtgacagacaatgaatatcatggaaggaaacctcagtcaggggtgacaggt

gagcaaagAaaAcagaagtactccctccgtcctacaatataaaagcatttttgacagtatagtagtgtcaaaaacgctct

tatattATgggaTgGatggactgttttgtactggagaatgcttcacattaccaaacgaacatttacAgagcatcttccag

tccagcgtggtggtgttatcccatctctgaatgcagaaacaatgaagcagagaggggagtgacaggctgcgtgcgtggtg

ctactatgataatacataacttctttacatgggtttcactgggcgctgtgtcttcttcgctcgtccgacagaccgggctg

ctgttgctgca

bPb-2420

TGCAGTACAACGGTAGGGACGCTGACGGTAAGATTGATGAAGAATTCGAAGATAAACCGATAAATAATTTTTATATAGAA

CAAAGCTAATCTAGGGTTAATATGTACTCCCTCCATGACAAAAATATAAGAGCAGTTAGATCACTAAAGTAGGGATCTAG

ACGCTCTTATATTTTTGTACACGGGGAGTAGTAGAATCCTACGACCAGTCAAGGACTATGTATCGTATTACCTTATTTAT

TCATTCTATCCGTGTCACTAAATTATTCTGTTTTTGGTAGATGCAACATCGTTAAATTTGTGAACAGAAAGTAGATGGGT

GTTGGGCCTTCTGCTCTTTTTGGCACTAGAAAATGTAATATTAAATATTTGTTTGCTGTGCTTGGATATCTATTGCTCTT

GTC

**bpb-8021**

bPb-8557

TGCAGATTAGCAAGAAACTCTGCTTTCTGTACTTCAACTATTACTCTGTTTTCTAACATTCAAGGGAGGAATTGACATGC

CTGTACTTCCACATGGCATTGTACTTGCGTGAACTCTTGTTTTCGTTTGCAGCATTTAACTGAACCACTTGTATGAATGT

GCTTCCACATGGCATGCTAGGTAGCAGGAGTGAAACACATTTCACTGATATGGAATAGATCGATCGTGCGAGAGCTCACA

GTTCCCTCTGCCGCTTCCATTGCAACCAAGACGGTCAAGAATACTGTATTTCTCCGATGCCGTTTGATCGTCTTACTCCT

GCA

bPb-5374

tgcagagacatacacaaagcgcccgcttgctcgacatggaacaactacaagaccagccataatcttgtcatgcatgctcc

cgcagccattaacgcactgtctaAcatcactatatggtttttaaatgactcgtcccagtttttagtaaatgcaaatacca

tgaaattaatgaagcAccataaaatgatgagatgctcttgtacaagatgatagacgtttctataaatttgtagatactgg

atggacagctaatgcattcaccaacctttcattgtgtaaccctgacccgcattataaaagggtgcactaccttgtatttt

cttcttggactagggcacaaactagaaagaggcgtgggttggaatgggacagttcattatgaatgagagacatacgggaa

cacctcatccattattcctgacatggatgtagaagcactacaaatcaatggcacacggcgacaaccatacaggattgatg

tcactgccaagaagctgca

bPb-7689

tgcagagacatacacaaagcgcccacttgctcgacatggaacaactacaaggccagccataatcttgtcatgcatgctcc

cgcagccattaacgcactgtctaacatcactatatcgtttttaaatgactcgtcccagtttttagtaaatgcaaatacca

tgaaattaatgaagcaccataaaatgatgagatgctcttgtacaagatgatagacgtttctataaatttgtagatactgg

atggacagctaatgcattcaccaacctttcattgtgtaaccctgacccgcattataaaagggtgcactaccttgtatttt

cttcttggactagggcacaaactagaaagagacgtgggttggaatgggacagttcattatgaatgagagacatacgggaa

cacctcatccattattcctgacatggatgtagaagcactacaaatcaatggcacacggcgacAaccatacAggattgatg

tcactgccaagaagctgcactggatccntcaaggggcnaattctgcagatatccatcacactggcngnngctcgagcatg

catctagagggcccaattcgccctatagtgagtcgnantacaattcactggccgtcgttttacaacgtc

bPb-3623

TGCAGTGTGCTCCTCGGCTCTGACCTACTCATGTGCTGACTGGGCCAAATAGGCCTCCTCTTTGCGTTGTCACGGTCCAG

CTATGCTCGGGCTGAAAATGTTGTGCAATGTTGCAGCTTCTTGGCAGTGACATCAATCCTGTATGGTTGTCGCCGTGTGC

CATTGATTTGTAGTGCTTCTACATCCATGTCAGGAATAATGGATGAGGTGTTCCCGTATGTCTCTCATTCATAATGAACT

GTCCCATTCCNACCCACGTCTCTTCTAGTTTGTGCCCTAGTCCAAGAAGAAAAATACNAGGTAGTGCACCCTTTTATAAT

GCGGGTCAGGGTTACACAATGAAAGGTTGGTGAATGCATTAGCTGTCCATCCNGTATCTACNNTTTATANAAACGTCTAT

NATCTTNTACAAGAGAATCTCATCATTTTATGGTGCTTCATTAATTTCATGGTATTTGCATTTACTAAAAACTGGGGACG

AGTTATTTAAAAATGATATAGTGATGTTAGACAGTGCGTTAATGGCTGCGGCAGCATGCATGANAAGATTATGGCTGGTC

TTGTAGTTGTTCCATGTCGAGCAAGCGGGCGCTTTGTGTATGTCTCTGCACTGGATCCNTC

bPb-7827

tgcagcagagcatgatgccaagatttcccgcatgatgaggaaggaaggaaaggagacaagtgcatgcaagaccgtccaga

gctggggatccctgtcctgtctggttcaaccacggcaagaacccaaagcccaacatcaaacgccatgcactatggcgttg

gagtaagcagcggaaacgccagtccttatagcgtttcttgctttgacagagacgagattgttaggcgtataagcgtggag

cagaaacatcaaggtaaattgatgtttttacatgcagcagcaacgccatgggctgtggcgtttctaaaaaggtcagatcg

tgaaatactttcagaccgacttcattctgtgtattcttttcgtctcgtgggtcagaacagtgattttatccgcaggagtt

aaacacatttcattgatatggaatagatcgatcatgtgagagctcacagttccctctgccgcttccattgcaaccaagac

ggtcaagaatactgtatttctccggtgccgttagatcatcttactcctgca

bPb-5312

tgcagagagaaagatagagagagagaatcgtgcggtttaaacgggatcacctgcgctgttaattaatgagagagagatag

actaccggtttgagatgatcaccttgtgtatttatttgtttgcgggttatttctttctgttaattttgtttgacagcatg

gcagagagagagcgagggagatgagttagcagttttttttttttttttgagcgggagcagttgtgtacatggaaagttgg

aaacaaatcttgacggaatatgatcgatatttcttgaccaacaagatagtatcttattggaggcaatggaaagctaatta

aaaccagcgaagatagtaaagacagagagaaaccaacattcaataaaaacttcatgaaagaaataacatgtactccctcc

ataaattaatatggcgcgtgctaccaatcagccgactgatccatgtgtgggatcagccggctcggcggccgttagatccg

tccgttagatgcggtctgtagacttgcccatgtcgttatcacctaGccttttccagcccccaccacagttacttctccct

gtacactattctatatgacctacgagcaacactgca

77164

tgcagtagctgctaattttttttgatggtggaggggcacgtccatctttgatggttgcctgtgattttaggttCtaagtg

atatCtgatttgacatagtACTgatcctagttccccctatgtgtttttaTataggCgttttgcaaagAgtatagacAGgt

tgctcctccTgttcAgtcgaggTgtCtttttggatccCGagatggCtggcaagtagacAgacAaggagcgCgaaTgacct

ttttaaaTtttgtTgtcgttacttgtatcaGttTgtttttttttttttgcatggactgctgtaatgtttttcaggaattt

ctggacctaattgtctgatgcaatgttccatgaaattgaaatcacgttgaagctatgctgagtttgttgaccttataaac

atttggagtacagtctcacactggctggaacgctgattcattttttctcttctttttttccttttgagactgatgaaacc

gagttgaggctgca

bPb-3130

GATGGTCGCTTAGTGGTTACAGCGTACTATCTCGCAGCATAGCGTTGCAAGTTCGAGTCCCGTACCCGCTGCACTTTTTT

GATGATTTCTAGATCCGAAAAATATs

**-------------------------------------------------------------------------**

**Chr. 4**

**bPb-42695**

bPb-7645

tgcagctaccctagtgtgtgtgcgtgtctgtgtgtggctgagtttgccgtttcttggatggatctggccctctcccatgg

ccatgactcctcctctcttcttgccttactttcgtttgctgtcatcaacttttctccgtgtttcctttgtgttttgtctg

tgccgatgatctccgcgttcattttgaggcactcttgtgttgttcctgcgtttcacggggtgccggtgatggtttccttc

atgcatacntacataggttccatcttccatgtgatggttcaaccGcattatttaccagagGCgctactcgtgtttactgc

gtgcatttgctCtTgtgtacggagggttgtctgaTttttacAaggcacggccggtgatgaaacaacgatggcTttttgag

atgtgctatgcatgcatatgcatccacagatgacaagaccgaatcaatttcttcaaagcatctgaggccagctagccAGa

tcttttactgcgtttgctgtgatgaacttgtgggtgttagtctctccctgttcttggatctgtctcttCTttgcatcctc

tttttgacgtgcacgtgccgatgatccgcggtagtcgtttcttggatctggtcctctcgatcccatgcatgcatggccat

gaatataatcctttgatcttggctttcgtttgctgttgatgaactgtcctccatgtgtggaggtctgttttagttggccg

catgcaaagggttatgatgaacttttgctgtaccacaaatttcttgtgtaagaagggttgtttgatttttacgaggccgg

tgacgagacatcattggctttcgagatgtgctacatatgcatgcacagatgacatgagcgaatcaatttcttcaaagtat

ccgaggcctcccaatcccaagcacgacgatttactttggctttggttccatcctgca

bPb-14836

tgcagcactagcacttactaattgcaagattagaggccagacatTgccaattttcagtagcacctagcaggttaatacca

cacccttcagaaaataaacccaagtaggcaagtaccaccgctaccacgcttggcggcagcaccatttctcaccaactaat

taaaggaatctctcgctcatgagctaattgattaacaggtcaagagacgcagcgcgcacctgattccggtggccaacacc

atcgccgctgtggccgtcggagaagtggcactaggcacaggcggattccaaaccctgctctgctccctctggatcccact

atggtgatggcgaaagagaacaccattaatattaacctcctcttcttattgagacaacacacaagcctttgcacgtcgcc

cgaagaaaaatgccaagaataaactgtcacaaatctgca

bPb-9672

cgtcgcgggaggaagatttggacgctggacataccttgatcccggatacgcagtctacgaaagggcgaattctgcagata

tccatcacactggcggcagctcgagcatgcatctagagggcccaattcgccctatagtgagtcgtattacaattcactgg

ccgtcgttttaca

**bPb-2909**

bPb-40823

tgcagtggagatgaagcttagcatcaacaggatgatgccagcaggaagagacccctccatggcgacacggagtcaggaca

ccatcacttgctacgagctggaacaaccttctgccgctagtatgtaataaggcgccttgtaatatcatatgaatggacca

gccatatgttatttccttagaataaccagctgaactcccttgccccgctggatattattgtacaatccacttggacctct

ttatagttcctactagagatgaagcacgtataattttccatgtgggtttgcctttgatttgacaaacatgcaagtttcta

aaatgcattttctccagtatttaccagaattcatgcgtttttcttatttctgtcaaacatgtaagtttacaaaattccta

tattttctaaaagaagtcctctgttttgcacttgtatgtctatcatattcttgtgtctgtcctttcctatgctttcagaa

tcccgtgtaccaaagaggcccttttgattggaaacagagaccggtttcacattacaatttcaatcaacaagcttaatgat

ttctaattagtctgaagatcatcatccactgacggtgttgcccgtctccagtcattcagagctgca

bPb-1469

tgcagtagattagtgcaatgtcactagcattgtgccctgatcccacggccatccatccattattggtgcctaaagtctat

tcctctgtcgtgcattgcatgcttgatggtcctgatcagaagaaacaccatccactgatctgtataagaaaagtacgcgt

gccaacacaaacacagagagttggaccttgtcatcttataagtatgatgattcaccttgttggtcgtggcaaactgattg

ttctgtAccttattccaattcagaggttatactttttgtgcatgaaatacatatttaccgtttgattttagaggttttgt

atcacatctcctctggagttaatttcttcctgacaaaatgaagacacagttatacgcagcctagttacatactctgtgaa

aaacaaatccggcacagccaaggaaatagctgatataagtgatctccagaattgcctgacataactcatcatgtggaaat

tactccttttctgttcctttagtcagtcttttttgatagaaatgatggtgtcctgatcccacggCCAtccatttattggt

gcctgattttttgagatcctccttttttccggtgccgtgcattgcatggTtggctgtcccaatcacaagaacttccagcc

acagcttctgtgtaggaaaaatatgtacgccaagaccgagaccaagacttgaaaaataccgtaGaccatacacaataata

ttgtgcttacgagtgaagtcAggatgaactactctctccattccattgttgttctcatcctgtagctagctgctctcttc

ctcgtggtaccaatatggctgccgagcttgggcatctcgcccgagcagctgca

bPb-8569

cagtgcagtagattagtgcaatgtcactagcattgtgCCCTgatCccacgGccatccatccattattggtgCCtaaagtc

tattcCtctgtcgtgcattgcatgcttgatggtcCtaatcagaagaaacaccatccactgatctgtataagaaaagtacg

cgtgCcaacacaaacacagagagttggaccttgtcatcttataagtatgatgattcaccttgttggtcgtggcaaactga

ttgttctgtaccttattccaattcagaggttatactttttgtgcatgaaatacatatttaccgtttgattttagaggttt

tgtatcacatctcctctggagttaatttcttcctgacaaaatgaagacacagttatacgcagcctagttacatactctgt

gaaaaacaaatccggcacagccaaggaaatagctgatataagtgatctccagaattgcctgacataactcatcatgtgga

aattactccttttctgttcctttagtcagtcttttttgatagaaatgatggtgtcctgatcccacggCCAtcCAtttatt

ggtgcctgattttttgagatcctcCctttttccggtgccgtgcattgcatggttggctgtcccaatcacaaGaacttcca

gccacagcttctgtgtaggaaaaatatgtacgccaagaccgagaccaagacttgaaaaataccgtagaccatacacaata

atattgtgcttacgagtgaagtcaggatgaactactctctccattccattgttgttctcatcctgtagctagctgctctc

ttcctcgtggtaccaatatggctgccgagcttgggcatctcgcccgagcagctgca

bPb-2837

tgcaGATTTgtgacagtttattCTTggcatttttcttcgggcgacgtgcaaaggctTgtgtgttgtctcaataagaaGAG

gaggttaatattaatggtgttctctttcgccatcaccatagtgggatccagagggagcagagcagggtttggaatccgcc

tgtgcctagtgccacttctccgacggccacagcggcgatggtgttggccaccggaatcaggtgcgcgctgcgtctcttga

cctgttaatcaattagctcatgagcgagagattcctttaattagttggtgagaaatggtgctgccgccaagcgtggtagc

ggtggtacttgcctacttgggtttattttctgaagggtgtggtattaacctgctaggtgctactgaaaattggcaatgtc

tggcctccaatcttgcaattagtaagtgctagtgctgcactgaatccatcaagggcgaattctgcagatatccatcacac

tggcggCCGCtcgagcatgcatctagagggcccaattcgccctatagtgagtcgtattacaattcactggccgtcgtttt

acaacgtcn

bPb-7275

tgcagatttgtgacagtttattcttggcatttttcttcgggcgacgtgcaaaggcttgtgtgttgtctcaataagaagag

gaggttaatattaatggtgttctctttcgccatcaccatagtgggatccagagggagcagagcagggtttggaatccgcc

tgtgcctagtgccacttctccgacggccacagcggcgatggtgttggccaccggaatcaggtgcgcgctgcgtctcttga

cctgttaatcaattagctcatgagcgagagattcctttaattagttggtgagaaatggtgctgccgccaagcgtggtagc

ggtggtacttgcctacttgggtttattttctgaagggtgtggtattaacctgctaggtgctactgaaaattggcaatgtc

tggcctctaatcttgcaattagtaagtgctagtgctgca

bPb-9204

TGCAGTTCGGCGGTGACCATCAGGGATTCAGGGGCCTTCCCTTCATCTATCTTTCTGTCTCGCTATATTTGCTTTCTAGC

TTCATGTGATTAGGCCCATGTTCTACCCTTCTTCTGTGATTGTTTATAGTCACCGGCAGATTTGTATTACTACGGTTGAT

TTATGAAGCACTGTTTTAAGGTTCATGTCTCCCCAATGTCATCCAATCAAAGTCAAAAAGTCAGAAAATTGGACTAACAA

TAGTGTTTCGGTGAAAATACTAAACTGAAGAAGTGAAGAAATTCTAGCCGACCTAACATCGTGAAAGGAATCTTTCCCTA

ATAATAAAGTACACATCGCTTCTGTCGTACGTCGTCGGTCATTTTACATAAAAAACCTTGCGTTTTTTCCAAATCAACCC

GTAGCCCGGATATAAGTGACAAGAACGAACCGCTTTTTGTAGTTTTGCAAATAGGCCCTTCTGTTTTCATGGAATCAACC

CGCAATCCGTTTTGTCCGCATTTTATAGAAAAAACCCTAACTTTTACGATAAATAACCTGCA

bPb-10678

tgcagttctctgggctccaccttctccaaaaatgtaagacttctcaccacactttgggcacttgaagcaactcatgttct

ctactaagcctagaatctgccaaaatatacagctgagtaatgctgggaagagtggaagactttgtgaaatattggatagt

ggcacatgctagaaatttaagctaattatttttcactcgtgaaatgaatatgcctactgcgattgaattctcatgaatag

atatgtgacagtaataaactaacaatagtaactgatgctggttggcaataacattaacaaataaacagggaaatgaacgg

tgtgataggaattagaattctatctgaattaatgtttaatatctaatacacgcaaacggctagaaactaagaaagaagta

cagatatcacaaaacagaagttacatagaacatacaggaacttggactttacgaaacatatttgctcctcttctagcatc

aattagagcaatatcttgaggagtagaaacaattaaagcacctgca

bPb-6145

aattcgcccttgatggatccaTGCAggtgctttaattgtttCtactcctcaagatattgctctaatTgatgctagaaGAG

gagcaaacaTgtttcgtaaagtccaagttcctgtatgttctatgtaacttctgttttgtgatatctgtacttctttctta

gtttctagccgtttgtgtgtattagatattaaacattaattcagatagaattctaattcctatcacaccgttcatttccc

tgtttatctgttaatgttattgccaaccagcatcagttactattgttagtctattactgtcacatatctattcatgagaa

ttcaatcgcagtaggcatattcatttcatgagtgaaaaataattagcttaaatttctagcatgtgccactatccAatatt

tcacaaagtcttccactcttcccagcattactcagctgtatattttggcagattctaggcttagtagagaacatgagttg

cttcaagtgcccaaagtgtggtgagaagtcttacatttttggagaaggtggagcccagagaactgcactggatccntcaa

gggcgaattctgcagatatccntcnnnnnggcggcngctcgagcatgcatctagagggcccaattcgccctatagtgagt

cgtattaca

bPb-1999

tgcaggcagcagttagcgtgcattttgttaactgacgcgtgcagcgttcttaaatgtaccggccagtgtaagcgccgggt

cacaaaaattttcttcgaaattctatatacttttaaatttttgataaaattttgccaaatttcgacgaacatgttttaaa

attaatggaagtttctccgaaaatcggtacacctttaaattctatgaactttctcgaatattgacgagctattttaaaaa

tgatgaacattttcagaaaatggaagatcttttttaaaattgattaaactttttaaaatgtaaaaaacaattcagttcaa

tgaactttttttgaaaatacatgaacattttttgaatttctatgaacttttttcaaattaaatgacctttttaaaaattt

aataaacttttttaaaattgatgaaccttttttcaagatatgaactttttttgaaatgaatgaacctcgtttttaaatct

ggaaacccttatttatttcatgtttttattttcataaaaatagaaaatgtagacggtttttttcctactgtatcaacaga

gaacaaaaaaagaaaaaaaactgagctgctcaccagcgtgctagtaggccggcccaagccagctactctaatggcgccgg

attgttaacgggtgctgcactgnatccatcaagggcgaattccagcacactggcggcccgttactagtggatccgagctc

ggta

**-------------------------------------------------------------------------**

**Chr. 5**

bPb-0170

tgcagatagaggaaagaagtaatagttccagcaattcttgaatctacatttaacttcataagaaagaaaccccgaaaata

gtgtatttcaaatatctctctgcgaaaggcaaaaaaggaaatgtatgtacacctgattcgatttactaggctttaaaact

tccgttctatgtggatagtggttaatttaagcatttaatgtctcagcccagttgtattacttttgagacagaaacaagga

tggccttcaggataaactacttcagatgagtagtatacctatgatgatggacctcagtgtaagggcgggtgtcaccctac

tttctaacaaagaagtatagctttgataggcaacacacaagatttacagtacatcaagatggatgtaattttggaccttc

ccatAtccattgagacaaatttaacggaattataaagaaatctcagtccaccccctccaatcacagcaacacttaggttg

atatagtattacccgtttgtttggatttgaaattacacttaggttacctttcgaagaactctttgattttcgatgtatgc

tgcactgtgactttggccaagcatattatctcaaagaactctttggtttacggtgtatatatgcaacattgttaatttca

ggacggcaaaaaccagtatgaaggagagacgcagcaacagtggaaggagagacggcaaaaatgttaaacatgcatacaaa

aacatattgccgtatacgaaaaatatagacatcaaaaaatatttctaaaaaaacctaattgtgtattcaaaaaaaatgaa

aatgtataaaataatactggggaaaacgatacaAACAacacaCaGAACaAaaaTAaaCAaaAttTaaaaggatctaaaac

caaagaaaaaaAGAaaacccatacaaaacgatataaacaacacttagaacacaaaaaattaaattcataaagtcatagaa

aagtggtgaaaaagaaaattaaatgaagaagacggtgaaagtgaagaaaaatgaaggaaattaagaagaaaaagaaaaaa

cagagaagccaataaagaaaaaaaaaaagaaaaaagtgaaaactgcgggctctacaggcaagtggagctatcatctcgct

atgagcgagatatagctcccacgatggtaccgggtggcgaagcgtgcacaaacctaaccaagtgggcctgca

**bPb-7676**

bPb-2591

tgcagctgctaactaagcaatatactatatgcatacatatacagagaagaaaattgctttgacagaacaccgacaagcac

atgggaacagatatacaacaggaaatggtaaaaagtacactggaatttcttttgaattttatttgaatacataaGtttat

ttgaattttttctgtactccttaaaatcagaatcCggattcttagatcatgattcatctcatcaagatattacaatttgg

acaggacaatggaagaacttggagccaacgttttttttctgcgatttgcagagctatcactatattattttggaggcaaa

gatgtccctgtactaaggagagcaagcttgaatacataagaacaccttcaagctttatattgaatccctgatttgttgca

actaccaaattgctagttctgaattccaattagatagctgcatagtccagctagctacactagaagctagatcaagctaa

tttcgtggcacacatggctgatgcaaccaattgggactaataaacagaagagtaatcaaaccactctcctaaccaattga

aataccaatggttctttgcttcaggcaaccaccaaatagagcacttcatgtacagcttgtctggcagataacagggagag

cagtaggagatatagaggacggagggagatggatattgggaagtgggattaatttgatgcccaaacttttgtttagttta

ttttgggacttattagcgggaagtgaaggagaaaggtttggttgctcttttgatcagtgagtcagaAcaagaacgagatg

caacatttagaaattgaccattagtatttaatgcatctttatgcttttcttcttctatgcatgcgctcctgca

**bPb-1084**

bPb-2460

tgcagagtgaacaaacaaaaaacggcccactacagcaacaacaaataacgggctgcaaacaaaacagagaagtggttcaa

acccatgaatcgcggcgcaaaatgatcgaacggttgagatggtgactgagaccaaacctgaatctcagctagctgagttt

cagcaatcccgaatgatattacccgggactgatgctgatccaacggccagaaacacatggcatataagtcttcaaaaatg

tattgaatatatccgttcaaacatttgactcacatgtagtgcattgtacactgccccaagcatacatgaggtgtccaagt

ttggattctagaaccatattgcatgctgccaaatacgaccgtggaaaaacgcatgcaacctaactatattgtggctataa

atacctccaccgcgagagctgca

bPb-8072

TGCAGAGTGAACAAACAAAAAACGGCCCACTACAGCAACAACAAATAACGGGCTGCAAACAAAACAGAGAAGTGGTTCAA

ACCCATGAATCGCGGCGCAAAATGATCGAACGGTTGAGATGGTGACTGAGACCAAACCTGAATCTCAGCTAGCTGAGTTT

CAGCAATCCCGAATGATATTACCCGGGACTGATGCTGATCCAACGGCCAGAAACACATGGCATATAAGTCTTCAAAAATG

TATTGAATATATCCGTTCAAACATTTGACTCACATGTAGTGCATTGTACACTGCCCCAAGCATACATGAGGTGTCCAAGT

TTGGATTCTAGAACCATATTGCATGCTGCCAAATACGACCGTGGAAAAACGCATGCAACCTAACTATATTGTGGCTATAA

ATACCTCCACCGCGAGAGCTGCA

**bPb-8866**

bPb-33276

tgcaggagcgcatgcatagaagaagaaaagcataaagatgcattaaagactaatggtcaatttctaaatgttgcatctcg

ttcttGttctgactcactgatcaaaagagcaaccaaacctttctccttcactccccgctagtaagtcccaaaataaacta

aacaaaagtttgggcatcaaattaatCCcacttcccaatatccatctccctccgtcttctatatctcctactgctctccc

tgttatctgacagacaagctgtacatgaagtgctctatttggtggttgcctgaagcaaagaaccattggtatttcaattg

gataggagagtggtttgattactcttctgtttattagtcccaattggttgcatccgccatgtgtgccacgaaattagctt

gatctagcttctagtgtagctagctggactatgcagctatctaattggaattcagaactagcaatttggtagttgcaaca

aatcagggattcagtataaagcttgaaggtgttcttatgtattcaagcttgctctccttagtacagggacatctttgcct

ccaaaataatatagtgatagctctgcaaatcgcagaaaaaaaacgttggctccaagttcttccattgtcctgtccaaatt

gtaatatcttgatgagatgaatcatgatctaagaatccggattctgatttgaaggagtacagaaaaaaatcaaataaact

tatgtattcaaataaaattcaaaagaaattccagtgtactttttaccatttcctgttgtatatctgttcccatgtgtttg

ccggtgttctgtcaaagcaattttcttctctgtatatgtatgcatatagtatatagcttagttagcagctgca

**bPb-47406**

bPb-1807

tgcagcgtcatgatattcctttctgaagctcaaggttgttgtgatataattattcagcatgcaaggagttaaatgaagtt

agacaatgtcaaccatgtgttttggctctcctttacaacaatgtgtatcttatttttcagaaaaggaatagctttactac

actatagactctgcatcccaagcacataagtgtacatcatttgacttagtggaatgtaaatacgctcctttgtttagtga

gatatcaataagttaattttcgaactatggatccatttggaactaatgcaggaaggtgaacactatttaggacgtttttc

atgggtatacttgaaactgtcattcaacacccttagcacatttttaaagaggttgtgcctgtatatgtctagaccaaagg

aatacagtaaagagcatatacggataatatgaaagcttgttcttagatacggtgtgagttggtttgagttcggaatattg

gaagtgtagttcatttattttactttggtggttttagttttggaactaatgaagaaaggcgatgctacttacaattgatt

tcatgggcttactcaaaacctgtattgaacaacCttatcacgggaccggagcagagcaacattgacatcatcagtgcaag

tctccgccccttggctactcgcttatcgccaccccttctaccagcacgtctctggagcaatactagctcccttaccttcc

cagccctgca

bPb-0091

tgcagcgtcatgatattcctttctgaagctcaaggttgttgtgatataattattcagcatgcaaggagttaaatgaagtt

agataatgtcaaccatgtgttttggctctcctttacaacaatgtgtntcttatttttcagaaaaggaatatctttactac

actatagactctgcatcccaagcacataagtgtacatcatttgacttaatggaatgtaaatacgctcctttgtttagtga

gatatcaataagttaattttcgaactatggatccatttggaactaatgcaggaaggtgaacactattTAGgACGTttttc

atgggtatacttgaaactgtcattcaacaccctTagcacatttttaaagaggtTgtgcctgtatatgtctagaccaaagg

aatacagtaaagagcatatacggataatatgaaagcttgttcttagatacggtgtgagttggtttgagttcggaatattg

gaagtgtagttcatttattttactttggtggttttagttttggaactaatgaagaaaggcgatgctacttacaattgatt

tcatgggcttactcaaaacctgtattgaacaaccttatcacgggaccggaGcagaGcaacattgacatcatcagtgcaag

tctccgccccttggctactcgcttatcgccaCcCcttctaccagcacgtctctggagcaatactagctcccttaccttcc

cagccctgca

**bPb-0351**

**bpb-8592**

bPb-3436

catgcgttagctaaggtaggagtgtctactgatttatgtaaggtttggttgcatgcggatacgcagtctacgaagggcga

attctgcagatatccatcacactggcggcagctcgagcatgcatctagagggcccaattcgccctatagtgagtcgtatt

acaattcactggccgtcgttttacaacgtcgtgnngggaaaannnnggncagtgaattgtaatacgactcactatagggc

gaattgggccctctagatgcatgctcgagcggccgccagtgtgatggAtatctgcagaattcgcccttcgtagactgcgt

atccgcatgcaaccaaaccttacataaatcagtagacactcctaccttagctaacgcatg

bPb-1820

tgcaggacctcctcgagcctcctgatggccctgatcaagctgcctttgagcacttgggtcatctcctattgagagatacc

taacaatcccctccgcgaagaggcaaagatcagcaagcttcaaacccaagaatgcaaccacacgttagttgcttcagtac

atgaacaaagcaagtgaaagcagattcgaagcaactgaagggtgacaccttcagaatttagggaaggaggtactaccgct

atgatcaaccactagttcttccacacttactctatggatcaggaatcacaaccgtacataaaaattccccctgttatgga

atttgactgctcttgatttatctattgcttaaccatatagtagaactctgca

**bPb-7320**

bPb-3892

TGCAGCATCATCAAACCTAGGGAGGATCCCGACGGCGGGGTCGAGCATACGTAGTACTTGCTATTTCAGCTCAACACAGA

GGTTACCACCTGCGGCAGACTCAACGCTGTATATTTTCGGGATCGGACTGAAATGCACACAAGACTTGCGAGAAAACATC

AGACCCTACAGCGCCGGCATGGAGAGAATACTGTATAGCTAGTTACGAGCTTCTGATGCTTGAAGCTGGATACATAGAAC

TACGCACATGATAGCTCTAGGTACGATTCATACGGTCATATCTTGATTACCCTAGAGACATCGTGCATTTACGCTGGGCC

GGTGACAGTTTTCATCAAGGTTTAAATCGCGGACACTGCA

**bPb-5075**

bPb-0949

tgcagagttcatctatccagcaaactcagttggagaccagtcaatactatacaatcaaattaagaaggctgaacttcaga

ggtcgctagaccctccgccgttgacaagcgggggttctccgactcgacctcggaacgtcagtggacctgtggcagctttt

ggttacagtgcaacacaactagcactaacacacgcacaacacaatagtcatagttcacatgctataataatgagcaatat

tagttacagttaagttttactgcacactcagaataacaagttctgaactcggagcttcagattagccatcagttatagtt

ggtttgactttgagctcacacttggaatcacaatttataatttggcatttcagattagtcatctgaactaatgcacacaa

acagcggacaaatttcacaggccataactatgagcaacattagttgcagtgaggttttacttcacagttagaatcacaag

ttttaaacttggcacttcagattaggcatctgatataccagcatgcgtgcacaacagactaacaatttactagaaaacgg

tgtcaattcctaagaaagtactctacgaaacattgtcagctagacaagtagtagtatcactttgatcacagttcacaact

actcTctccgtacctaaatataagtctgtttagagatttcattagataactacatatagagtaaaatgagtaaacataca

atctaaaatatgtatacAtccatccgtatgtagtcacctagtgaaacctctagaaaaacttatatttagaaacagaggaa

gtactgtattagggggaggcagcacaatagttaacatcgcgggcctgca

**bPb-0786**

bPb-6824

ntgctggaattcgcccttgatgganccagtgcaggtgattcaatccatcgctaacgctttagcaactactccctacgtcc

aaaaataagtatctcaaacttaatatgatttatattaacattaatacaaagttaagatacttattttcagacgaaggaag

tataatatccgctcatacgtgattttctttacaagaaattgaaacatcactgattttgaaaaagttcataaacacgcaaa

aaagttcatgaaccctattagaaaacattgataaaaatagttcagaaaaatttgaaaaagatcatcaattttgtaaaaaa

ggtcatccattttgaaaaatagataacaaattcaagaaaattcatcatttttgaaaaacaaatcataaattttgtaaaaa

gttcatcacttttaaaaatagtcatcgaattgaaaaaaaatcatcaaattttggagaagatctccgaattgtttgaaaaa

gttcctgcaatttggaagaaagtccatgaacaatggaaaagaagattccattgattttgaaaaaaaatacgcgaattcaa

aaacaaatccgtgcatttagaagaaaacaaaaaaaaggaagagaagaaaaatcgaataaaaatgtcaccgaccagtgcac

tgggaaaacaaaacgtgaatagtagaaagaattaaaggaaaaacaagatcattgttgcaaaaccgtttgacatacacagt

ggttgcaagttcaaaacatggcggctgca

bPb-8924

TGCAGACTTCAGGAATCCAACACCTCCAAAACAACAGAGCAAGAAGCAAGAAAATAGGAACGGTAACACGAATCAGCGAT

TGTCAGCCCGTCATCAAATTAATTACATGGGACTTCTTTACGGTACCAATGGAAAATCAAGCATTTTACTCTGTTTTGAT

CTATCTGCTATTCTATTCTGAGAACAGCAAAGCATTTGTGACCTTCAAAATTCTTAAAAGGTAGCTATCGAAGCCCTTTC

TCATAGTCAAGCAAAGAACAAAAACCATTATCAAGTTTCACATCACGCCATGCAACTAAGAACAAAATCTGTATCACCAT

TCGCCACCACACTATGTCACTACCTCATCAGCAAACGCCTACCTCATCAACAAACGCCACGTTCTCCTCCTAAGTAAAAG

AAGTTACTTCTTCAGATCCGAGGGCTCAGCCTGTATTTTCATTATCAAGAACCACACCGTATACAAAGCTCATACGGACA

AGGAAACACATAAAGCACAAGAAGTAATCCATACAGACTCCTAACAGAATCTGATGTAAAGATCAACAAAGGACGGGAGC

AATATGTAGTCGGAACTGCA

bPb-5207

tgcagcaaatgcatgcatgctcctcgcttccgtagacgtcagcgtctccgcgttgtgtggccataactttaccgctgtga

ttcttgcgtgatgagggttgccaatgctgcatagttttatctgctggactgtttcacgctgttgtcccttcttcactgtt

gtggtcacgccatcatcaacgcaaatgttcggtagagtctatgtgctgataacgtgttaaatgtaaaaagtaaagatcga

ggagaggtgaatgaaaaatgattcattagtctttattgatgatagcaatggggtatttatacccgggcggaagtacacat

gttgcttgggagtcaagtaacgtgaagttacttaagggccaagttattacatagtttccttgagagtcaaggaagtgcat

gattgcttgaggactaagtaacctatctaagaattaacttaatcctaatcagcttaattaataagcaactaatctattct

taacagttttaaaagttgaggcttgtatattagacaattttgtagttcagggttgtgttttagacttcggtggtagttca

aagttggaatattctcttttactccatccgtaaacttttataagacgttctagatcactaatttcgttagtcagtttctt

ggcccttgccactttttggatcattctctgca

**bPb-4418**

**bPb-8637**

bPb-8803

tgcagcaacgagaacaatccatccatgagattcttgcaaacacatgtacctaaaacagttctggcctctaataatataaa

tatggtctttagattgatgtgagttggtttgtcctccgatagtagtaacatcaagcacggcggtcaatatatttgctgaa

atcagacaagtaatctggttcgttctacccatttgcttcgaatctattccctctgttcctaaatataagtttaaaaaaaa

ttacactacagactacatataaatgtatatagagttatagacatactttaaagtatagattcattcatattgctccatag

gtagtccgtagtggaatctctagaaagacttatattaggaacggagtgagtagtatgggccccttacacaaactgcaaga

aagctgtgagcaagtaccctttcaaagcccatcccaaccagcttaagctcttcatcaaatgtactgtcagtctgca

bPb-5271

TGCAGGATTTCAATTTTTGATAGATTTTCGGCCTTTTTCGTCCTAATTTCGTTTGTTTTTCGTTCAAAAATTCAAATTTT

AGGTTAGTCTTTTCGTCTGAATTTTTCCAAAAAAAATCCGAAATTTCTGAATTTTTCCCATTTTGTTTGGAGGCGATAAA

ATTTGCAAAACGAAATCCAAATTGCTGTCAGAGGCACCATAGATATGTGTGGCGGGCGATATACTTCACCTGCCAATGCT

AACTTTTGGAAAATTTGTGGCGTGCAAGGATTTGGCCAGCAAACTGCTGCTTGGAAAATTAGTAGCGACCAAGATGTCGC

CTGCCACTGCA

bPb-6805

tgcagagatggaaaaacaaaagaaggctgctgagaagagaaggattgcagaggaaaagaaaatgattacaaataaaaaat

accagcaggaggaggcagcttttctaatacatcaagtagaggagtttgagaggcacgagagttggagaaaattcaaggag

gatgaggaggctgcaaggcaaaatgcttgggagcaacagaacaatcaacaagacacgacagggaatgaagaatgggagct

gagaaagaggagagctgaagaaacaaacaagaaagcaatggatgacaatatggcaggtcaagcaaggaagcaagctgcgt

tggaggaacaaataaaacaaagagctgcagcaactagttcaagggccgaggaggcaagatttaatccatgccctattcaa

ggaaacaatacaaagaaaaaatctgcacctaccaatcagaacatgacatgttttaagaaacctagaaaggttaacatgtt

cgatgaatttaggtaaagggacgtgaagtttggaacaatcctcatctcttttttgtgtttctgcttatgatcaggcgcaa

aacttgtaatggcaatattttttttaggatcattgcgttgatcatgaagtggaagagctgtcttttattaGgaaacaact

attacagactttgtgttatttgtactgtgcaacttaattgaccagattgttcaagtgcaaatataatgcttatgtgtgct

taatttgtgcaaatctgatggccggcaacatgcgtggcagctacatgagtgcgacagctgatggagggatttgggtgctg

ca

bPb-2988

tgcaggatgagcaataccagtattacaagaacttggatgatgaaaagaagtatctcttggtgcttatgttgggtgatttc

caagatgccatggtaaccctattacctatctgtgttttttgaagatgaatgatgtttgtacgagtatctagtactttagt

tccgttaattgcagtcaatttcttgacgggcagactattcagttagtgtacgaaaaattccacatggttttgggggcaat

tacaagtaaggtccagagccattttccagtctaacatggtccctctcataatctcattaagaccatgttaaatttgctcg

tttcttaaagactaggaatgctaagagtcagatatcagaaaggcaaagtcgagatataccttgattataactgttgtcat

ttggttgcgcataaactatccgtagaagaagtcttatatctgtctgca

**bPb-8809**

bPb-9733

ngccagtgtgctggaattcgcccttgatggatncagtgcagctTgaTtttgtcCtcgtgcttgcctatagcatgtctgac

gaggtTggtgtgttctgtgcccagctgcaaagatggggaggataccgtccctgaagaacttcaatgcgttcccgcacgcc

gaagaccacctgctgaagaagacttactcgggtgctataggtacatagatacataatatgctgcctcgtgaactgttttc

tgctaatggatgcttatatcttggagatttgtgtctaatacgcgccatcgttggtatgttggcagtgacgattctcgggc

taattgttatggtaacacttttcgcgcatgagctcacgttttaccttacaacctatacgatgcatcaggtacgttttgtt

cttgttcaccttttcatgtagtttggctagcatatgtagcttattgtgatgtctatgttgttctgtccattcttacccaa

cttttatgactctagtttgtgttgggaatatgcctgtagatgattcaaatttctcctgtttaagtttattcactatgact

tacgaaaccaatctggccaaggaatctgcatgatgacaaaggaactatgaaagagagcagttagttacatttatataaaa

ttgtgtgccctttgatagtctcattgttgttaatatctgctagttgttgttttcatattagtgaaaaccaacttggacgt

tgtggttctatttcccttgttcggaatgccatctcaagtttttcttcaatatctgca

bPb-5529

TGCAGATTCGATCGGTCTCGACGAAGTCCACAAAAATAATCCATGGACATGCATACATTCGTAAAAGAGGAAAACTGCAT

CAGGAGATTGTTCAATTTGTCTTTGTTTTCTTGCAATCAATCTAAGATCTAATAACTACTTGTATTTTGGGAATCGCAAA

CAAGAAGTACAGGAGGAAGAACTGTGTATTTCCTTATCAAGAACAGATAAAAATCAAGAAGAACATGAATCAGGAAGCAT

AAATCGAATAAGAAAAAAGGGGAGTCAACTCACGAACTAGTAGCATACTTCACACACGAAGTACACAGATCAAAACTCTT

GAACCTTAATTGCTACAACATGAGTTGATCCTAATCCAATTTTTCTTGTTCTCACACACGAATACTAACCTGCTGGACAA

GGCCAACGGCGACGTAACGCTGTCCAGCGCAACCATGGCCATCGGCCTTCTCGGAGGCGGCCGGCGGCCATAGAAACCTC

CAGGCACTGCA

bPb-5379

tgcagtgtctggaggtttctatggccgccggccgcctccgcgaaggccgatggccatggttgcgctggacagcgttacgt

cgccgttggccttgtccagcaggttagtattcgtgtgtgagaacaagaaaaattggattaggatcaactcatgatgtagc

aattaaggttcaagagttttgatctgtgtacttcgtgtgtgaagtacggaatctgatttgatgctactagttcgtgactt

gactccccttttttcttattcgatttatgcttcctgattcatgttcttcttggtttttctctgttcttcataaggaaata

cacagttcttcctcctatacttcttgttttgcgattcccaaaatacaagtagttattagatcttagattgattgcaagaa

aacaaagacaaattgaacaatctcctgctacagttttcctcttttacgaatgtatgcatgtccatggattatttctgtgg

acttcgtcgagaccatcgaatctgca

bPb-8822

TGCAGAAAAGGTAGCACCAGAAAACGTGCTACACTGGTGGATTCTACTAATCACCATGGCAATATATTGAATGCAACAAA

TAAACTTAATTTGGACACGGTTCAGTTGTCTCAGAAACCACTATCCCATTCACCATGGTCAGAATGGTACTGTGTTTATA

TTTCAGGAGGCACGGCCAAGTTTGCCATTCGGTACAGGAGTAAGAAAAAGAAAAAAGAAACAGGCGTGTACTTGATTGTG

ACCAAACATAAATTACAGAAATGGATCCATGGAGATATCCGGACTTTTAAGTGCAGGAATGGGCAACTGCCACCAGCCCA

ACATAAGTTGACTCTCACCAGCATATAGACTTGGTACTTGTACAGTTAACCAGAGACTAGGTATGACTTTAACCGTTAAA

GGATAATGGCAACCCCACCATCCATTAGTATTACCATGGATTGCGGTTTACAGTTCTCCACTGAATAGTGACATAGCACA

AAGTAGATGCAACATCAAAGTACAAGAAGATTTACATACCTTCCTTCGGTTTCGGATTACGTGTTGAAGCATGCTCTACA

GCGGGTGATTTTCTCTCAACCATTCTGCTGTCCACCAATAGCTGCA

**bPb-1821**

bPb-4012

tgcaggcgggtccaaggaacgctggagcaaactgaaaaaaaacaaacagaatgatatatacagatgcctcgtatgttcca

tttccgaaccgtcgatttctacgacgccacaaaattgtacatataataccaatcataaattgtaaaactactgttacagg

tacgcgattatgctcgctgccatatatgtggaacacgataatgccgaagagcagggtgaagctattgtggggaagaaccc

acaatagaatgataggaactggaaataatagaatgatagcactagcagatgcctCgtatgttccatttccaaaacgttaa

tttCtatgacgccacaaaactgaacatataataccaatcataaattgtaaaaccaccgttacaggtacacgattatgctc

catgtcgtacatgtggaacacgataatgcccaagagcagggtgaagctattgtggggaagaacccgcaatttctattggc

ttacatgagcagtcagatcggcatcatgccatctgaacttgctgctcttttaggcaaaaaatacactgtcatggttactc

caagcagcaaatcgcttggcggcaatcacaattatttcccaaatgaaaaaggctaagcctttacgggagatgtcatcatc

agtctttggagctcagtacttcttcggtggctaaagaagatgcccagtgttctttctcaagaactttgtcagctggcata

ccatagagtagccttaacgcatcgccccacctgccagctcgagctcaaaataaataatattccaaatgtactagtactag

ctatgaaccttttagttttcattgttgccattttcatgcattcaagcagagggtcaaaaaGatttgaagaaGaaagaaag

caaaACaAccaagggcttctcgcatgggtgaataaagatgcagtctgctaagaatgacagacagtatttcttagtctaga

ccgtggacgttgtgggtcaaggtcaatgtagtttagatcatgtagtccatgtcatgattctgtgcgctgtatctaaagac

atggtcactgttactccttattagaatgactactgaattataatcgacacgtcagtgttatattgtgcaaactctttccc

attttttaatgttttcgcttctgca

**bPb-3316**

**bPb-5182**

bPb-2425

TGCAGAAAAGGTAGCACCAGAAAACGTGCTACACTGGTGGATTCTACTAATCACCATGGCAATATATTGAATGCAACAAA

TAAACTTAATTTGGACACGGTTCAGTTGTCTCAGAAACCACTATCCCATTCACCATGGTCAGAATGGTACTGTGTTTATA

TTTCAGGAGGCACGGCCAAGTTTGCCATTCGGTACAGGAGTAAGAAAAAGAAAAAAGAAACAGGCGTGTACTTGATTGTG

ACCAAACATAAATTACAGAAATGGATCCATGGAGATATCCGGACTTTTAAGTGCAGGAATGGGCAACTGCCACCAGCCCA

ACATAAGTTGACTCTCACCAGCATATAGACTTGGTACTTGTACAGTTAACCAGAGACTAGGTATGACTTTAACCGTTAAA

GGATAATGGCAACCCCACCATCCATTAGTATTACCATGGATTGCGGTTTATAGTTCTCCACTGAATAGTGACATAGCACA

AAGTAGATGCAACATCAAAGTACAAGAAGATTTACATACCTTCCTTCGGTTTCGGATTACGTGTTGAAGCATGCTCTACA

GCGGGTGATTTTCTCTCAACCATTCTGCTGTCCACCAATAGCTGCA

bPb-8101

tgcagaaaaggtagcaccagaaaacgtgctacactggtggattctactaatcaccatggcaatatattgaatgcaacaaa

taaacttaatttggacacggttcagttgtctcagaaaccactatcccattcaccatggtcagaatggtactgtgtttata

tttcaggaggcacggccaagtttgccattcggtacaggagtaagaaaaagaaaaaagaaacaggcgtgtacttgattgtg

accaaacataaattacagaaatggatccatggagatatccggacttttaagtgcaggaatgggcaactgccaccagccca

acataagttgactctcaccagcatatagacttggtacttgtacagttaaccAgagactaggtatgactttaaccgttaaa

ggataatggcaaccccaccatccattagtattaccatggattgcggtttatagttctccactgaatagtgacatagcaca

aagtagatgcaacatcaaagtacaagaagatttacataCCtTCCttcggtttcggattacgtgttgaagcatgctctaca

gcgggtgattttctctcaaccattctgctgtccaccaatagctgca

bPb-1241

tgcagaaacgcgcacgtgacgcataaagcccaccagagacatgccgttgtgcaaaagggcagcaacttgacgcacaacgc

gtggaactccctcgctgggtggcttcgccagcagttgtaattcttaaaaatgttaaacaactttctttcttcttttttcc

tttctgtttttattaaaagaaatgttttttattttcaaaaaatgttcttcaaattcttttttttgctttttaaataattg

ttctggattttcaaaaagtgttcttgattttcaaagaaaatatctggatttggaaaaatgtttgagaattggaaaaggtg

ttccagtatttgaagaaatgtccggattttcaaaaacatgtttcgggaatttgaacttttttttcggaattttgaataaa

tgtttcggaattagataaaatgttccgaaattttagaaaatgttttggaatttttatctaaatgtttctgatttttcaaa

gaaatgttccgaaattttacaaaatgttccataatttgaaaacatgtagaattttcgaaatcgtgaaaaccgttctggaa

tatagaaataagttttgagaacatgaacatttgtttcaatttgtgaataaatttcagtatcgctaacagttcgaagcatc

agacgaaactgca

bPb-8022

tgcagtttcgtctgatgcttcgaactgttagcgatactgaaatttattcacaaattgaaacaaatgttcatgttctcaaa

acttatttctatattccagaacggttttcacgatttcgaaaattctacatgttttcaaattatggaacattttgtaaaat

ttcggaacatttctttgaaaaatcagaaacatttagataaaaattccaaaacattttctaaaatttcggaacattttatc

taattccgaaacatttattcaaaattccgaaaaaaaaattcaaattcccgaaacatgtttttgaaaatccggacatttct

tcaaatactggaacaccttttccaattctcaaacatttttccaaatccagatattttttttgaaaatcaagaacactttt

tgaaaatccagaacaattatttaaaaagcaaaaaaaagaatttgaagaacattttttgaaaataaaaaacatttctttta

ataaaaacagaaaggaaaaaagaagaaagaaagttgtttaacatttttaagaattacaactgctggcgaagccacccagc

gagggagttccacgcgttgtgcgtcaagttgctgcccttttgcacaacggcatgtctctggtgggctttatgcgtcacgt

gcgcgtttctgca

**bPb-7395**

bPb-6579

TGCAGAAATTATTGACTGTCCTATATGGTTCTACACATTAACATAGTTATATTGACTGCACATGGCAACCCCACATGTTA

CTTCATGAACTAGAGGACTGGTAAATGTAGCCAACTTTGCTGATGTATAACCATCTGGGAGCTGTGATATTTTCTAGTGG

GGTGGTGCTGGATGCAGTGGTGCTCCTCACTTAGCATAGTAAAACAGTAATTATAATAGTGATAAGAGNNATGAACTAAT

GATAGTATCTGCTTAACTTTTCCCTTGA

bPb-0882

tgcagaaattattgactgtcctatatggttctacacattaacatagttatattgactgcacatggcaaccccacatgtta

cttcatgaactagaggactggtaaatgtagccaactttgctgatgtataaccatCtgggagctgtgatattttctagtgg

ggtggtgctggatgcagtggtgctcctcacttagcataataaaacagtaattataatagtgataagagtcatgaactaat

gatagtatctgcttaacttttcccttgaaacaaatgacttaagtttctcatgggagtgtttttttcttttgagaaaacgc

aagaagcctttgcgtttcattgtatagaaaagaggggggggggtacatcctcctaaAaggctgca

bPb-3879

agtgcagctagctacagacaatgtgatgaatgattttgggtgagtgatcagacattgcgccaactcaggcaacttgcaag

ctgctgtattttccagtgggcattgcagattcacaattaacatggggtcctcgtgccagaccggtcagtatggcaatctc

caagccaagctccatcacacaccaaacacttgctccagcacaattgtggctgctcatggtcttgtctcctgcacatggca

ggcctgagatataaaaccatggctaagtatctctaagcaccatcgctccatccacaggacaaagcattccaagctcgagc

tgacagaacaatctcttcagttttcagaaacacaagaagctaccatggtgagtgccacgagacttgctcagctggccaag

aaatggcagagaatggcggtcctcggaaggaagaggctcacttggtcatcggcggtgctcgaaaaagcaatcgaaggacc

gtgcagcgcgtcgtattcggtggtggctggcaagggccactgcgttgtgtatacggctgacggtgtacggttcgaggtgc

cgctggcgttcctaggcacggtggtctttggcgagctcctaaggTtgtgtcaagaggagtttggcttcgcaGgtggtgac

ggcaagatcacacttCCctgcgatgcagcgttgatggagtacgccatgtgcttgattaGgagaaGcgcctccgtggagat

ggaggtagcgttcctcggctccatggcaatgacattaacatgccactatgatgcaagttgtgtggtttcacgtgtgggag

ttgcccagcatgttgctctctgca

bPb-6126

tgcaggcatcatcttctcttgctttaccttatcgtttgctggtgtgggtGaaAGAggagcagtgctttttagctttgcat

ctttcctttGCttTGcccttgcgtggcttcccctgaatcatatgcattcatcgtgaggtataaggtatcagtggaggatc

gacacggattttgttgcgatcagcctatcaattcaacttttcatggcgcgagcagatgagttgacaagagatgaattatc

aacaaatcaagaacaaggaagcacagaagacaaaatatcaggatcaaggagtgattgttagaagtaatattgtctagcct

gagttagtcgtagtactagtggcaaattcctaaatatactactccccccgtttcaaaatataagtctttttaaagattca

actagatgattacatacggagcaaaatgaacgaatttacatcttaaaatgtgtctatatacattcgtgtgtagtattcta

gtgaaacttctatgtagtgcccttgcacgtcatgtgttctagtaatacaagttgtaatcaggtttggtgctacggtggcg

tgcaggtacaggttccggtcctgctgcatcatcagctcggccagcctgaagcctgaaatctgcaatgtttttttttcagg

tcatggttgcacgcaccatgatattcttccctgctaaaacaaatactctggatgattagcaaaacaaaagcacggcctgt

gttcgagtttttaaaaccaacgaactcttacatgatcggtgcctgactaataccaccctgctcatcggctgca

bPb-4583

tgcagggagatcaggagacaatgaggattatgctatgtatgaagaggaaacgcccctacatattgatatcagaaggttca

cgtacgcagagctgaagcacataacaaacaacttccattcaatagttggaaaaggaggttttggaactgtttatcatggc

acaatggaaaatggtgatgaagtagctgttaaggtgcttatggagacatcaatagcggagtcaacagatttcctccctga

ggtataccacaaaaccaactaggtcaaatcaataatttcttcaatagaaagtaaactataagtgtgcaatatatcaaccc

tttccagagtccatatccaattaaataatacaattgtcttcagctctaggatgatcggtaactatatctgttgctgca

**bPb-6135**

bPb-0710

tgcaggagcaaccatgcattcatatagacaccaccggggggaggggagtgttcaagtaagaacacaaatgtgagaaagac

tctgactacaaggaagcataagaatcaacccagaccctgaggagggtcagagatgcctccttaaatctaaaggattgaac

atggacttcttttaggacgagccattgtctaatagcggctattttgccaaaaaaatctatgtgttcctttgtttccagat

taagccaacaggccacgacaaagatgcctaagaatcaattccctgataaaccagcagtacagtggtgcaggatgtcttcc

ataccgtttctctgccggtctgcacaactgaatgctaagcttgttcgaatagttgacattaaaggggcctctaaaggaaa

gatagatgctggttttccatgccatttctacagcccgttgctgcaaaggacacaagggttaccattggagaaagtgaagc

gccgccgcgcaagaatttctttgtgtttacatgatcaaactaaatctatgtccagaaactcaatgctgca

bPb-3241

tgcagccgaatccaataggtcttaactggtgtgtaacttactgttccaaacgcctgccaaagcagtcaaagtagtattac

agcgtaGtatatttaaggaacctgcattttgcatagagcacacctaggcgggtttattcttgcaattaaggccctgttgt

ttctttaaaaagtcttaggacttttttagtcccaactaaaaagtctttagtccctacccgtttctttccaaggactaaac

atggactagaggtcactaaatgacatgcaaaaagatcatgttactcctagtaatatactagaagttattaaatgacatgc

taaaagtaggggcattgttgaaaaaaaagccaagaaagtctcaaaaagtccctccccatagagacttcttcttttagtcc

caaataccccttttagtccctaaaagtcccttctgtttctttcacatgggactaaaagggattttttttagtccgtacat

caaaaagtcactggaaagaaacaccccctaacagtactgatattagtgggctcgctacttgccctgtgagcttactgctg

gttacctgca

bPb-4115

tgcaggtaaccagcagtaagctcacagggcaagtagcgagcccactaatatcagtactgttagggggtgtttctttccag

tgactttttgatatacggattaaaaaaattccttttagtcccatgtgaaagaaacaggagggatttttagagactaaaag

gggtatttgggactaaaggaagaagtccctatggaagggactttttgagacttttttgcctttttttccaacaatgcccc

tacttttagcatgtcatttaataacttctagtatattactaggagtaacatgatctttttgcatgtcatttaatgacctc

tagtccatgtttagtccttgaaaagaaacgggtagggactagagactttttggttgggattaaaaaaaagtcctaagact

ttttaaagaaacagggccttaattgcaagaataaacccgcctaggtgtgctctatgcaaaatgcaggttccttaaatata

ctacgctgtaatactactttgactgctttggcaggcgtttggaacagtaagttacacaccagttaagacctattggattc

ggctgcactggatccgtcaagggcgaattccagcacactggcggccgttactagTggatccgagctcggtaccaagcttg

gcgtaatcatggtcatagctgtttcctgtgtgaaattgttatccgctcac

bPb-8771

TGCAGGATGCAGCAATACACTTCGATGTTTATATTTGTGTCTTGTTTTTCAATGTCATCTTCTGCATGCATTTGTACATG

TCAGCCTGTCAAACTGTAGTATGTCACTGTTAATCCTTACTTGATTTGCTTGCCTGTCACCATTATCAGCCTGTCTCACA

TTTCTGCTAAGCTACTTGGTTATTTGTTATCCTAACTAACATGACTCTTGGTTATTGGTTATCCCTTCTAATCTTGCTAT

AGATTATCCGTTGTGCTCTTCTGCCTGTTTGATCTGAGTATTATTGGAACAATATCTGCTACTCTGAATTTGTGCTGAAT

CATATTAAAAGTAGTTATATTTGACATTTGGTGTGCCGTGATTGATGAATTTGCGATGGTTATGTTATTTTTATCCGGCC

TGTTGTGCACCGTTTGNNCTACTTTACCTAGATCTGTTGTTTTTGGCCAGTGNTTGCCAAAGGCATAAAAGTAGTCACAA

TTGAATCAAATA

bPb-8731

tgcagagagagagagcatgttctgttctgcttctacaaatcctgaaccagattttctaagcacctccgaaaaagttaata

caatttcatgaaaaaaaagatcacaaattcacaaaaaaaatgtttacaagttcaaaaaacacttcatgaatttgaagaaa

gttcatacattttaaaagttgtcaaaatttgaaatagttcaagcattttcataaagaataaagaaagaacataaaaacac

agttcataaaaattaaatgaaaaaagaaatgaaagacctttttatttaaaataagtttacaaaatttctccacgaatctg

attttttttggttagtttcaaaatagtttttggaattgagaaaagttcaaataatataaaatacataaaaattcatgaat

ttgaaaaaaattaaaatgaacaaaatttcataaaaaaatagccaattttaaaagttcacaaattgagaaaatgttcatgc

atttgaaaaaagggagaacgaaaacgaaaatgataaaaacaagacaagaacgggccgaaaaaatgtgaaacaaaaggaaa

atgtttaaaacaactggatgatatctgaaggaaataagtcgtctttcccgcgtgacatgcgtcctcctagggccacgcac

cgcgtcgctgcttcCcttatcCttatccttagcgtcaacgcgagccgctcattccttctcttctatccagtgagatcgct

attcctcttCCAttttctagttctgccacacagctaaccatcttttgcatgctatcagccctcgtcggagcagctgtggg

cggcttgtaatgttccagcggccgacaatttgtagcaaacctgattcaacctgca

bPb-3700

tgcagagcttgcggacctctccctgtgtgccggttgccttgcacatgtgagactaaatatttccacatgttgccttgcac

aaatgggctgcgtgggcctctaggatttatttagggatttctgaaattatatattgggctggcacaaaactagataaaat

tccagcatgcaatgcgccttaaaccTtgtatattatgaaaatacacataaattttaaccgcgtcttCtaaaccgcgacgg

aggtagtatatatagtagagatatgtgactgaaatgtatatacgtgcacaacatcaatcctgttattgttgatctcgatc

atattccctccgttcctaaatataagtctttaaagaggtttcattaatggactacatacggatgtatatagacatacttt

agagtgtagattcactcattttactccgtatgtagacttctagtgaaatatcttaaaagacttatatttaaaaacggagg

gagtataacgtaacattttattagaaagtgatcaatgccacccaaatcaaaagaatggaaggtgcggtcagaagtaagta

catcaagatgagtccgaataaaaccaaaaaggagataaaatacaaccaatccgactggaaataaacacaaacgtctcgCC

Attatcggatgtttcctcagaaaatgacgacgatgatatcatcagttgaacttgctgca

**bPb-3910**

bPb-8462

tgcagagcttgcggacctctccctgtgtgccggttgccttgcacatgtgagactaaatatttccacatgttgccttgcac

aaatgggctgcgtgggcctctaggatttatttagggatttctgaaattatatattgggctggcacaaaactagataaaat

tccagcatgcaatgcgccttaaaccttgtatattatgaaaatacacataaattttaaccgcgtcttctaaaccgcgacgg

aggtagtatatatagtagagatatgtgactgaaatgtatatacgtgcacaacatcaatcctgttattgttgatctcgatc

atattccctccgttcctaaatataagtctttaaagaggtttcattaatggactacatacggatgtatatagacatacttt

agagtgtagattcactcattttactccgtatgtagacttctagtgaaatatcttaaaagacttatatttaaaaacggagg

gagtataacgtaacattttattagaaagtgatcaatgccacccaaatcaaaagaatggaaggtgcggtcagaagtaagta

catcaagatgagtccgaataaaaccaaaaaggagataaaatacaaccaatccgactggaaataaacacaaacgtctcgcc

attatcggatgtttcctcagaaaatgacgacgatgatatcatcagttgaacttgctgca

bPb-4058

TGCAGCGAATTAAATTATTCACTACAGAAGCAAGCTCGAATTGGGCTTGTAGTTGGTGGCGCACCTGCACATAGAAGACA

CTATGGGCATGGAGGACAGAGAGTTGTAGATGTGGAAGCAATTGTCGCCATGGCGTGGCGCAAGTGGTGAAGACCCGTT

**bPb-3887**

**bPb-1420**

bPb-4970

TGCAGATATTGAAGAAAAACTTGAGATGGCATTCCGAACAAGGGAAATAGAACCACAACGTCCAAGTTGGTTTTCACTAA

TATGAAAACAACAACTAGCAGATATTAACAACAATGAGACTATCAAAGGGCACACAATTTTATATAAATGTAACTAACTG

CTCTCTTTCATAGTTCCTTTGTCATCATGCAGATTCCTTGGCCAGATTGGTTTCGTAAGTCATAGTGAATAAACTTAAAC

AGGAGAGATTTGAATCATCTACAGGCATACTCCCAACACAAACTAGAGTCATAAAAGTTGGGTAAGAATGGACAGAACAA

CATAGACATCACAATAAGCTACATATGCTAGCCAAACTACATGAAAAGGTGAACAAGAACAAAACGTACCTGATGCATCG

TATAGGTTGTAAGGTAAAACGTGAGCTCATGCGCGAAAAGTGTTACCATAACAATTAGCCCGAGAATCGTCACTGCCAAC

ATACCAACGATGGCGCGTATTAGACACAAATCTCCAAGATATAAGCATCCATTAGCAGAAAACAGTTCACGAGGCAGCAT

ATTATGTATCTATGTACCTATAGCACCCGAGTAAGTCTTCTTCAGCAGGTGGTCTTCGGCGTGCGGGAACGCATTGAAGT

TCTTCAGGGAC

bPb-4318

TGCAGATATTGAAGAAAAACTTGAGATGGCATTCCGAACAAGGGAAATAGAACCACAACGTCCAAGTTGGTTTTCACTAA

TATGAAAACAACAACTAGCAGATATTAACAACAATGAGACTATCAAAGGGCACACAATTTTATATAAATGTAACTAACTG

CTCTCTTTCATAGTTCCTTTGTCATCATGCAGATTCCTTGGCCAGATTGGTTTCGTAAGTCATAGTGAATAAACTTAAAC

AGGAGAGATTTGAATCATCTACAGGCATACTCCCAACACAAACTAGAGTCATAAAAGTTGGGTAAGAATGGACAGAACAA

CATAGACATCACAATAAGCTACATATGCTAGCCAAACTACATGAAAAGGTGAACAAGAACAAAACGTACCTGATGCATCG

TATAGGTTGTAAGGTAAAACGTGAGCTCATGCGCGAAAAGTGTTACCATAACAATTAGCCCGAGAATCGTCACTGCCAAC

ATACCAACGATGGCGCGTATTAGACACA

bPb-6179

tgcagcagcaccgttttagttaatgttcggatccacataaagattcacaattaaattataaaagaacataaataggacac

tttgataggttatatctcaaagtacatatgcaatttatatgcaaaagaaatatgtaaactaaacttgtacaacacaaata

ttatatgcaccttcggttcatttttgaaaataccagataattggagatttgatgtgtatagttagaagaaccaacatgtc

tctacatagaaaataagtgttggccatgcaaaaaacatccgtaatcatgtttgctagaaataatagtggactatattggc

gtcgtttgattggacttgaaagtttgctcaataaaaatcctatgaaattcctttgaattaaaggagcccttatttcctat

aagtttcctattcctataataattaatattaccaaaggagaccttatatttaggaacggagggagtaaaacccttcgtca

catttatcgaaggttcttttgcggtgagttttctttgtcattgtgatagtcgacgtcagtgtttcactgca

bPb-4595

tgcagcagcaccgttttagttaatgttcggatccacataaagattcacaattaaattataaaagaacataaataggacac

tttgataggttatatctcaaagtacatatgcaatttatatgcaaaagaaatatgtaaactaaacttgtacaacacaaata

ttatatgcaccttcggttcatttttgaaaataccagataattggagatttgatgtgtatagttagaagaaccaacatgtc

tctacataAaaaataagtgttggccatgcaaaaaacatccgtaatcatgtttgctagaaataatagtggactatattggc

gtcgtttgattggacttgaaagtttgctcaataaaaatcctatgaaattcctttgaattaaaggagcccttatttcctat

aagtttcctattcctataataattaatattaccaaaggagaccttatatttaggaacggagggagtaaaacccttcgtca

catttatcgaaggttcttttgcggtgagttttctttgtcattgtgatagtcgacgtcagtgtttcactgca

bPb-0835

tgcagtgaaacactgacgtcgactatcacaatgacaaagaaaactcaccgcaaaagaaccttcgataaatgtgacgaagg

gttttactccctccgttcctaaatataaggtctcctttggtaatattaattattataggaataggaaacttataggaaat

aagggctcctttaattcaaaggaatttcataggatttttattgagcaaactttcaagtccaatcaaacgacgccaatata

gtccactattatttctagcaaacatgattacggatgttttttgcatggccaacacttattttctatgtagagacatgttg

gttcttctaactatacacatcaaatctccaattatctggtattttcaaaaatgaaccgaaggtgcatataatatttgtgt

tgtacaagtttagtttacatatttcttttgcatataaattgcatatgtactttgagatataacctatcaaagtgtcctat

ttatgttcttttataatttaattgtgaatctttatgtggatccgaacattaactaaaacggtgctgctgca

bPb-1719

tgcagtagttaacctatagaaagagtattgttaatagtttcttattttgattggcttgccaaagccaagccagacagggg

acgacaaagagtctcatgttgtttgccaatagagacccatcatgatgatttttaggcagggggaatggtagtttccatgt

gttcctctattccctattgttgacaagttatttcttcggtacttctaacgctgacgttggtattcctttttcggtggttt

atcttacggatctacagacacggaaatcgtacgtgtttatgcgtactggtgctggtcgggtgctcgttgaatcagatgct

gggagggggcaggtcttggagctgttccggtagcaaactccacaggaaacttcggctgcatctgggctgca

**bpb-0799**

bPb-5766

tgcagtgagctggatgatgagagctgcattacacgctcctcatagtcatgcttgacctctccaagtatcttcaacacaga

aacagctgggaaacagtttaatacttttgaagatgatgatttgctgctgctgctgctaagatgagatacatctaattcga

gactcttaactgaacggaacacactactgccatccagcccttcagggaacaaaccgtcgcatcttccgacagtcatatct

tCtactttatccagattgGgggaggccaaaccaccgtcataccctcctgtaacagccaatttctttccatcataataaaa

aaacGTtcccaaatcaggtcgcggaaagccttttctccttgaattatctcttttaacaacaAaatatgtcagtttggagg

tgtgaggcatggaaggcaGagacatctTggGgcacccaGaaacttcaagtctgcgcacattggtgcaacatacctccgac

aagggcatcacacggagattgggacaacggacgcattcgattatttcaagacttggaaaggaacaacaattaggttccac

aacccattcagcaagttctggcatctcataaaactcaacttccttcaagcgcataaaacatttgcctgaagcgccaccac

agacaggcccaaactgaagcattccagaaattttcctcaatttgagtcccttgagatttggtagcttcgcaaaaggtggg

agggtgctccnagatacgccttgtagagtgagagtctctanacttgtt

bPb-7360

tgcaggttaacaagtttagagactctcactctacaAggcgtatcttggagcaccCtcccaccttttgcgaagctaccaaa

tctcaagggactcaaattgaggaaaatttctggaatgcttcagtttgggcctgtctgtggtggcgcttcaggcaaatgtt

ttatgcgcttgaaggaagttgagttttatgagatgccagaacttgctgaatgggttgtggaacctaattgttgttccttt

ccaagtcttgaaataatcgaatgcgtccgttgtcccaatctccgtgtgatgcccttgtcggaggtatgttgcaccaatgt

gcgcagacttgaagtttctgggtgccccaagatgtctctgccttccatgcctcacacctccaaactgacatattttgttg

ttaaaagagataattcaaggagaaaaggctttccgcgacctgatttgggaacgtttttttattatgatggaaagaaattg

gctgttacaggagggtatgacggtggtttgGcctcccacaatctggataaagtaGaagatatgactgtcggaagatgcga

cggtttgttCCctgaagggctggatggcagtagtgtgttccgttcaGttaagagtctcgaattaGatgtatctcatctta

gcagcagcagcagcaaatcatcatcttcaaaagtattaaactgtttcccagctgtttctgtgttgaagatacttggagag

gtcaagcatgactatgaggagcgtgtaatgcagctctcatcatccagctcactgca

**bpb-4725**

bPb-9244

tgcaggttaacaagtttagagactctcactctacaaggcgtatcttggagcaccctcccaccttttgcgaagctaccaaa

tctcaagggactcaaattgaggaaaatttctggaAtgcttCAGttTgggcctgtctgTggtggcgcttcaggcaaatgtt

ttatgcgcttgaaggaagttgagttttatgagatgccagaacttgctgaatgggttgtggaacCtaattgttgttccttt

CcaagtcttgaaataatcgaatgcgtccgtTgtcccaatctccgtgtgatgcccttgtcggaggtatgttgcaccaatgt

gcgcagacttgaagtttctgggtgccccaagatgtctctgccttccatgcctcacacctccaaactgacatattttgttg

ttaaaagaGaTaattcaAggaGaaaaggctttccgcgacctgatttgggaacgtttttttattatgatggaaagaaattg

gctgttacaggagggtatgacggTGgtttggcctcccacaAtctggataaagtaGaaGatatgactgtcggaagatgcga

cggtttgttccctgaagggctggatggcagtagtgtgttccgttcagttaagagtctcgaattagatgtatctcatctta

gcagcagcagcagcaaatcatcatcttcaaaagtattaaactgtttcccagctgtttctgtgttgaagatacttggagag

gtcaagcatgactatgaggagcgtgtaatgcagctctcatcatccagctcactgca

bPb-7292

TGCAGGCATGGATGAAACAATCATAAGAAGATGCTGTGAGCGAAGCTAGCTGGTGGTATACCACTACCACGTGCACTAAA

CTCATTGCACGTGATGTGAGTTAAGCTCACAAGTTATTCAGCATTCTGAATTAGCCAATTGATCATGGATTTGTGGTCAA

TATACATCTTCACAATCTCTTCTGCTGCCTCTCTAGCTAGTGAGGGAGTGAGTGACACTGACCAGACTAGCTAGCTAGGT

AGTGACATCCTTTGCAGTAGCCAATTGATCAAGGAGGGGGACAAGTCACAGAATTAATCTACAATTCGCCCACAGGCTGC

TCCGTCATCAAGTCAATGTAGCCGCTTGGTTTTCACTCAGTTACATTGGGATTACCTTACCATTCTGCA

**-------------------------------------------------------------------------**

**Chr. 6**

bPb-5926

tgcaggagtatgtagctcctccaaaaacttatttgtacaaatgatataattagcacattttcataaccttttcattcatt

actgtatatttccgtaccgtgtaaaccgaaacatttttcaaacaggatagaagtcatattacaacaaccttttcttcatt

tcgtgcttcgggattagtctttgcagcagtatgtggctccttttctatagaaAtcatacatctttagcaccattttCgta

ccctctttattcatcactctatattttcacttcatgtcaaaatgaaaatatatttgtatagcatctccacatattcaact

cacatttccatcatttcaatcttcaagagcagttgccgttgcattatatacctcacacaacgacttacatgtggaaatcg

gacaacttgtaatattattcgtattgTttcggtccaacatggtatattTtctcttatctgaaaataaaaattttcaaata

gacgaatcgacatatgctagtaatcccgcttcatttcttgctttcgaatatgtctctacagcaagatgtagctcctccaa

caacttatatatggaaatgatatcatttaacacttttctcgtaaccgttttattcattacatatagattttctcttcccg

gggaagtggaatatttttcaaatagggtagtcgccatattacaacatcattttcttcatttcatagtttgccactacttt

gtgcagcagtatgtggctccgccttatgtaggaattatatatcattagcatcatttttgcatcatcttagtcagtcactt

tagatttctgttcgtctaaaactgaaaattatattatatggcattgtcgatagataagagtagctttcgaccaattctat

attacatcatagtttttgttgcatgatgtacctcaaacaacaacttatatgtagaaatcggacaactcgtaatatgtttt

gtaccatttcgatccatcatggtatattttctcttttgtgaaactgaaaagttgcttaaataggaaaatttacatatgat

atcagtcctgcttcatttgttgatgtcaaagttgtctctgca

bPb-4754

catggcaagataaaaaaagttgttatgattgtgagccgtgataatttgagtccaccaaatcaatgtgtcaggtctacatg

cggatacgcagtctacgaaagggcgaattctgcagatatccatcacactggcgGccgctcgagcatgcatctagagggcc

caattcgccctatagtgagtcgtattacaattcactggccgtcgttttacaacgtcgtgantgggaaaacannggcnagt

gnnttgtaatannanntcactatagggcgaattgggccctctagatgcatgctcgagc

**bpb-2380**

bPb-8937

tgcagtagatacacacatgaccttttatttttttatagaaaattcatgacctaattaacatggcaagataaaaaaagttg

ttatgattgtgagccgtgataatttgagtccaccaaatcaatgtgtcaggtctacatgcagtttggcaatagagtaacat

aaatcatcacgtaaaaaaaattctttcattgcaaaaaaagaaatctggagtctccaaatagatctacaccataataactc

aggcccaccggataaacggctctcaacgtgtgaatttcttgggagtctctaattagtattagtatagaagtatagattaa

caAcctctgcaacaatttattaagcaaacgtgagtacaaaagtactcagcaagactttgtgataaactatctactcatgc

aatgtatcaataaggaattgtggggtttcatgcggaaagccaacatttgactcatggctagacaagttgcattttcaaat

agtttagacaactttgatttctcgcacacgagtccactaacaccacaacaatactccatcatggaaccattccgtctcca

tacggaaatgccgtccacgacactcacgcttatcttgacaattttatgagtACcaactatagttatctatgaacaacata

tgtttccaagtagtccatatccgcggacgtagctattcgaatagatcataacCCtgca

bPb-5433

tgcagggttatgatctattcgaatagctacgtccgcggatatggactacttggaaacatatgttgttcatagataactat

aattggtactcataaaattgtcaagataagcgtgagtgtcgtggacggcatttccgtatggagacggaatgattccatga

tggagtattgttgtggtgttagtggactcgtgtgcgagaaatcaaagttgtctaaactatttgaaaatgcaacttgtcta

gccatgagtcaaatgttggctttccgcatgaaaccccacaattccttattgatacattgcatgagtagatagtttatcac

aaagtcttgctgagtacttttgtactcacgtttgcttaataaattgttgcagaggttgttaatctatacttctatactaa

tactaattagagactcccaagaaattcacacgttgagagCCGtttatccggtgggcctgagttattatggtgtagatcta

tttggagactccAgatttcttttttttgcaatgaaagaattttttttacgtgatgatttatgttactctattgCcaaact

gcatgtagacctgacacattgatttggtggactcaaattatcacggctcacaatcataacaACtttttttatcttgccat

gttaattaggtcatgaattttctataaaaaaataaaaggtcatgtgtgtatctactgca

**bpb-6554**

bPb-9447

TCGAGCGGCCGCCAGTGTGATGGATATCTGCAGAATTCGCCCTTGAGTAGTGCCAGAACGGTCCATGGGTTGTGCGCTTA

CAAAAAGTTCATTGATGCCTACTGTTGCCTTGCGAGAAATTTGTGTTATGCGACAGTCGATCCGGATACGCAGTCTACGA

AAGGGCGAATTCCAGCACACTGGCGGCAGTTACTAGTGGATCCGAGCTCGGTACCAAGCTTGGCGTAATCATGGTCATAG

CTGTTTCCTGTGTGAAATTGTTATCCGCTCAANNTAAACGAATGNNTTGTAGCTTGATTGNTTCCTCCACGCTNNTGCAC

AAGATANCCATCNCNCGATCGTCACTGATCCTCNNNGCTGCA

bPb-7829

TGCAGGGAGCATGATTTGTTCTTACTCTGTCCCACAAAGCGAGTTGATGGCCTGAGAATATTAACTATTCTTAGCTTCGT

TCAACTTTTCACCTTTTCTTCCAGTAGCACTTTTTACTTTAGTTAAATTTAAACATGTGTCTAATTTCAACAGAGAATTT

ATTTTACTTTAAATTTTTCTGTTGGTCAATACTGTCTCGGGCTATTTATGTGACCATAAAATGCACAAGAAATAAATATG

ATTCTTCACATACTAGAAAACACTTTGTGCTGTGTTATAGATCAGTTATCTAAAATAAACTCACCTGACCATGATCTTGG

GTTCTTGGGCAAGCAGCCAGTGACTTCTGCCGTTCCAGATGTTAATATCTAGATACAGTATCTCAAAACTTCAGTTCTGC

TGGTCGGTCATTCTTACCCAGTTATTTATCACATTAGCTACAATGATGGCCCACATTTCCATCTTTGAGAAAACTTCTCT

TAAAGATGGGGATTTGCTGCCTTTGACTTTTGATTTACTTCGTCCAGCTAGAATTAGTTGCAGCTCAGATTTGCTTATAG

TACGGCTGTCGATAGGGTCACTTTCTACATTCAACAGCCATACAGAGAGCCACAAGTAACCAAGTGATGCGAAGAAGGCA

AATGTTCCGGCGAGGCCTATATGTGACATGATGATCGGTGTTGCCAGGAAGCTTACGACGTTTCCAAGATGGAATCCTCC

CATGGAAAGGCCAACAGCAGTGGCACGTTCATGTGTTGGGAACCACCTGTTTCATGTTCNAAAATGAAATTAANAGCCTG

AACTGAAGTTAAGTTTGATTGCATCAATATATGTAGTGAATGNTTCATTTTCTGTAAGTAATTGGNNNNNNNNNNNNNNN

NNNNNNNNNNNNNNNNNNNNNNNNNNNNNNNNNNNCAAGGACCAGAGTGCAGCAGCACNNNCCATCACCTTCTTCCCCCC

GTATCGGTCCNCCAAAGCTCGTACAACCATGGATGAGAAAACATACCCCCATAGAAATGATGACTGCA

bPb-6071

tgcagctcaagcctctaatgcgcaagtagttcggcaagatggccaagcctgtcaagaactaggtactaacgccacaaaaa

ttctacttgaaaagttggaatgcacatgttatgcttggtttctatgctgcaactcatttgtttgtgcacctgctcttata

gtcattaaacttttttatggattgtactgttaacatttaagaagttttcctgtcattggagattttgttttggtctttgt

ttttaaatggtgaccaaaggtatattgatgtaattagccgtaacaacaaggatatgaagtggtcaattggttattcagtg

aaatgttttttacgcttggcatctatattcaaccggttaagaaaattgttttttataaaattagattcacgttagtgagc

gtatattaaattatttttatccgttgcaacgcacgggtatttttgctagtacacctaaaaacgccaaataacatgggcac

ggcggcaaaccctatctagcgccaccggcaccgttgaaacataaaatgtgctaaccggcgcctgca

bPb-9873

CACTAGTAACGGCCGCCAGTGTGCTGGAATTCGCCCTTCGTAGACTGCGTATCCGGATCTATAACACAGCACAAAGTGTT

TTCTAGTATGTGAAGAATCATATTTATTTCTTGTGCATTTTATGGTCACATAAATAGCCCGAGACAGTATTGACCAACAG

AAAAATTTAAAGTAAAATAAATTCTCTGTTGAAATTAGACACATGTTTAAATTTAACTAAAGTAAAAAGTGCTACTGGAA

GAAAAGGTGAAAAGTTGAACGAAGCTAAGAATAGTTAATATTCTCAGGCCATCAACTCGCTTTGTGGGACAGAGTAAGAA

CAAATCATGCTCCCTGCACTGGATC

bPb-1009

tgcagGgagcatgatttgttcttactctgtcccacaaagcgagttgaTGGcctgagaatattaactattcttagcttcgt

tcaacttttcaccttttcttccagtagcactttttactttagttaaatttaaacatgtgtctaatttcaacagagaattt

attttactttaaatttttctgttggtcaatactgtctcgggctatttatgtgaccataaaatgcacaagaaataaatatg

attcttcacatactagaaaacactttgtgctgtgttatatatcagttatctaaaataaactcacctgaccatgatcttgg

gttcttgggcaagcagccagtgacttctgccgttccagatattcaaattgcagtataattgtcctacaggtgcatttctc

caataagtttttcccagcagttgctgca

bPb-8075

tgcagaagaagggagacatggccaaggacaagttctccaaggtcctttggggcgccgttaccaactaccggacatggatc

tttgtcctcctctatggctactgcatgggtgtcgagctcaccactgacaatgtcattgccgagtactacttcgatcactt

ccacctagacctccgtgccgccggtaccatcgccgcttgcttcggcatggccaacatcgtcgcacgtcctatcttgtgca

cttcccgtcgctcttgtcatgggcatcatgatcatggcttgtacacttcccgtcgctcttgtgcacttcccacagtgggg

ctccatgttcttcccagccagtgtcgatgccacggaggaagagtactacgcctccgaatggtcggaggaggagaaaagca

agggcctccatatcgcaggtcaaaagttcgccgagaactcccgctcggaacgcggtaggcgcaacgtcattcttgccaca

tctgccacaccacccaacaacacgccccagcacgtataagatccaaattttcttctaccaaccaaatgtaaagctcgcat

aaatatacacatcatatatgtctactcctgccaaccccagttgttctgtacttgtagatgttaggaaaccttcaacttca

tctgtctgca

bPb-2751

tgcagctcagagctcaccggcataatagctcaaccaatatttgcgacaccgtacccacgaaggaatctcacacacatgca

tgcctagactcgccattacgtctactttctttctttatataaacataaaaaagaggttttcccactcgattaattatgta

atactccagtgttggagtacattgtagcaccacaaaatagtagttaagctgcaaatccgatggtacaacctcgtcggaac

acacgatatattaaccggcaacagctgcggctgcaatgccgctgacgatatcgatcttagctgcaacattaaagagcaca

gtagctagtactccatcgaagtgcagcaagcaggtaggtaggcggtcgatcgaatcgacatggacctaattaagggttgt

agtagacgatcttgatctcggtgacgccgtggcagttgccgccggggacggagtgctcgaagcagtggccgccggtgtcg

cagccggcggtggggcagtggccctgca

bPb-2628

CATGCTGGTAAAGTCCCAGAGAACTGATTATTGCTGAGATCAAGAAAGATGAGTCGTGTACATTTCTGCACTGGATCCGG

ATACGCAGTCTACGAAAGGGCGAATTCCAGCACACTGGCGGCCGTTACTAGTGGATCCGAGCTCGGTACCAAGCTTGGCG

TAATCATGGTCATAGCTGTTTCCTGTGTGAAATTG

bPb-4315

tgcagcgacgtagatgtaAgacaccCagctTgCtgctgtgtgaatgttggaATATGCaATCTgggattgtcgccaatTgg

taatagtagttataatctggtaagatgaagcaggatgttgcctcagaaagcaccatgccaatatcatcaggtagcaggcc

cttcttgtgctccaccaaagccaagtgatcatgaagcctgcactgcatgtggcgacgaatgacatcatcattacagaaat

aaagccatttcgtagagatgtttctctgtacagcatcggctactccccatgccgacgtatcgccttgcccttgtataatc

ccatcgcatacaaagtagttagaaacccagttctcccaattgaacatactgcaatgtttgagtgcatcggcatactggat

gcacttctcaactattatatggttcatgtaaggttctggaatgccgatataagcggcaatctccttggactcttccaata

ttgcttcatcgcggatgcaatcatcatgaaaataattgaccataacatcagtgtgcctctccatgtaatcaaaaccgggt

ccaaacctcccgtgcgaggtccataataccttaatactcaaaaatgatattactgggactccacactcgtacagatcaat

gtattcatcactcccattgtgaaatatcaccacaaatctactgctagcaagctttatgaagatttctgttctgatatctt

ctagcacccctcgagagccttcgtctatcccactgaaatcatcctcctcgtcctgctgatcgaagatggccatcaccgac

ggcggaagctccagctcctccgcgactgccttctgca

bPb-7036

tgcagcgacgtagatgtaagacacccagcttgctgctgtgtgaatgttggaatatgcaatctgggattgtcgccaattgg

taatagtagttataatctggtaagatgaagcaggatgttgcctcagaaagcaccatgccaatatcatcaggtagcaggcc

cttcttgtgctccaccaaagccaagtgatcatgaagcctgcactgcatgtggcgacgaatgacatcatcattacagaaat

aaagccatttcgtagagatgtttctctgtacagcatcggctactccccatgccgacgtatcgccttgcccttgtataatc

ccatcgcatacaaagtagttagaaacccagttctcccaattgaacatactgcaatgtttgagtgcatcggcatactggat

gcacttctcaactattatatggttcatgtaaggttctggaatgccgatataagcggcaatctccttggactcttccaata

ttgcttcatcgcggatgcaatcatcatgaaaataattgaccataacatcagtgtgcctctccatgtaatcaaaaccgggt

ccaaacctcccgtgcgaggtccataataccttaatactcaaaaatgatattactgggactccacactcgtacagatcaat

gtattcatcactcccattgtgaaatatcaccacaaatctactgctagcaagctttatgaaGatttctgttctgatatctt

ctagcacccctcgagagccttcgtctatcccactgaaatcgtcctcctcgtcctgctgatcgaagatggccatcaccgac

ggcggaagctccagctcctccgcgactgccttctgca

bPb-2768

tgcagcatactcaacaaaatgtattacttattcagtcttcacaggccggctttataccaaaagataccactacttgtagc

atctgtatgtgcatggtgtggcaatgtcttttgctcaatgtcttcggtcacatctatggtgttgcaatctttttgctaaa

caacatgcctatgcttgtgctttcccagctgtcgtgttgctacaaatttttgaaggacgactacnagaaactacactcaa

ttgatgcaaccgctctttatagaattgcgctatcATCggatcccgatgcggttgttagcgtcagtggcagtgattccgtg

tgttcagacggcccaGcaaAcaAcagccatagccAaatgacgtcaagcgaaaacgaaagcatatgcaggaTcggtaggtc

ttactcctagtgctgcttaggagtaaaaagTtcagtacacatgtttcattttcccatagcacacatacaattactatcat

gtgtttttaacctgctgctgcccagtagtccaatgcaaattgctggtcttagccttcacaactcaaagtcaactaagtaa

ccgctcaaacagttaaggtgctgcttctttttgggaagcattgaagcacaactcttaacagaaatgttagcaacggtctg

ca

bPb-0742

tgcagccactgtgtaatgtgattacaggcccacatgttggcagttggcacctcccattagtggaagaacttgaaaaataa

ctatggcccttgattactcaagcaagaataaataaacgtggcttttgatttgctacattgatgtgtttcgagtaactgtg

tacggctgaagcctcgcgccgttgttgttcttcactgccccaatatgtctgcaccgcagtggcggacctagaatctcagc

tttggggatgccacaattcataattaacactaattatcattgtagtatcaataaatagtcttaaaatagcatcactttga

taactatttagtgttaaatacaaaaagaaatatagtgtacaattttatagggggggccgcccactggatccgcccctgct

gcaccgtatatgtaaacaagcgatactgggatcttgaatatgtcaccaatgttccaagactcaaaagcagtatatgtagc

tagcgcggtaaagtagctgagagcaaaatgtcctgctccaagctatagcatctgtcgctgcatttagatgatgaattagc

tctgttattgaggtcagcaggcaacgagtcagaaattgatctagttcctgca

bPb-3586

TGCAGACAGATGAAGTTGAAGGTTTCCTAACATCTACAAGTACAGAACAACTGGGGTTGGCAGGAGTAGACATATATGAT

GTGTATATTTATGCGAGCTTTACATTTGGTCGGTANAAGAAAATTTGGATCTTATACGTGCTGGGGCGTGTTGTTGGGTG

GTGTGGCAGATGTGGCAAGAATGACGTTGCGCCTACCGCGTTCCGAGCGGGAGTTCTCGGCGAACTTTTGACCTGCGATA

TGGAGGCCCTTGCTTTTCTCCTCCTCCGANNNNNNNNNNNNNNNNNNNNNNNNNNNNNNNNNNNNNNNNNNNNNNNNNNG

AGGTCTAGGTGGAAGTGATCGAAGTAGTNCTCGGCAATGACATTGTCAGNGGTGAGCTCGACACCCATGCAGTAGCCATA

GAGGAGGACAAAGATCCATGTCCGGTAGTTGGTAACGACGCCCCAAAGGACCTTGGAGAACTTGTCCTTGGCCATGTCTC

CCTTCTTCTGCA

bPb-4246

tgcagaaggcagtcgcggannagcnggagctnnccgccgtcggtgatggccatcttcgatcagcaggacgaggaggatga

tctcagtgggatagacgaaggctctcgaggggtgctagaanatatcagnacagaaatcttcataaagcttgctagcagta

gatttgtggtgatatttcacaatgggagtgatgaatacattgatctgtacgagtgtggagtcccagtaatatcatttttg

agtattaaggtattatggacctcgcacgggaggtttggacccggttttgattacatggagaggcacactgatgttatggt

caattattttcatgatgattgcatccgcgatgaagcaatattggaagagtccaaggagattgccgcttatatcggcattc

cagaaccttacatgaAccatataatagttgagaagtgcatccAgtatgccgatgcactcaaacattgcagtatgttcaat

tgggagaactgggtttctaactactttgtatgcgatgggattatacAagggcaaggcgatacgtcggcatggggagtagc

cgatgctgtacagagaaacatctctacGaaatggctttatttctgtaatgatgatgtcattcgtcgccAcatgcagtgCA

GGCTtcatgatcacttggctttggtggagcacaagaagggcctgctacctgatgatattggcatggtgctttctgaggca

acatcctgcttcatcttaccagattataactactattancnattggcgacaatcccagattgcatattccaacnttcaca

cagcagcaagctgggtgtcttacatctacgtcgctgca

bPb-5027

TGCAGAAGGCAGTCGCGGAGGAGCTGGAGCTTCCGCCGTCGGTGATGGCCATCTTCGATCAGCAGGACGAGGAGGATGAT

TTCAGTGGGATAGACGAAGGCTCTCGAGGGGTGCTAGAAGATATCAGAACAGAAATCTTCATAAAGCTTGCTAGCAGTAG

ATTTGTGGTGATATTTCACAATGGGAGTGATGAATACATTGATCTGTATGAGTGTGGAGTCCCAGTAATATCATTTTTGA

GTATTAAGGTATTATGGACCTCGCACGGGAGGTTTGGACCCGGTTTTGATTACATGGAGAGCCACACTGATGTTATGGTC

AATTATTTTCATGATGATTGCATCCGCGATGAAGCAATATTGGAA

bPb-2058

tgcagcacagactccgtctgcgacccggcggtggcaccaacctgtaagacgacggacccccttacttgcattgcatttgc

ttcgtttatagatgaatttctcatgaccattgatgtctatgttctctgttttctgttgacatgacaaaaagcactatctg

ttctgatgatcttgtgatCgtcttgacgattttgcagatgcacaaggtgttgcggattgcatagctgagtgcaagaagag

gtgggggcaataattaacgccaagtaaccttaaaataaacagctggagaatctgtggatcatgggatcatatgatgctga

acgtctgtcaccgctgaaccaatcaataaaatgtgttccgcgtgggtcggtgccacttgtctgattttttttattttttt

tttgcaaatctcagctgcatatttatttcgtaagaagttaacacatactacaagaattaaatcattagcaatcccgtcca

agccgtccgtgaatcccagtgaaagcaagcaactcataagaacaaacacacaagatgggacggcatacgcataaatctga

tgcggaattctccccgacagatgcaacacagaaaacagtgtgacagtatgaacaaacacagcaagatcaaccgaaaagta

ctgattaatacaagcaagtcattatgtcttcaatcttcatgtgaattggCaaatttactacagcaaactcatccaagcct

tgCCtgattgttaatgaaacgaaactaattcgaaattcatcagtgagaaccagataaacattcaccaagaataaacgcag

aaaatgggggctaaatagaaatctggtgcagaatctaagaaaacaatgaaaagatgaacacatacataattaaggcaagt

aagatcaattgaaaggtactaactaatcgaagcaagcgaacaagccacttctgcatcttggccaccggaacgtaactcta

gatacctgca

bPb-9807

tgcagcacctcaatataagtgatcattcgagtattgtcctgtctccgattggaggcgagggccatgtcaaccagctggtt

ccagttgaagaactcagcatttctaaaaatggtgctaacggggaagtattgactcaactactctctcatttcccgaagct

caccatcctgcatatagctttttgcatgaaggtaggagaacttggtgtgatggagcagcaaagtggatcccagaagcagc

agacaaaagaagaggaggaaatagtggcgacggcagaaggaggggtgctgctcttgccaacccaactacagatcttgacc

ctccaacgttgccgtcgggtgagaatagtacccagttcagggggcgacaagaacgaatcagcaggaggtctccaacgtct

acgctccctccgcatagtgagcgcagcttactgccccgagttgctttcctcctattcgggctcctcgtctggtttccctt

tcccgacctcccttgaaagacttaaactataccaaactgaacaagtcaatctggccctcacaaacctatccaacctccaa

gaactcgagcttagctacatggagtccctcgaaaacctaagcatcaGccgttcctcagaactgaCcataaattactcctc

cggagttctgtcagagtccatctgca

bPb-7995

CATGGCGAAGCATTTCTCTTTCCAGCTATCGCCGGCGTCGCTGCCGCTGCACTGCCCTCCGTTGATGTTCCTGAACGTGG

CCCCGAAATCGCCGTCCAGCAGCTCCTCGAGGCGCCTCATGCGGATACGCAGTCTACGAAAGGGCGAATTCCAGCACACT

GGCGGCCGTTACTAGTGGATCCGAGCTCGGTACCAAGCTTGGCGTAATCATGGTCATAGCTGTTTCCTGTGTGAAATTGT

TATCCGCTCAA

bPb-4917

tgcagctcaaatacatgttcgtgttctttctgtttgatatcctgttacaacttcttgttcttgcttctatcgtggatatg

ttcagttcatatttactattttaaatgctatttgcataccctctattagattgcatgacttgtcttatctttGTtgtaat

agtcatgttttatctattatttttgttaattaaatcatatgatgaattgctgatatttccaAcaatatatgtacacacct

cgtcatttcattaggaagtgatcccagtctctagtaTtaaactattttttactacatatagccAtttaattatgttgtgc

tttttaaagaatctatacataacgtactactccctctgttTggaaatacttgtgggagcaatggatatatatttctgaac

aggaaatacttgtgggagcaatggatatatatttctggacagagggagtatatacgaagttaggaacacaactatctcta

tcccgaaagttaatttaatagaaattgtgtcttggttctttggatgctgatacaacaagaagcaaaatttggaaggagaa

acagtagagccttaactcttatataaaaaaggcttcttatacttcctctctAttGaataagcgaattggtcatgaatcaG

aataacacgcccaagaatgtacGagaaccaaataccaaaaagtaaatagggagaaagcataattagaaaatccatgtttg

tgaactcggtctcaaaagtaaaGactttaatgtgtttcatgtacagttactgca

bPb-4843

gctcggatccnctagtaacggccgccagtgtgctggaattcgcccttcagtcaagttagatggtgcagtaactgtacatg

aaacacattaaagtctttacttttgagaccgagttcacaaacatggattttttaattatgctttctccctatttactttt

atgtatttggttctcgtacattcttgggcgtgttattctgattcatgaccaattcgcttattcaatagagaggaagtata

agaagccttttttatataagagttaaggctctactgtttctccttccaaattttgcttcttgtggtatcagcatccaaag

aaccaagacacaatttctattaaattatctttcgggatagagatagttgtgttcctaacttcgtatatactccctctgtc

cagaaatatatatccattgctcccacaagtatttcctgtccagaaatatatatccattgctcccacaagtattcccaaac

agagggagtagtacgttatgtatagattatttaaaaagcacaacataattaaatggctatatgtagtaaaaaaatagttt

aatactagagactgggatcacttcctaatgaaatgacgaggtgtgtgtacatatattgttggaaataccaacaattcatc

atatgatttaattaacaaaaataatagataaaacatgactattataacaaagataagacaagtcatgcaatctaatagag

ggtatgcaaatagcatttaaaatagtaaatatgaactgaacatatccacgatagaagcaagaacaagaagttgtaccaGg

atatcaaacagaaagaacacgaacatgtatttgagctgcaccatctaacttgactgaagggcgaattctgcaGatatcca

tCACACtGgCGgCCgctcgagcatgcatctagagggcccaa

**bpb-14536**

bPb-8054

gctcgagcggccgccagtgtgatggatatctgcagaattcgcccttgatggatccantgcagttaacaacctgaattttc

tgattcagaatcccctttcctagatgagcatttctctgctgagcggatttgaggcgttggtgttcatttgacctataaac

tttgggatcaccttgtggttcttgatttctgtccatgtgccatgtcaatgtcaccagactgaccgaacctcgagcactcc

gctgatcctgatgtgtgctttactgaaacagagacccttatatcggatatctgattttgcttgtttgcttctgaaacagg

gagcgcttactttcagcattgttttacaaccatatgaagctgccctgacatttttcagttcgttttcttttgcaggctag

tgctttcttggatgttctaactgtgtgagttcgcatgagactgaagaccaaaaatgtctgctgca

**bpb-4405**

bPb-4565

tgcagcagcacaccaaatagacagcccgggccatgatgaatatataaacatcatagcgtgttaattaaatgtaaacaaca

agCaacatatatataagtgatcaacggagtatatagaactCtgagacttgattttctatctgtcccggcttCtacccatc

tcagtgaggtgtgagacaactagtatggaggcccccacgacagagtgaatagtggacatatatgtttgaagtaaaaaaaa

gatgatactccctccgccttaaaattattgtctttgatttgtctaaatataaatgtatctattcacaaaaaatggttgtg

agggcattgttgttgtcgaatgttgtggttcatccAagactttatcagcaatgatacctcgggctctatatagtgtatat

atgtgatacctcgggctctatatagTgtatatatgtttttttctacgtttggcctagactttttcaaagataatatagga

ttgacagtaaaaaaaagttcccaaagcaatactCcAAaaaagaaaagaaaaatccAacaataactgttgtagctagcAaG

aaaacttgggttccaaaaagggcccaaccgggttcgaaccggtgacctattgatctgcactggntcc

bPb-8283

tgcagtcgaacctcaggagtgtactttcagggacttctgttatcgacagcagatattgtgtgaattcacggtcacacaat

attttggattccgaatgatcaaattttctcgtgggggggcgagattctcgattaccacggttaccgagaaataccgaata

aatttcgtacgaatttataatggaattttgaattcaaattttgaaaactgtacaaaagacttgaacatgaatagatatga

aaagtgttagtagctcagtggtatgtcctgtacgtgctctctaattgctgtaagatcacgagtccgactctcatttgggt

agttttcttttttttgtaatataatggaattttgaattcaaattttgaaaactgtacaaaagacttgaacatgaatagat

atgaaaagtgttagtagctcagtggtatgtcctgtacgtgctctctaattgctgtaagatcacgagtccgactctcattt

gggtagttttcttttttttgtacgtcttgaaaaaaataaaaagttactgtcaaagtggtcgaacctcgaacctcttgaga

tcggtggtaaacggttaccaccaccccacagctgca

bPb-3068

ccagtgtgatggatatctgcagaattcgcccttgatgnanncagtgcagatgcaaggcctgtcttaattttaacagaaat

gatcctgtcttagttttaacagaaatgatccatgctgcttctagctagctagtagttgccgtgaccttttgatgttgcgt

cagagtgagtgagcaacttggactgtagttagtgacaacaactatggttttgcctctaatagctaacagacactctctac

cattacttagaaatgagtgagttcgggttgtaatcagtgacaccaacaattagggtatttgtctgtgttgtttttttaga

aacaagaggaaattcatttactaatatgcaaacaagagagtaaattcatttacccaagacccaagaggcacgactctttc

tgtgatatactgatattactagcagagattctcaagtttgatgcagaatttcttcaatgtagttatttttaccaaggttc

tgctcttcaattcctgca

bPb-9130

agtgtgatggatatctgcagaattcgcccttgacccagtgcagtatgtaggtgcagaactgattgcatgaatggaccgca

aagagataaaacatgccgcaaatttgttctgaagcacaagagtaaaacaaattgcacttcgaaattgcgttgttccGTtC

AacaaaaggaaaatatgcatggAAccatcaaaagaaactcaacaggtgccgccgcagtgaacagcaaaagcacttattta

tccaacatactcctacacaaccttggctcaaattgcgcacatctatcatcaaagatgcacaggaaagataacaacgaata

gtggttatattcatggaagaaattctgatatgaatcaagttttctgtatctatgcagatatgaatgaaagaattccagat

atcaatctgtttggaggtcgacaaaaattctgacaccaaagatctccatccaatctactctaagattatctgtctgttgt

atcatccgctggctctctttggcatggatgtgcctgagcttcactccagactccagagtctcgggtgatagtgtagctcc

tctgcgctacatcatggtgatgtctgtgaaatgaaacagcaaattaaatccaattagtgacaatatcacagccattaact

tgattaagattcgaaggatgcaggtacggataacaagaacaatcaaaggctgca

bPb-6607

tgcagccccgaggatacgggagctcgagctcctcctccggcagcccccctcatcCgagcagccaccagcgccgcCgcatc

caacctccaagGtCGgcccgtcggccctatcctTgccctgacctctctGctttccctcatcctcacctcgtttctctctc

tctctctcctagacacagaacgtcgtggtcgtgtcggcagcacgcctccgcccgagccagcagttgcatcctctctgtgc

atcgccctCGCtgttgttcgcctgcatcggtcgcgccagcgacctcagccgacggagcagcgaaacccttcttccctgcc

ctctctccttcttattccttccccaaaatttccctctCtgaagactctgtttgtcttgttttcttcagccaacagcagaa

tcgatgccaactcctgctgccttccatcggCgacgtcacgtgccgatctcgccggtgagaccccgctccctcatgtcccc

GtttctcccCttcttctctctctaaCctcacatgctctcCctatctttctgca

**bpb-4125**

bPb-6357

TGCAGGAGCGAACCAGCGCCGACCGAAGCGCTCGGTGGATGCGGACGGGGCGAGGTGAGCCATCGCGACCGCGGTAAGTG

CGGCGTGTTGGAGGCGACGATGCGATGACAACTTGAGGAGGAGCACCCGGAGGGCCGCGTAGCGGAGGCATGGAGTGGTG

CCAGCCAATGGTGCAGGTTCAGCGGCGGGCCAGCTTGGCGAGCGTGGCGAGCCAGTGGCGGCATAGGCGAGCTCGGAGAA

GCGTCGGCACAGGCGAGGCGCTGTGGAGGCGGAGTTGGCGGCTCGGACGCACGGCGTTGACTCCGTAATTGACCCGATAT

GTGGCCCCTTTTTATTTTTATTGCAGGAACTCTACCCCGGGACGGGAGTTGATATTTTGGGTGGATCATAATACTGCA

bPb-7786

tgcagttatctatctatatatctggttattaccacaaccagagacctttttttttatgaggcaaaccacaaccagagacc

ttgtacagtgcatgacagactttcactacaagaaggcttttattttatttaaattctcggcctataccgcggccaaactg

tgtctgtcctccaagtctccagccagcctctcccagccctcatggagctgcatcctcctcagcgacattgaactcgccgt

gatcggcaccccccggcaagagcagccatgctgccctcagtgacgttgaactcgccgtgatcggcacccccggcaggaac

agccatgctgccctcagtgacgttgaacttgccgtgatcggcaccctctgca

bPb-1933

tgcagtgggaatactagatgaaaccgtcaggccaaatccagcatggtctacaacgagcaggaaaccgtcgggccaaatcc

agcaaggtctacaacgagcaggccaccgaagataggtcacgggtctgctgaaggatgacgagaatcgcgccaagaaccgc

ggccgctgccttacacggaacgtcggtcctgagccgattgtgtaccgtctgaggtttgggaaatgcggatcccccagaag

gatttcggaagaaggttccttttgtcagccctgaacagccagccgatgtgggactcgcaggcagcacacagcgcaaccgt

ccacgtatatctgca

bPb-1176

tgcaggtgctattttatagctggaaacttattttcgtattcttcattctagtgaagaaatggatcaaaagatgttcatac

agggcaacaagcaagagaattaaatcgattagttcattaggattctgatgattttctcttttcggatgataacttgttaa

aacatatcacaattagctcagttagttcactgtgattatgataattttactaatttcaaatgaaaacttgctaaaacata

acacaacttttttttaaagagctttggaatctgtcatcgagcacacggatgacaatctttgagctcacagctcaaagaag

gacaagctaaacatgcattgattctatgtttctgttttcagacagcaagttattaagagattaataaaaagaagacataa

aaataatactgagtttgctgccactagtagtaccacaaacagtagcgaatggctgcgcctctgca

bPb-7446

tgcagctcaaatacatgttcgtgttctttctgtttgatatcctgttacaacttattgttcttgcttctatcgtggatatg

ttcagttcatatttactattttaaatgctatttgcataccctctattagattgcatgacttgtcttatctttgttgtaat

agtcatgttttatctattatttttgttaattaaatcatatgatgaattgctgatatttccaacaatatatgtacacacct

cgtcatttcattaggaagtgatcccagtctctagtattaaactattttttactacatatagccatttaattatgttgtgc

tttttaaagaatctatacataacgtactactccctctgtttggaaatacttgtgggagcaatggatatatatttctggac

aggaaatacttgtgggagcaatggatatatatttctggacagagggagtatatacgaagttaggaacacaactatctcta

tcccgaaagttaatttaatagaaattgtgtcttggttctttggatgctgatacaacaagaagcaaaatttggaaggagaa

acagtagagccttaactcttatataaaaaaagGcttcttatacttcctctctattgaataagcgaattggtcatgaatca

gaataacacgcccaagaatgtacgagaaccaaataccaaaaagtaaatagggagaaagcataattagaaaatccatgttt

gtgaactcggtctcaaaagtaaagactttaatgtgtttcatgtacagttactgca

bPb-7877

gtaaaacgacgnccagtgaattgtaatacgactcactatagngcgaattgnnccctctagatgcatgctcgagcgnccgc

cagtgtgannnatatctgcagaattcgcccttgannnatccagtgcagaggcgcagccattcgctactgtttgtggtact

actagTGgcagcaaactcagtattatttttatgtcttctttttattaatctcttaataacttgctgtctgaaaacagaaa

catagaatcaatgcatgtttagcttgtccttctttgagctgtgagctcaaagattgtcatccgtgtgctcgatgacagat

tccaaagctctttaaaaaaaagttgtgttatgttttagcaagttttcatttgaaattagtaaaattatcataatcacagt

gaactaactgagctaattgtgatatgttttaacaagttatcatccgaaaagagaaaatcatcagaatcctaatgaactaa

tcgatttaattctcttgcttgttgccctgtatgaacatcttttgatcCAtttcttcactagaatgaagaatacgaaaata

agtttcCAgctataaaatagcacctgcactggatcnatcaagggcgaattcnngcacactggcggccgttactagtggat

cnnagctcggtaccaagcttggcgtaatcatggtcatagctgtttcctgtgtgaaattgttatncgctcacaattncaca

caacatacgagccggaagcataaagtgtaaagnctggggtgcctaatgagtgagctaactcacattaattgcgttgcgct

cactgc

bPb-1758

tgcaggtgctattttatagctggaaacttattttcgtattcttcattctagtgaagaaatggatcaaaagatgttcatac

agggcaacaagcaagagaattaaatcgattagttcattaggattctgatgattttctcttttcggatgataacttgttaa

aacatatcacaattagctcagttagttcactgtgattatgataattttactaatttcaaatgaaaacttgctaaaacata

acacaacttttttttaaagagctttggaatctgtcatcgagcacacggatgacaatctttgagctcacagctcaaagaag

gacaagctaaacatgcattgattctatgtttctgttttcagacagcaagttattaagagattaataaaaagaagacataa

aaataatactgagtttgctgccactagtagtaccacaaacagtagcgaatggctgcgcctctgca

bPb-5885

tgcagctgctcctttcttatcttactttcacgaacaggaaacagtaccctctctgcaactctgtctggtccaaaacctcc

atgctccaaagaagctgcctgaccgggtccggctgcgaggccagactacctttctgcccggagccagcctcggtgcccgt

gccctccgactgtagcgtcccatcgaacgccgcgaaccacccgaccacaacaagggcagagctttctccaatcaatcagg

gtcaagttactactagcatataatagtacaatgcgaacatgcgcattcttcagttcaacacggacgaacaattcccttct

ctttgccagatttgattagttcctttttcatgatataatgatcgataaatgaagtcattgtgctccaactttgtaataag

atgttcattaaactgcactgctctccgtaaagagcagaaaccaaccactgttgtaaactgaaaagaacggacacctgcac

tttcttgtggctaggattggactttgaacttttcagttatggatcggagtgggtatgggcagctctcagaccattcccag

cttgaaatatatagtgctaagctcccctgca

bPb-5515

tgcagcaaagaagacaaaaggtttcaagccgtctccaatctcatgctgtacaaactcgtttatttttaccatagcgaatt

tagcgcttttgacttcgatgcttttcgctgctgctgctactactacttacgaactgcggattgcaacatgcagcaggttc

ttctccgaagccgctcagttctgtcgcgtcaatgagttggatatccgggccaagatacagaggaagatgctctcgcaaaa

cacaacgtatgtcgtatgcttggtgttcaagctagcatatgtatactctggtccgatttcacgcacgaggtggcatccgt

tggcgttgccgggagggagtcgacccggcaagtttgcgtgcaaggctacgtcgacgatgtggatggggctggcaatcctc

ccggagaggaagttcatttccctcgtgagaaagccgacggctggatggaggtggagttgggtgagttccataacgaggag

gatgacgatggtggcgaggtgtccatcagcttcacgggagaaagcaagtctggtcttaccgtgctgggcattgagctcag

aagtaagcaacaaaagccagcatgaatgtagcataaagtaacgctttgggggtaaaaatattattactatttgaaatgtc

tttggttacatgcatgcatgtgaagaggctaagaatttaagttacagtagtgattttttttaatataAggGGtTtccccC

ctgctCcAttttattgatgaagcaaagaccaCCagttttacaActctcacacaaacaaacaaggaccaacacagatcgaa

agtatctcaaaacaagaatttagccaagtcaccacctctaaacagaaaaacaagctaacttttaacagaaacatcagcac

atgtagcattcagctcctgca

**bpb-0649**

bPb-1029

tgcagctgtcagaagagggtagtagggccacgctgtcgaacacaatcaaagcatgcagtcaaaattatgttaaaaacaat

aaatactcttcatagaacatgacattctgataatgagcaccaaccaagttcagcattgtagccactgagtaccaagctgc

caatcgatgcagcaccaaaggcactgatccattgcttggtcgaatccccaaaccttaatgctgtggacttaacacctact

ttgaggtcatcttctttgtcctgacatgtattgggataccaccataatgaaaaacgataatcctgataagatgcttttcc

tcagtttagatatgaaagaacaccactagtaaaatacctgatgtgcatatatggtatcgtacaccaatgtccaacatata

ccggcagtatatagtggaaggatgactgca

bPb-8382

tgcaggtatatttatatccagatcagtttcattttgatcacaaaccatcttgtatagctctgcaagcaaacatatgtcag

ttgttaacgccaaaaacacgtgttgaaatataaggcatgagccaaaagtgtgctcttctaaatacgcagacatgcaccac

caaatgtgtacacacaactatgaaaaacaattgtccaataaataatcaaaaggcttaagtaaggTtgtgatggtcattct

agtggtaaaaaatggtaatatacgttgtcaggcaacttgtcttgttgatctcatttaactgtatcccaAcatcttatctt

ctaattaaataccagttgagaaaggtcatacatggaaaAcattgcctgttgaattttatgagacagatcatatagcacat

accatgccgattattcaggattacaatagcagaagcaccagcatcttcagcaAccttagcttttgtagtgaacttacaat

ctcctctttgcactagcaaaacttctccagcaacctaaataaaagaacaattaatcactacttcctctgtACCTaAaTAA

ATGTAGTTggggagaactaGTCtaGTTCTccccaActacaAttatttaggcacagatgGaATacataacaaTgtaggata

ctataaaaatttcgcaaacagctaaatcAtgttttcccagaaaggtaACcttttctttgagaggggtacaacaatcaaaa

gggtctgctaatagtagtcctgtccggttcgcgtgcttttcctttgactctattatggggccaaaccgagcaccaacacc

aacaaactcatcggtctctctgtttttgacccaagttcgcacctttacctgataaaaaggtgaggaaacaaaatttgaga

atgagtacatagtccattaaaaagaagtaagtagatgctgaagtacaaaatctccgccggagaaaaaaggaagagctgca

bPb-6875

TGCAGCTCTTCCTTTTTTCTCCGGCGGAGATTTTGTACTTCAGCATCTACTTACTTCTTTTTAATGGACTATGTACTCAT

TCTCAAATTTTGTTTCCTCACCTTTTTATCAGGTAAAGGTGCAAACTTGGGTCAAAAACAGAGAGACCGATGAGTTTGTT

GGTGTTGGTGCTCGGTTTGGCCCCATAATAGAGTCAAAGGAAAAGCACGCGAACCGGACAGGACTACTATTAGCAGACCC

TTTTGATTGTTGTACCCCTCTCAAAGAAAAGGTTACCTTTCTGGGAAAACATGATTTAGCTGTTTGCGAAATTTTTATAG

TATCCTACATTGTTATGTATTCCATCTGTGCCTAAATANNTGTAGTTGGGGAGAACTAGACTAGTTCTCCCCAACTACAT

TTATTTAGGTACAGAGGAAGTAGTGATTAATTGTTCTTTTATTTAGGTTGCTGGAGAAGTTTTGCTAGNGCAAAGAGGAG

ATTGTAAGTTCACTACANAAGCTAAGGTTGCTGAAGANGNNGGNGCTTCTGCTATTGTAATCCTG

bPb-7644

tgcagttttcagcatccacatgatcacaccgagaatcttcctgtcaacaagatacagtacactctggggtggttttagat

cgtgacaaggatgacaatgtcgatcactggatagtatcttgcaaattcagcaaaatctccgctggtaaaatgaggaggga

ggattttacatattaaccatctgatataatacagtactatgaggaccgggcgctttgctatgccctctgattaaaaaaga

ctaatcactcttttaaaatacaacatgtattactttgttaaaatggcaaaccttttaaatttaatcaaatttgtagataa

atatgtcaaaattaataatatcaaattctcgtgattagattcatcgttaaatgaattatcatacttttttatttaatatt

atggatgtcgatatttttactctcaacttggtcaaagttgaagaacgttgaaaagatgttgtagagctcaagctagctgc

a

bPb-0443

tgcagctagcttgagctctacaacatcttttcaacgttcttcaactttgaccaagttgagagtaaaaatatcgacatcca

taatattaaataaaaaagtatgataattcatttaacgatgaatctaatcacgagaatttgatattattaattttgacata

tttatctacaaatttgattaaatttaaaaggtttgccattttaacaaagtaatacatgttgtattttaaaagagtgatta

gtcttttttaatcagagggcatagcaaagcgcccggtccttatagtactgtattatatcagatggttaatatgtaaaatc

cttcctcctcattttaccagcggagattttgctgaatttgcaagatactatccagtgatcgacattgtcatccttgtcac

gatctaaaaccaccccagagtgtactgtatcttgttgacaggaagattctcggtgtgatcatgtggatgctgaaaactgc

a

bPb-6677

tgcagatcaataggtcaccggttcgaacccggtTGggCccTtttttaaacccaatttttcttgctagctacaacagttat

TgtTgGatttttcttttcttttttggagtattgttttgggaacttttttttactgtcaatcctatattatctttgaaaaa

gtctaggccaaacgtaAaaaaaaacatatatacactatataaagcccgaggtatcacatatatacactatatagagcccg

aggtatcattgctgataaagtcttggatgaaccacaacattcgacaacaacaatccggtttgaaagaccattttttgtga

ctcctccaaatgccgtcacaacaccatactgtcaatcctatattatctttgaaaaaagtctaggccaaacgtagaaaaaa

aatatatacactatataaagcccgaggtatcacatatatacactatatagagcccgaggtatcattgctgataaagtctt

ggatgaaccacaacattcgacaacaacaatccggttcgaaagaccattttttgtgactcgtccaaatgccgtcacaacac

cattttttgtgactcgtccaaatgccctcacaaccattttttgtgactagatacatttatatttagacaaatcaaagata

ataattttaaggtggaggaagtatcatcttttttttacttcaaacatatatgtccactattcactctgtcgtgggggcct

ccatactagttgtctcacacctcactgagatgggtagaagccgggacagatagaaaatcaagtctcagaGttctatatac

tccgttgatcacttatatatatgttgcttgttgtttacatttaattaacacgctatgatgtttatatattcatcatggcc

cgggctgtctatttggtgtgctgctgca

**bpb-2940**

bPb-2863

tgcagcagcacaccaaatagacagcccgggccatgatgaatatataaacatcatagcgtgttaattaaatgtaaacaaca

agcaacatatatataagtgatcaacggagtatatagaactctgagacttgattttctatctgccccggcttctacccatc

tcagtgaggtgtgagacaactagtatggaggcccccacgacagagtgaatagtggacatatatgtttgaagtaaaaaaaa

gatgatactccctccgccttaaaattattgtctttgatttgtctaaatataaatgtatctattcacaaaaaatggttgtg

agggcattgttgttgtcgaatgttgtggttcatccaagactttatcagcaatgatacctcgggctctatatagtgtatat

atgtgatacctcgggctctatatagtgtatatatgtttttttctacgtttggcctaGactttttcaaagataatatagga

ttgacagtaaaaaaaagttcccaaagcaatactccaaaaaagaaaagaaaaatccaacaataactgttgtagctagcaag

aaaacttgggttccaaaaagggcccaaccgggttcgaaccggtgacctattgatctgca

bPb-4626

tgcagtgtccttcttcagaggtggccaaacgggccagcccgacaaggcacgatacacgttaatcgtgcttggcatgaacc

gatacatgaggttacggttagtaaacgggttatgctataccgccccgtgtggtgcgcctgttagcctaggcacgagccct

tggctaatcggaccgccagtaggcacggctgggctgcatattctcggtatgctaagccttttagtgtgtgacagctgaaa

aatcctatctgacgccctcGgGcgttgaaaggatgtgGCttcgctgaagcgtggccacccgacgccacagtttgGgccgg

gccatggtcgctctgccttttcttcttttgttcaagatatattttgaaaagtatgttgaatgtgcatttaaatatcgttc

agcatatatgtgtacttaaatattgcgagttcgagtcggttcgcccgcctagcgctcttcggcgaatggcctaccttgcc

atttagccggtgccgtgtgttgagtgggtgctaacgggccagcccgccggtttggccttcttcttcctcctcctcctcct

tcttttggatctgctatatataaaaaaactgaggcataggagcagaggattgcagtgcacgtgccagcattcaatggacc

tcacacccacacccagacggtcacgggcacacacacatgatggaatggaaggaactgcgctacgtagatttggatttgga

tttggatttcaatttagatttggatatagggcgcgtacgaattgcattccttcattgtttgatttgatttgattaaGGCa

agcaagcaagcatctctcccgtcactacttactgtacttacttactactagtataatgttagaaacgagattaagcgtgg

tatggatggatgaaccatgaaccaacacttggcttggctgca

bPb-9817

tgcaggtacaagagattggaggtcctcgccgccttcaccaatgccgtaagcgtcgcaacctgccttttaacatcctttgt

ttgcatatatacaattttacggtggcaaatcatgtatgtacagaagcagcaataaccaagtagcagtgccgccaagtcca

ccttacatgtgtcaaacacaactgtcccctgttttctccacgataccgatgtcgaatagaaacaagacagtgcaagcacc

atgctatgataaaaacatatgcttcaattgcaaggcagcacaaaccaacacttgtttttAtcttcttatgtatgattttc

tatgtgccaaactaagctagcaacaacagctagattttcttgtagcacaaaactggaacagtctgttgatgcctttacat

tacataccaagtgaccttttggatcatttgtttaatttctttatacttggtggcacactttctggcctttctatcttttt

ttgtcctaacgttttgaccaccctccatatggttgtttatgtagacagctgttcctgctgttcctgtctttctccttggc

tgttgaagcgctgcattcatttatgcaggatgaatccgagcacaagtaagcgccatttatgtatgctcacagatggctct

gttcaaataattcttttctatgttggttgttgaatgtgtttaccttaGgaggaaacaaaggagatcggatgtgaaaaatc

cagtcactgtaatgcgttatgttgcatcgtactagtaTgataagtatgtgttggTtcagtccattgtgggtaactaaatt

gtcttctcaGaggaaactgcttaaaaaGaaaGaTAaTtgcatctacagtttgTagtcagtgTtacaccgcaattgatcat

ctgctgtccaacctctgtgacattattttcctgagaaaccgagaattgtttggtcaacttttatttccatgcatctactg

cctcttttcatcagccatggcctatcaatctaaactctttgctagcctaatattttccttgctcttaaaaaattatcttt

caggcattacctcattgtttctgca

bPb-9890

tgcaggtacaagagattggaggtcctcgccgccttcaccaatgccgtaagcgtcgcaacctgccttttaacatcctttgt

ttgcatatatacaattttacggtggcaaatcatgtaagtacagaagcagcaataaccaagtagcagtgccgccaagtcca

ccttacatgtgtcaaacacaactgtcccctgttttctccacgataccgatgtcgaatagaaacaagacagtgcaagcAcc

atgctatgataaaaacatatgcttcaattgcaaggcagcacaaaccaacacttgtttttatcttcttatgtatgattttc

tatgtgccaaactaagctagcaacaacagctagattttcttgtagcacagaactggaacagtctgttgatgcctttacat

tacataccaagtgaccttttggatcatttgtttaatttctttatacttggtggcacactttctggcctttctatcttttt

ttgtcctaacgttttgaccaccctccatatggttgtttatgtagacagctgttcctgctgttcctgtctttctccttggc

tgttgaagcgctgcattcatttatgcaggatgaatccgagcacaagtaagcgccatttatgtatgctcacagatggctct

gttcaaataattcttttctatgttggttgttgaatgtgtttaccttaGgaggaaacaaaggagatcggatgtgaaaaatc

cagtcactgtaatgcgttatgttgcatcgtactagtatgataagtatgtgttggTtcagtccattgtgggTaactaaatt

gtcttctcaGaggaaactgctTaaaaagaaagaTAattgcatctacagtttgTagtcagtgttacaccgcaattgatcat

ctgctgtccaacctctgtgacattattttcctgagaaaccgagaattgtttggtcaacttttatttccatgcatctactg

cctcttttcatcagccatggcctatcaatctaaactctttgctagcctaatattttccttgctcttaaaaaatatctttc

aggcattacctcattgtttctgca

bPb-2410

tgcaggtacaagagattggaggtcctcgccgccttcaccaatgccgtaagcgtcgcaacctgccttttaacatcctttgt

ttgcatatatacaattttacggtggcaaatcatgtatgtacagaagcagcaataaccaagtagcagtgccgccaagtcca

ccttacatgtgtcaaacacaactgtcccctgttttctccacgataccgatgtcgaatagaaAcaagacagtgcaagcacc

atgctatgataaaaacatatgcttcaattgcaaggcagcacaaaccaacacttgtttttatcttcttatgtatgattttc

tatgtgccaaactaagctagcaacaacagctagattttcttgtagcacaaaactggaacagtctgttgatgcctttacat

tacataccaagtgaccttttggatcatttgtttaatttctttatacttggtggcacactttctggcctttctatcttttt

ttgtcctaacgttttgaccaccctccatatggttgtttatgtagacagctgttcctgctgttcctgtctttctccttggc

tgttgaagcgctgcattcatttatgcaggatgaatccgagcacaagtaagcgccatttatgtatgctcacagatggctct

gttcaaataattcttttctatgttggttgTtgaatgtgtttaccttaggaggaaacaaaggagatcggatgtgaaaaatc

cagtcactgtaatgcgttatgttgcatcgtactagtatgataagtatgtgttggttcagtccattgtgggtaactaaatt

gtcttctcagaggaaactgcttaaaaggaaagataattgcatctacagtttgtagtcagtgttacaccgcaattgatcat

ctgctgtccaacctctgtgacattattttcctgagaaaccgagaattgtttggtcaacttttatttccatgcatctactg

cctcttttcatcagccatggcctatcaatctaaactctttgctagcctaatattttccttgctcttaaaaaattatcttt

caggcattacctcattgtttctgca

**-------------------------------------------------------------------------**

**Chr. 7**

bPb-0427

ggancaaaaatgtaatgtggagctcgaaccgggttgagaatacgaaatattcacacatactccgtgtgccgttcaagaca

gaagacggaaaggatttgcgccagtgggtgtcccggtttgacatttacccttacctagagagatacactcaggtttgcca

gctgcgaattattaAcaatcagtactcttgtgtgagtgatcttgagtaaaaaattatcccgatatgatttcaagatgctt

ctgccaagatccttgacattctagagggcaaaccagacttgatcattggcaactacactgacggaaacttggtggcgtcc

ctcatgtcaagcaagctaggagtcacacaggttaaaaaaatgtcttctaatcaaataaaagccctcaagatttcttttac

caccaatcacagttgacatgacatccctaagtggtcttcctgacagggaacaattgcacatgctctcgagaagacGaagt

atgagaactcaGatgctaagtggagagagctggaccaaaaataccacttctcctgccaattcactgcactggatccntcn

agggncnaattctgcananntccntcnnnnttgcnggcngctcgagcatgcatctagagggcccaattcgccctatagtg

agtc

bPb-0108

tgcaggtcggaatctatgaaagccttcatagcacggaggacgacgaagtggattggaagtggaggagagccgacgaggaa

actgagggagtctcccttgtttacacgttacacgttacacgtttgggaacactaattcacatgggcactttgatcatttg

caaatttacattacaatgatgtattgtgaatatgccgaagataggcgtgggtagggaaactgaatcagaaggacggccat

acatatggagcaaaatgaatgaatctacactctaaaatatgtctatatacatccgtgtgtagttcgtagtagaatctttt

aaaagacttatatttaggaacggagagagtagatagcaagctttatatatgtagctgacaagctcatgtagaattcagta

cggtacgaatagatatcaagctttatatgtatccagctggaagcacgcgtccgtccaactgtcacctcagcctcgcccgg

ctacctctattcctctctggcatagctgattgtaagcccctcctctgctccctctgatggtaaactctgaacttggttgc

ttgcttactttctgca

**bpb-9729**

bPb-8921

tgcagcagCTtcacaatatttcgatttggactggataaatgtctctttctcatcaaaaacaatggcgaaaggtttcaata

aaaacagttcttcgcgagaacggtcgtgctgcacttcaccatcggcggcaacctccatccttgcggaatcggcaaaggca

gaggaacacgaagcatgtatgccaccgtcagatctgtagagaccatgaatggcaatcagacttgtaaacgggaattctgt

ccacagccttgtactaacgccctaacaatcaagatcaaatagcttaactgacaaatcgactgagcctcatgtgtgccctg

ttttcctgcataaataaataaataaatactgaaatcgtttaactgacactgttacccaatctgaagccacgaaagaagtg

gatcagaatcggcggatgataacatcggatgtacctgca

bPb-2216

tgcagtggtgtttttcaaatctgctagaatcatcaatggctgtctttttcaaaaaaggacagtaacgatttctcctgcgc

tcgctaatccgcctatctgcgtgcgttgcatgcagcctgttaggagctcctaattagatgtatGCaaaaaaatgatgtca

gtggattctttttaaaaaTtGaaaAtTaaaaatgttttgtagcttaaactatccactcgattgaaaaactgttttcacat

aaaagattcttcatcacgagAccttcaaaattatatcgcgttttgataaatttcaacgacattttaaaaagattataaca

tctaaactacgtaacctatcacacctatAacacataaattatcatggtttttacagtgaggctattgggtatattttaac

cttttttgtctaagtcaaaatataccacatgattcgtaagtaaattattaggtccgcaattccaccgtccatcatcgatg

tcatgcctagccgcaaggtgaggttcacatgccgatgttgtatggcccgctcataaacatgactttgtgagtcgtgcttt

gctaggttcgtttaggcatgtgccgggttgtgattattgtacatgcattatctgca

**bpb-2875**

bPb-0043

tgcagtggtgtttttcaaatctgctagaatcatcaatggctgtctttttcaaaaaaggacagtaacgatttctcctgcgc

tcgctaatccgcctatctgcgtgcgttgcatgcagcctgttaggagctcctaattagatgtatgcaaaaaaatgatgtca

gtggattctttttaaaaattgaaaattaaaaatgttttgtagcttaaactatccactcgattgaaaaactgttttcacat

aaaagattcttcatcacgagaccttcaaaattatatcgcgttttgataaatttcaacgacattttaaaaagattataaca

tctaaactacgtaacctatcacacctataacacataaattatcatggtttttacagtgaggctattgggtatattttaac

cttttttgtctaagtcaaaatataccacatgattcgtaagtaaattattaggtccgcaattccaccgtccatcatcgatg

tcatgcctagccgcaaggtgaggttcacatgccgatgttgtatggcccgctcataaacatgactttgtgagtcgtgcttt

gctaggttcgtttaggcatgtgccgggttgtgattattgtacatgcattatctgcactgnnntccatcaagggcgaattc

cagcacactggcggccgttacta

bPb-5259

tgcagcatcttgatctctcatgtcatttttacagagaaggaatgtactcaagagacatttcatgggtagcgcgcattcct

aatttgcagtctcttggaatgaatgggGtaaacctcagcacggtagttgattggccttatgttaTcaatatgatcccttc

cttgaaggccctcagtctccagtcttgctctcttccaaccgcaaatcaatcactcccacatattaataaccttaccgaac

tggagaggcttgatctctctggcaacatctttgcccacccaatgtcaaggggttggttttggaatttgacaggcctccag

catctttaccttgccggcactctactgtacggtcaagcacctgatgcactggcacgtatgacgtcccttcaagtccttga

tttgtcaggtaatcgtgacatggggatgatgagtagtacaagcttaaagcacctatgcagtctgaaaattctggaccttt

ctttttgtcaaattgatggaaatataaaggatattatagggaggatgccccagtgtccattgaacagactgcAnnggatc

ca

**bpb-6976**

bPb-3418

TGCAGAAAGTAAGCAAGCAACCAAGTTCAGAGTTTACCATCAGAGGGAGCAGAGGAGGGGCTTACAATCAGCTATGCCAG

AGAGGAATAGAGGTAGCCGGGCGAGGCTGAGGTGACAGTTGGACGGACGCGTGCTTCCAGCTGGATACATATAAAGCTTG

ATATCTATTCGTACCGTACTGAATTCTACATGAGCTTGTCAGCTACATATATAAAGCTTGCTATCTACTCTCTCCGTTCC

TAAATATAAGTCTTTTAAAAGATTCTACTACGAACTACACACGGATGTATATAGACATATTTTAGAGTGTAGATTCATTC

ATTTTGCTCCATATGTATGGCCGTCCTTCTGATTCAGTTTCCCTACCCACGCCTATCTTCGGCATATTCACAATACATCA

TTGTAATGTAAATTTGCAAATGATCAAAGTGCCCATGTGAATTAGTGTTCCCAAACGTGTAACGTGTAACGTGTAAACAA

GGGAGACTCCCTCAGTTTCCTCGTCGGTTCTCCTCCACTTCCAATCCACTTCGTCGTCCTCCGTGCTATGAAGGCTTTCA

TAGATTCCGACCTGCA

**bpb-24334**

bPb-3461

cggataacaaattcacacaggnaacagnttatgnncntgattgcgccaagcttggtgcngaggttggatccACtagtaAC

GGccgccagtgtgctggaattcgcccttgagggatccagtgcagcatcttgatctctcatgtcatttttacagagaagga

atgtactcaagagacatttcatgggtagcgcgcattcctaatttgcagtctcttggaatgaatggggtaaacctcagcac

ggtagttgattggccttatgttatcaatatgatcccttccttgaaggccctcagtctccagtcttgctctcttccaaccg

caaatcaatcactcccacatattaataaccttaccgaactggagaggcttgatctctctggcaacatctttgcccaccca

atgtcaaggggttggttttggaatttgacaggcctccagcatctttaccttgccggcactctactgtacggtcaagcacc

tgatgcactggcacgtatgacgtcccttcaagtccttgatttgtcaggtaatcgtgacatggggatgatgagtagtacaa

gcttaaagcacctatgcagtctgaaaattctggacctctctttttgtcaaattgatggaaatataaaggatattataggg

aggatgccccagtgtccattgaacagactgca

bPb-4808

TGCTCGAGCGGCCGCCAGTGTGATGGATATCTGCAGAATTCGCCCTTGTAGACTGCGTATCCGGATCTCGGCTTGCTTGG

AAGGTTGCCGGCCCTTCATACTCTGCTGTTAAAGAGCCTCCGCCATG

bPb-1285

CATGTATAATCAGCCCTCCTTCAAGCTGTGTCAATGCCCCAAGTTCTGTCAGTTCAAATCCCATGAGGGAGAGGACAGAA

AGGGAGGAGGGAGGACGCACGGGGGACTGATCCGGATACGCAGTCTACGAAGGGCGAATTCCAGCACACTGGCGGCAATT

ACTAGTGGATCCGAGCTCGGTACCAAGCTTGGCGTAATCATGGTCATAGCTGTTTCCTGTGTGAAATTGTTATCCGCTCA

ANNT

bPb-4552

tgcagcatactcaacaaaatgtattacttattcagtcttcacaggccggctttataccaaaagataccactacttgtagc

atctgtatgtgcatggtgtggcaatgtcttttgctcaatgtcttcggtcacatctatggtgttgcaatctttttgctaaa

caacatgcctatgcttgtgctttcccagctgtcgtgttgctacaaatttttgaaggacgactacaagaaactacactcaa

ttgatgcaaccgctctttatagaattgcgctatcatcggatcccgatgcggttgttagcgtcagtggcagtgattccgtg

tgttcagacggcccagcaaacaacagccatagccaaatgacgtcaagcgaaaacgaaagcatatgcaggatcggtaggtc

ttactcctagtgctgcttaggagtaaaaagttcagtacacatgtttcattttcccatagcacacatacaattactatcat

gtgtttttaacctgctgctgcccagtagtccaatgcaaattgctggtcttagccttcacaactcaaagtcaactaagtaa

ccgctcaaacagttaaggtgctgcttctttttgggaagcattgaagcacaactcttaacagaaatgttagcaacggtctg

ca

bPb-6109

tGCagaccgttgctaAcatttctgttaagagttgtgcttcaatgcttcccaaaaagaagcagcaccttaactgtttgagc

ggttacttagttgactttgagttgtgaaggctaagaccagcaatttgcattggactactgggcagcaGcaggttaaaaac

acatgatagtaattgtatgtgtgctatgggaaaatgaaacatgtgtactgaactttttactcctaagcagcactaggagt

aagacctaccgatcctgcatatgctttcgttttcgcttgacgtcatttggctatggctgttgtttgctgggccgtctgaa

cacacggaatcactgccactgacgctaacaaccgcatcgggatccgatgatagcgcaattctataaaGagcggttgcatc

aattgagtgtagtttcttgtagtcgtccttcaaaaatttgtagcaacacgacagctgggaaagcacaagcataggcatgt

tgtttagcaaaaagattgcaacaccatagatgtgaccaaagacattgagcaaaagacattgccacaccatgcacatacag

atgctacaagtagtggtatcttttggtataaagccggcctgtgaagactgaataagtaatacattttgttgagtatgctg

ca

bPb-8103

TGCAGACCGTTGCTAACATTTCTGTTAAGAGTTGTGCTTCAATGCTTCCCAAAAAGAAGCAGCACCTTAACTGTTTGAGC

GGTTACTTAGTTGACTTTGAGTTGTGAAGGCTAAGACCAGCAATTTGCATTGGACTACTGGGCAGCAGCAGGTTAAAAAC

ACATGATAGTAATTGTATGTGTGCTATGGGAAAATGAAACATGTGTACTGAACTTTTTACTCCTAAGCAGCACTAGGAGT

AAGACCTACCGATCCTGCATATGCTTTCGTTTTCGCTTGACGTCATTTGGCTATGGCTGNTGNTTGNTGGGCCGNCTGAA

CACACGGAATCACTGCCACTGACG

**bpb-49307**

**bpb-2678**

bPb-8558

tgcagatgataatcagctcaaattggattcctccctttaaattgaagttggcctatttcccttctagcaaacttggacct

cagtttcctttgtggcttaaagggcagggaaatatcagttatcttgacatttctaatgcagacatagttgaccaactccc

agattggttttgggatgtgttttcaaatattcagtatctgaacatctcttgtaatcaaatcagtggctggttaccgagta

cgttggaattcatgtcttcagatgcgggtatcatatttgacctcagcttcaacaacgtcactggtgttttacctcagtta

ccaaggcatttggtcgaacttgacatttccagaaactcattatcagggccactaccacaaaattttggagctccattcct

tggtgatttgttgctttcagagaacagtatcaatggcactactcccatacatatctgtgagctgca

bPb-0714

tgcagatgataatcagctcaaattggattcctccCTTtaaattgaagttggcctatttcccttctagcaaacttggacct

cagtttcctttgtggcttaaagggcagggaaatatcagttatcttgacatttCtaatgcagacatagttgaccaactccc

agattggttttgggatgtgttttcaaatattcagtatctgaacatctcttgtaatcaaatcagtggctggttaccgagta

cgttggaattcatgtcttcagatgcgggtatcatatttgacctcagcttcaacaacgtcactggtgttttacctcagtta

ccaaggcatttggtcgaacttgacatttccagaaactcattatcagggccactaccacaaaattttggagctccattcct

tggtgatttgttgctttcaGagaacaGtatcaatggcactattcccatacatatctgtgagctgca

bPb-1645

ccagtgcagctgtggcacgcacggtgtgtaccacagagcaaaaggattgcggagcaaagtgactcactcgagtgacaact

tctatgcttctactccctcagtttttaaatataagaccttttagagattttattatagattagatacaaagtaaaatgag

tgaacctacatcttaaaataaatgtatgtagttcatagtaaaaatctctaaaaggtcttatacttcatttaggaacggat

gtagtatatattaacctaggagaagcatgccaaggatgaagatacgggccaccagtcgaccgcatgccatactatgttac

ctacaagactcaacccttacacatacagatgaactgacacggtcctgcactaaacaaagggcagtgatgactaagcaaat

atcgttgaaagtctgagataaatttaaaaataatgcaagcatcaatgttaaatttaggacttgattaaatttaggacttg

agccctgacgagatgaggatatcactattcttctaaccatccaaccacattgctgcttcgtagaagttgtattcgattta

atgcttggaggggctcacttctgctgccggtgcgggcaagaatgccacgaggagcaacagcagcgccaggatggcggcga

ggaggggggaggccgccatctcttcccgtcactccctgcactggatccntcnaggggcgaattctgcagatatccntcnn

nnntggcngnnnctcgagcatgcatctagagggcccaattcgccctatagtgagtcgtatta

bPb-2191

TGCAGATCTTCTCCGTCGAGGTCACGGAGCTCAGCGGGGGCCTCCAGTGGCCGCTCGAGGTCTACGGCGTGGTCGCCACC

CGAGACTCGGTGGACCATAATCGCAACGTCATCTTCAGGCGCAGGAGGGACGCCTCCCTCGTCCTCACCCAACAGGTTTG

TTTTTCCTCTCAACCATCATCTGTAGACTGTAGCTGACATTGGTGTGGTATTCTGTTTTCCGGATGCAATCGGTTGAAGT

TCATCTGTTAACGTGTGCAACTGCTGACTGCA

bPb-4312

tgcagcatgtcagaacaatgtatagcctgaggcagaggtagtgatctgtgttcatgggaagaaatccgtagtatgggact

ggggagtggggaaacgtggttggttacacaggcttctggccctagcgcaataatcatctgggtcgttaggattagtttac

cttataatactagggagctgtttggcagccctccacggagcggagcgcgctgttaattaacagctccgcaaaatattgct

ggatgcgcgcgcgactccgctccaacgcgcagtaaattgacagaaaccgcactaacaatatgccatttgtgtgttgcagg

ccccgctatacgtggacacgaagatggttatgagcatggtaacgtggggaggctctgtgtcacaggtgatcatgccaaca

ccagacgcgtacgccagtgcgggggcaagctggatcgggcactgccgcccggtctgcctgcccaacttgggccaccgact

gca

bPb-7382

tgcagagatggaacacggataatttngggtgatgaaactaantgcggagattaatacacctatgatggcatacacgattt

acactctttgaattctcttaggtgcccaaaacactcggacggtgggagttgctagttacaaaaaaaatgatgtatttatt

aatattgatcgatctattttggaataacgcaaattcagtttagtacaaaaaatgcaacttgcacgtttaagaaagaattg

tagctgcatgtagtgtgcatattagaagaaaaaattcaatcaactggtggacctaACaataaatgcatattgcacgtatg

ccctttgtggttaaacagtgcaagcattgatgctagccgctccatgcccacgcccacttctcacacaatcctttgcttct

tcgttataaatatgtgttcagcaatctaagcacgaaatataaggctctcaattttctcttattgattcatctacAatttt

tcaagccgttccggccaccaccgccgccatggtaccctcttaatttttctgtgcgtagctacAgtaaccacactcctata

cctcaTgagccgttcacgcacggtataccttgttgattatgcgtgctttcggtctcactcgaactaccgcatcagcaagg

ctgaatggattgagaacattcaccactcccggtcgtacgacGaCAgcggcaagcatcgcttcctgacccgtatttctgaa

cggtcaggccttggtgatgaGacctacctcccaccttaccatcatcatatcccGccgtattattatttaagtgaagcccg

tgctgaggctgagttgtctatcttcacgaccatagatgatttgctcctgaagacgtgtatcgaccttgatgcaatcgcca

tactcgtcgtgaactgca

bPb-8660

tgcagtcggtggcccaagttgggcaggcagaccgggcggcagtgcccgatccagcttgcccccgcactggcgtacgcgtc

tggtgttggcatgatcacctgtgacacagagcctccccacgttaccatgctcataaccatcttcgtgtccacgtatagcg

gggcctgcaacacacaaatggcatattgttagtgcggtttctgtcaatttactgcgcgttggagcggagtcgcgcgcgca

tccagcaatattttgcggagctgttaattaacagcgcgctccgctccgtggagggctgccaaacagctccctagtattat

aaggtaaactaatcctaacgacccagatgattattgcgctagggccagaagcctgtgtaaccaaccacgtttccccactc

cccagtcccatactacggatttcttcccatgaacacagatcactacctctgcctcaggctatacattgttctgacatgct

gca

bPb-6747

tgcagttcacgacgagtatggcgattgcatcaaggtcgatacacgtcttcaggagcaaatcatctatggtcgtgaagata

gacaactcagcctcagcacgggcttcacttaaataataatacggcgggatatgatgatggtaaggtgggaggtaggtctc

atcaccaaggcctgaccgttcagaaatacgggtcaggaagcgatgcttgccgctgtcgtcgtacgaccgggagtggtgaa

tgttctcaatccattcagccttgctgatgcggtagttcgagtgaggccgaaagcacgcataatcaacaaggtatacagtg

cgtgAacggctcatgaggtataggagtgtggTtactgtagctacgcacagAaaaattaagagggtaCcatggcggcggtg

gtggccgGAACGgCTtAaaaaaTTgtagatgaatcaataagaGAaaATTgagagCCTtataTtTcgtgcttagattgctg

aacacatattataacgaaGaagcaaaggattgtgtgagaAgtgggcgtgggcatggagcggctagcatcAatgcttgcac

tgtttaAccacaaagGgcatacgtgcaatatgcATtAttgttagntccaccagttgattgaattttttcttctaatatgc

acattacatgcagctacaattctntcttaaacgtgcaagttgcannntttgtactaaactgaannngcgttattccaaaa

tagatcgatcaatattaataaatacatcattttttttgtaactagcaactcccaccgtccgagtgttttgggcacctaag

agaattcaaagagtgtaaatcgtgtatgccatcataggtgtattaatctccgcaattagtttcatcacccaaaattatcc

gtgttccatctctgcactgga

bPb-2533

tgcaggactatgcctaggcctatcaagaaacaaactggcctgattatttcaccatgaagaagcaacatggcaccgcgaaa

ctgcccagtagaggaccatgaaacatgtggaaaacaaaggataacttttttgcaatggacatctaagagaactctaaaaa

tgattcttaccattggcccagcaacttcatcagccttttctttgccatacagttgttcaaccaaaacaatagaatactcc

atggttgttcctgtcccacggctagtcacacaattcccatcaatctgtaaccttgattctacagcttgcacctctgaagg

aagtttgttcatgcacggcggatgacaggttgcctatcacaaacactcaaatgagctgataatacaaagcatacagaaaa

tgaataatgggtaaaattgtaacctttaatacattgagcaaaccccaagctcccagtgccacagctggtgcagcacatac

tgca

bPb-5852

GGGCCCTCTAGATGCATGCTCGAGCGGCCGCCANTGTGATGGATATCTGCAGAATTCNCCCTTGATGGATCCNNNNNNNN

NNNNNNNNNNNNNNNNNNNNNNNNNNNNNNNNNNNNNNNNNNNTGCAGTATGTGCTGCACCAGCTGTGGCACTGGGAGCT

TGGGGTTTGCTCAATGTATTAAAGGTTACAATTTTACCCATTATTCATTTTCTGTATGCTTTGTATTATCAGCTCATTTG

AGTGTTTGTGATAGGCAACCTGTCATCCGCCGTGCATGAACAAACTTCCTTCAGAGGTGCAAGCTGTAGAATCAAGGTTA

CAGATTGATGGGAATTGTGTGACTAGCCGTGGGACAGGAACAACCATGGAGTATTCTATTGTTTTGGTTGAACAACTGTA

TGGCAAAGAAAAGGCTGATGAAGTTGCTGGGCCAATGGTAAGAATCATTTTTAGAGTTCTCTTAGATGTCCATTGCAAAA

AAGTTATCCTTTGTTTTCCACATGTTTCATGGTCCTCTACTGGGCAGTTTCGCGGTGCCATGTTGCTTCTTCATGGTGAA

ATAATCAGGCCAGTTTGTTTCTTGATAGGCCTAGGCATAGTCCTGCA

bPb-2373

tgcagcttatcttcttagagggagataggccagagtcggggagaagtgagacacgcgaggccttccgcgtcgttccgctg

atcccggggtacgagcgcattctggatctaagaggaatatgccattctagggaaggagtgggttctggttcctcgacgag

acgaaacacgagaatgacccttaggaacggaaagtacccgttatattccatccgatgatctgaaccaaaaagatcaaaat

gatcatccgttgagaaaaggatcaccctaagatgatcatctcatggctattggaaatgaatcaaattagatggttctatt

tctcaacctttctgacctgctcctatggaaccaaggtcgaaaggattggaaaagtcattcattcacaaggactcatgaag

ggttcctcaaaaaaggtaaggattagtagttcttttttgacatcgatttcagaaatgaatggattcggtcctgtcataca

taccgaaatctcccacttgaagactatgccgctgaaaagccatgtatttagttgtaatgaatggagaagctagctaggta

gctaccagctaattatattgttaaaccatgtgccctcatatgacattaggcaactgttgggagaggagctatctggtacc

actgttcaagatctgca

bPb-9181

CTCGGANCCNCTAGTAACGGCCGCCAGTGTGCTGGAATTCGCCCTTTCGTAGACTGCGTATCCGGATCCAGTGCAGCTAT

GGCAGTTTAGGCAGCTACCGCTTTGGTTGGGTTCGGGTCCCTGTCTCACATG

bPb-4646

tgcagctatggcagtttaggcagctaccgctttggttgggttcgggtccctgtctcacatgttggtgtatggccgatgag

acgtttccatctgggaaggtccaagtgagactctggcatcctttcagtgaatgaaaaacagtaaaaagacccaataagaa

acactgagcatctcggcaaaggaacttaaatattgccaaactgatctcacaaagtaaccgaaagttaattgtagtgcaag

tctgtaactgtacaatatgtcaatatccagcttcttgaactgtccggacgtatagttttgcacaatatttcatttgcgtt

gatgcaatttttcagccccttggttaaaacttaaaaggcagaatagacacacaagttcaactgca

bPb-4541

tgcagttaaatgtgaacccccaccgGccatctgaagataattatttccatttgccggataagggacatttgctgaatagt

ttggcatagccatgggtgggacataaactggagaatacagaggatgtcgatatggcatgaaattcggataatgctgcaca

tgcattggagggtacatctgtgctgcttgttgttgctgggacatatgttgctgttgtgtagacattgcaatagggctgct

gctcgcaagctgtgaacttagagcctgcaaaaacacaacaggtgaaagataccatgagataacagtacttgcgatcaaga

gatgagttgctgggcttgcttggcacatgataggggagaaagtcagtgggttagttattttttcttccctccgtgaagca

cagccAccaaacatatactccctccgttcacaaatataagatgttctaacttttttctgaatcggacatatatagacaca

ttttagtatgttcgttcactcatttcagtccatacgtaaccatattgaaatatccaaaacatcttatatttctgaacgga

gggagtacctatcccgcccccaatgttccagatctgctggggccacttcactattcctgca

**Supplemental Figure 1.** The sequences of the relevant DArT makers that are associated with the QTLs in the present study.
